# Replicators, genes, and the C-value enigma: High-quality genome assembly of barley provides direct evidence that self-replicating DNA forms ‘cooperative’ associations with genes in arms races

**DOI:** 10.1101/2023.10.01.560391

**Authors:** M. Timothy Rabanus-Wallace, Thomas Wicker, Nils Stein

**Affiliations:** University of Melbourne, Melbourne, Victoria, Australia; Leibniz Institute of Plant Genetics and Crop Plant Research (IPK), Seeland, Germany; University of Zurich, Zurich, Switzerland; Center for Integrated Breeding Research (CiBreed), Georg-August-University Göttingen, Göttingen, Germany

## Abstract

The C-value enigma—the apparent disjunction between the complexity of organisms and the sizes of their genomes—could be in part resolved if it were definitively shown that tolerance of self-copying DNA elements incurred an occasional selective advantage. We leverage the power of the latest genome assembly of the exceptionally repetitive and well-studied cereal crop barley (*Hordeum vulgare* L.) to explicitly test the hypothesis that the population of genes that have been repeatedly replicated by the action of replication-inducing sequences has undergone selection, favouring genes involved in co-evolutionary arms races (such as genes implicated in pathogen resistance). This was achieved by algorithmically identifying 1,999 genomic stretches that are locally rich in long repeated units. In these loci, we identified 554 geanes, belonging to 42 gene families. These gene families strongly overlap with a test set of pathogen resistance and other likely evolutionary ‘arms-race’ genes compiled independently from the literature. By statistically demonstrating that selection has systematically influenced the composition of replicator-associated genes at a genome-wide scale we provide evidence that tolerance of repeat-inducing DNA sequences is an adaptive strategy that may contribute to enigmatically inflated C-values, and invite more detailed research on how particular genes become prone to duplication, to the organism’s advantage. To this end, we examined the genomic sequences surrounding several of the candidate gene families, and find a repeated pattern of genomic disperse-and-expand dynamics, but where the repeated genomic unit itself varies between sites of expansion. This suggests that genes effectively form opportunistic relationships with replication-inducing DNA elements. We mention implications for agriculture.

## Background

*―Thus, some selfish DNA may acquire a useful function and confer a selective advantage on the organism. Using the analogy of parasitism, slightly harmful infestation may ultimately be transformed into a symbiosis*.

—Orgel and Crick, 1980^1^

The C-value enigma describes the counterintuitive observation that the sizes of organism’s genomes seem to bear little correspondence to the organism’s complexity and the number of genes present in the organism’s genome. The human genome of 3.5 Gbp is dwarfed by the Axolotl (*Ambystoma mexicanum;* 32 Gbp) and the Australian Lungfish (*Neoceratodus forsteri*; 43 Gbp), and even these values are middling compared with plants, the longest known genome of which belongs to the octoploid Japanese canopy plant *Paris japonica* at 149 Gbp. What is perhaps more striking is the variation withing groups with, for instance, diploid grass genomes vary in size by three orders of magnitude, with C-values ranging from 0.3 (~294 Mbp; *Oropetium thomaeum*) to 13.80 (~13.51 Gbp; *Lygeum spartum*)^2^.

For several decades it has been understood that genome size differences are primarily driven by the presence or absence of self-replicating DNA elements. The replication of long (kilobase-to-megabase-scale) genomic sequences occurs via several mechanisms including unequal crossing over (UECO), retro- or DNA transposition, strand slippage during replication, and even whole-genome duplication. A DNA element that promotes these phenomena can in theory continually increase its frequency in the genome over generations, even while having neutral or slightly negative fitness consequences for the organism^3^. Despite dominating the literature of the 1980s and 1990s, the ‘selfish replicator’ interpretation raises the critical question of why genes have not arisen that completely suppress this ostensibly wasteful replication, especially given evidence that the activity of some self-replicating elements is amenable to genetic control^4^. Evolutionary biologists since the 1970s have explored the idea that self-replicating elements might confer occasional selective benefits. In particular, duplication of genes involved in evolutionary arms races might confer an advantage at the lineage level by allowing the mutational landscape to be explored more comprehensively. If these occasional advantages outweigh their negative impacts on average, then lineage-level selection can favour replicator tolerance (fig. 1). For this hypothesis to be viable, it must be systematically shown that selection influences the association of arms-race-related genes with replication-inducing elements.

**Figure 1.**
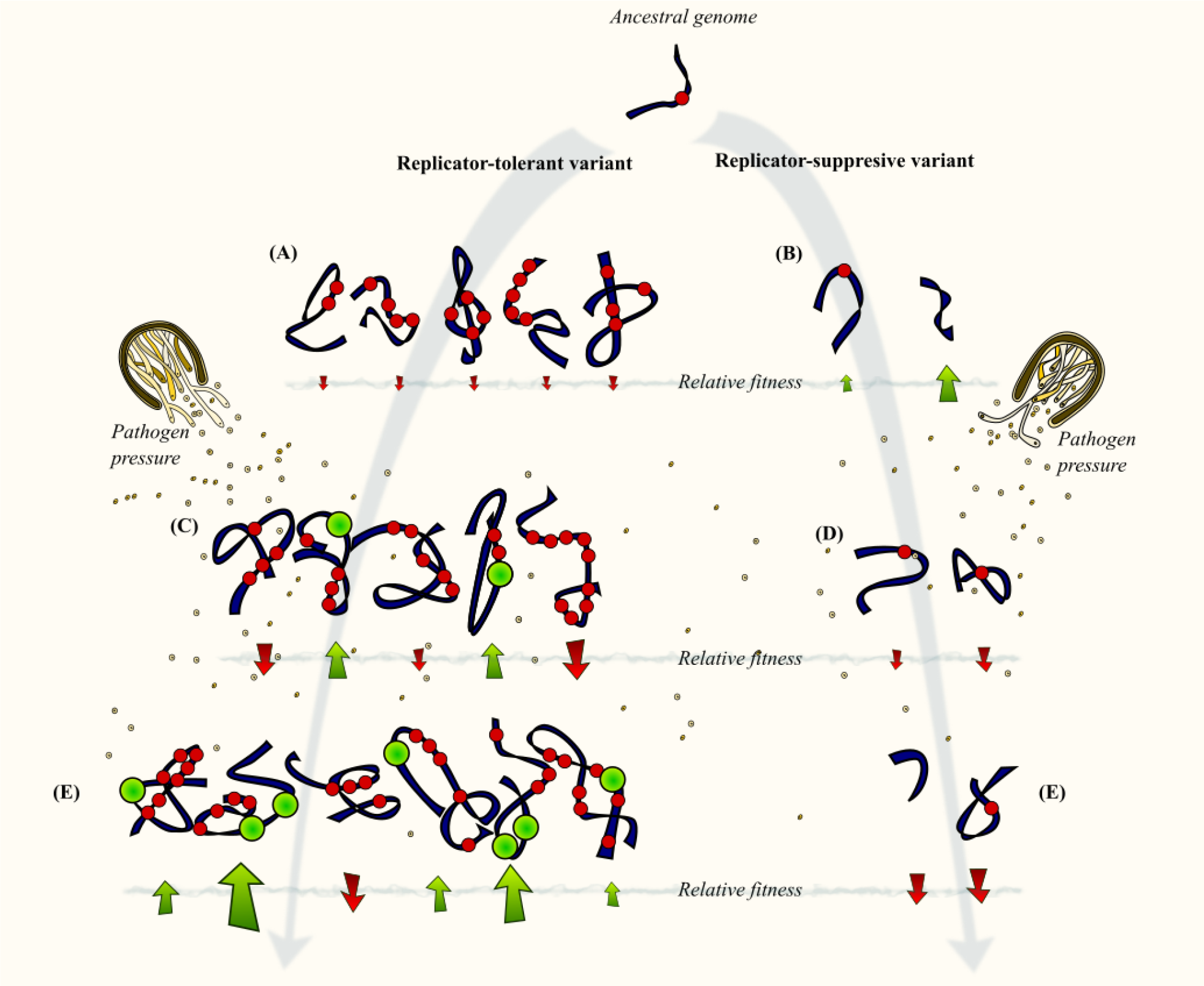
The Replicator-Gene Symbiosis hypothesis, a conceptual diagram. Navy strips represent the genomes of individuals in populations over time (grey arrows). Overlain circles represent repeated DNA with slightly negative (red) or positive (green) fitness consequences. The pathogenic fungus depicted is stem rust (*Puccinia graminis* Pers.) **A**. In the replicator-tolerant variant, slightly deleterious self-replicating DNAs including l-DPRs accumulate. The genome expands. Many gene variants exist in the population. **B**. In the replicator-tolerant variant, there are fewer self-replicating DNA elements. The genome size remains similar. Gene diversity in the population is comparatively limited. **C**. In the replicator-tolerant variant, some replicated genes confer pathogen resistance, conferring a fitness boost that outweights the detrimental effects of replicator proliferation. **D**. In the replicator-tolerant variant, fewer opportunities to develop pathogen resistant strains occur. **E**. Over generations of coevolution with pathogens, replicator-tolerant lineages are favoured, and the genome expands until the detrimental effects outweigh the arms race advantage.

Research and scientific discussion relevant to this hypothesis has come from at least four broad research areas. *Firstly*, the literature on gene duplication: As early as 1970, Susumu Ohno’s book *Evolution by Gene Duplication*^5^ discussed the possible selective benefits conferred by increasing the dosage of certain gene products, by the establishment of permanent effective heterozygote advantage, or by creating beneficial neofunctionalisation opportunities. Early literature on the phenomenon was firmly grounded in multi-level selection theory, depicting self-replicating elements in the genome as a ‘community’, analogous to a community of organisms in a biome, and with similar dynamics such as competition, population demography, migration, adaptation etc^6,7^. Other authors have emphasised that genetic redundancy may be selectively favourable for its ability to provide a buffer against undesirable future mutations^8^, and yet others note a beneficial regulatory role for self-copying DNA. Following her discovery of ‘jumping genes’, Barbara McClintock suggested the activation of transposable elements under stress may be adaptive, serving to bring about gross changes to gene regulatory networks. Studies producing empirical evidence for these phenomena continue to accumulate^9^. *Secondly*, important evidence comes from functional genetic research into the influence of self-replicating elements on particular genes: Since the genomic era studies routinely present evidence for specific cases in which the elements responsible for C-value inflation are shown to confer a selective advantage, often by altering gene regulation, in some cases even describing in detail the molecular mechanisms by which this occurs^10–16^. *Thirdly*, there is the literature on specific gene families, primarily those involved in pathogen recognition and response: These have frequently noted the tendency of genes involved in pathogen recognition and response to come in diverse families often resulting from multiple localised replication events^17–19^. And *fourthly*, there is research focusing on the self-replicating content of the genome as a whole^7,16,20,21^: This field has largely embraced and supported the ‘community’ view of the self-replicator landscape, especially given the ability to date LTR-retrotransposon transposition events, making inference of historical transposon population expansions and contractions possible.

All four of these directions provide some circumstantial evidence in favour of the hypothesis that replicator-gene associations might be under selection, and hence that beneficial replicator-gene association is a valid explanation for replicator-tolerance, which contributes to the enigma of C-value variability. Following the historical literature on the topic, we embrace the multilevel-selection-based interpretation that a beneficial replicator-gene association is most usefully described as a symbiosis between gene and replication-inducing element, referred to here as replicator-gene symbiosis (RGS), instances of which are called replicator-gene symbiotic units (RGSUs).

In the present study, we aim to tie these directions together by explicitly testing the RGS hypothesis using a novel analysis approach (fig. 2). Using the annotated barley genome for practical reasons, we used a gene-agnostic technique to identify regions heavily affected by replication inducers, based on the presence of long, locally-repeated sequence stretches. We call these long, duplication prone regions (l-DPRs). We then identified gene families with a statistical tendency to comprise members that occur in l-DPRs in more than one copy per l-DPR. These represent candidate RGSUs. Independently, we compiled a list of gene families likely to be involved in evolutionary arms races, in particular, pathogen/predator resistance and inter-organism interactions. We were then able to statistically test whether selection based on function had influenced the candidate RGSU pool, by testing for a greater-than-chance overlap between candidate RGSU genes, and candidate arms-race genes. Following this we examined several candidate RGSUs in detail to better understand their dynamics in the genome.

**Figure 2.**
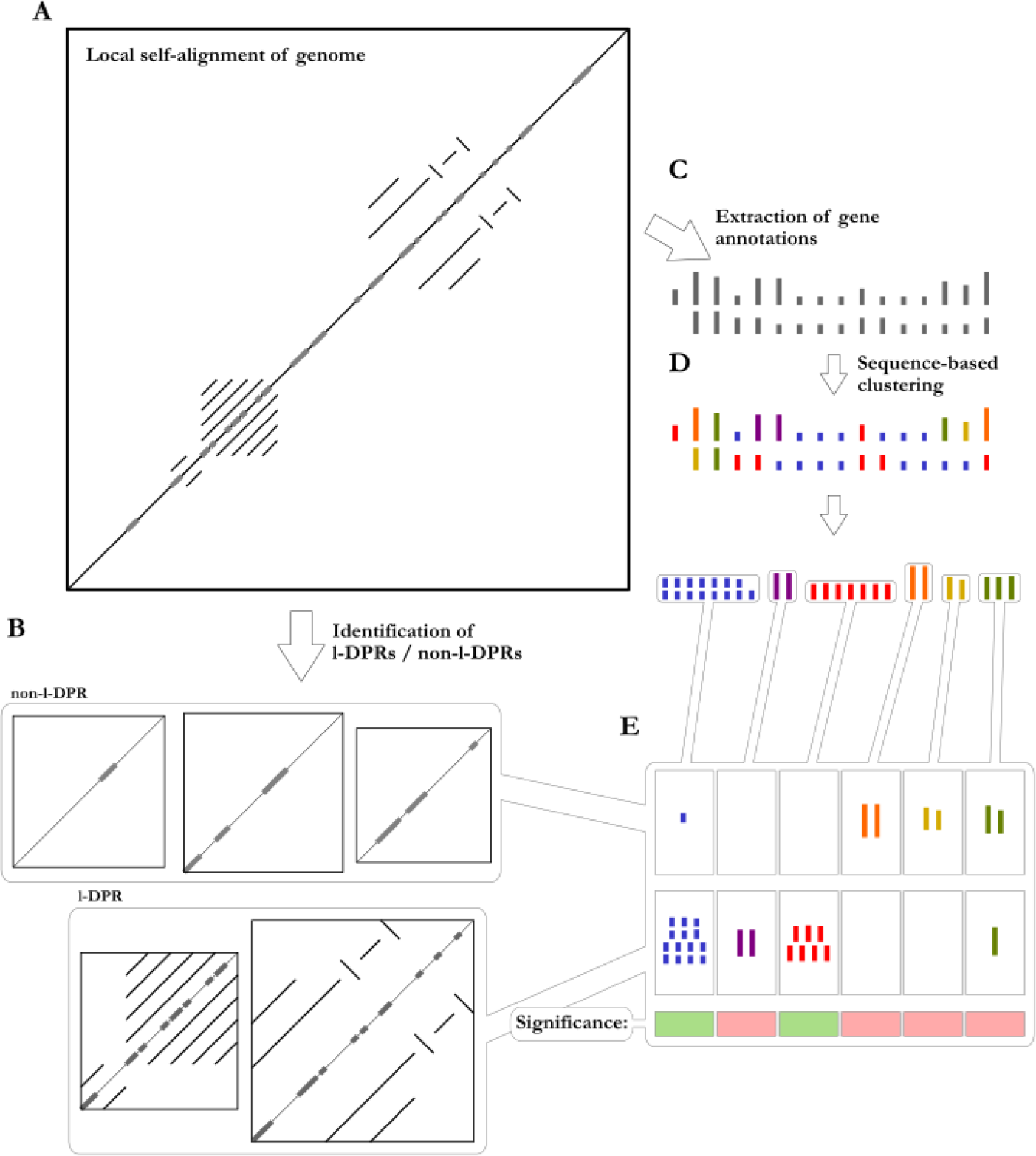
A methods overview for testing the association of l-DPRs with particular gene clusters. Local self-alignments of the genome assembly (A) are used as a basis for classifying genomic ranges as l-DPR or otherwise (B). Genes from the MorexV3 annotation (C) were assigned to gene clusters based on their sequences (D), and the independence of gene cluster membership from l-DPR association was tested for each gene cluster (E).

## Methods

### An appropriate model genome

Identifying candidate RGSUs requires a model organism with several properties. *Firstly*, it must have a large and repetitive genome, and thus can harbour many l-DPRs, sufficient to attain good statistical strength when assessing associations. *Secondly*, to assure that selection is a viable influence on this association, the model’s survival must be highly responsive to disease pressure, and preferably have a short generation time. *Thirdly*, since the accurate reconstruction of repeated sequence is a major flaw of inferior genome assembly methods—and l-DPRs are by definition repetitive—the model must have a genome assembled to a very high standard of completeness and accuracy, using high-fidelity long–read technology. Highly homozygous (e.g. self-pollinating and pure-bred) diploid lines are a major advantage for ensuring maximally accurate assemblies. Diploidy and homozygosity via inbreeding are further advantages since they limit the possibilities for genes to explore the mutational landscape and thus the relative selective advantage of exploiting RGSUs may be elevated. MorexV3, the high-quality assembly of the diploid inbreeding annual crop barley (*Hordeum vulgare*) cultivar ‘Morex’, proves ideal in all these regards^22^.

### Gene-agnostic identification of long-duplication-prone-regions (Supplementary Figures S1)

The pipeline achieves sequence-agnostic identification of l-DPRs based on the assumption that a candidate l-DPR will contain (1) an elevated concentration of (2) locally (3) repeated sequences in the (4) Kbp-scale length range. We first aligned the MorexV3 against itself using lastz^23^ (v1.04.03; arguments: ‘--notransition --step=500 –gapped’). For practicality purposes, this was done in 2 Mbp blocks with a 200 Kbp overlap, and any overlapping l-DPRs identified in multiple windows were merged. For each window, we ignored the trivial end-to-end alignment, and of the remaining alignments, retained only those with a length exceeding 5 Kbp, and falling fully within 200 Kbp of one another. An alignment ‘density’ was calculated over the chromosome by calculating, at ‘interrogation points’ spaced equally at 1 Kb intervals along the length of the chromosome, an alignment density score that is simply the sum of all the lengths of any of the filtered alignments overlapping that interrogation point. A Gaussian kernel density (bandwidth 10 Kbp) was calculated over these the interrogation points, weighted by their scores. To allow comparability between windows, the interrogation point densities were normalised by the sum of scores in the window. Runs of interrogation points at which the density surpassed a minimum density threshold were flagged as l-DPRs. A few minor adjustments to these regions (merging of overlapping regions, and trimming the end coordinates to ensure the stretches always begin and end in repeated sequence) yields the final tabulated list of l-DPR coordinates (Supplementary Tables 1). The pipeline steps following Lastz alignment were implemented in R making significant use of the package data.table^24^ (see Code Availability).

### Function-agnostic assignment of genes to gene clusters

Primary transcript protein sequences from the MorexV3 annotation^22^ were clustered using a global-alignment-based variant of k-means clustering, implemented as the Uclust^25^ (v11) algorithm (Supplementary Tables 2). A clustering cutoff of 0.5 proved adequate to ensure that A) the gene cluster sizes were adequately large for statistical power, and B) the collections of functional descriptions given with the MorexV3 annotation found within each gene cluster tended to be overwhelmingly dominated by a single description (i.e., that within a gene cluster, if a combination of the descriptors ‘MADS-box protein’ and ‘MADS-box-family gene’ is found this is acceptable. If broadly differing descriptors are found, this is undesirable).

### Testing for gene—l-DPR associations

We categorised each member of each gene cluster as falling within- or not-within an l-DPR based on the annotated coordinates of the coding sequence of the longest transcript of each gene (Supplementary Tables 3). Any overlap was considered sufficient to count the gene as within an l-DPR, but we imposed the additional constraint that to be eligible for testing, a gene cluster must contain multiple members within each l-DPR. For each gene cluster, we calculated a p-value on the null hypothesis that membership in the gene cluster was independent of occurrence within l-DPRs (essentially, an urn model) by applying Fisher’s exact test (two sided) (Supplementary Tables 4). Since only non-singleton gene clusters were eligible to show significant association with l-DPRs, we nominated an appropriate p-value cutoff to indicate significance as 0.05 divided by the number of gene clusters possessing more than one member = 0.05/6,419 =~7.79 x 10^-6^.

### Testing for evidence of selective action using gene function information

We required a pool of ‘test’ genes likely to be enriched in functions conferring a selective benefit in arms races. To ensure maximal objectivity, we assigned pool members strictly based on compilations and assignments from the literature and public databases. First, we selected all genes whose GO ontology terms fell under the parent descriptor GO:0044419 “biological process involved in interspecies interaction between organisms”. We added to these genes any with homology-based gene descriptors matching the list of domain descriptors shown to play a role in pathogen resistance in barley and at least one other cereal species, compiled by Krattinger & Keller^26^ (See Supplementary Notes 1 for details on the regular expression used for matching human-readable descriptors). Any gene clusters having greater than 50% of its members in this pool (n=458 of 17,188 gene clusters) was assigned to the probable-arms-race pool. Only 17 of these 458 gene clusters had any members not in the pool. We then tested for significant overlap by fitting a logistical regression model (arms race pool membership ~strength of evidence of association with l-DPRs [-log(p)]), assessing the predictive power of l-DPR association on arms race pool membership with a likelihood ratio test (Supplementary Tables 5).

### Phylogenetic trees for gene clusters

Protein sequences for gene clusters of interest were multiply aligned using MUSCLE (v3.8.31, default parameters) and maximum likelihood trees were constructed using PHYLIP (v 3.696, default parameters).

### Possible functional effects of particular gene clusters under study

While not the main focus of this study, we conducted short investigations to inform speculation on the function of the specific gene clusters that we used as representative examples (Supplementary Notes 2, Supplementary Data, and Supplementary Tables 6). We examined protein structural predictions (ColabFold^27^), the EoRNA Barley gene expression database^28^, and pan-tissue PacBio IsoSeq transcript sequences^29^ aligned to the MorexV3 genome^22^ using BLASTn (v2.10.0; default parameters).

### Investigation of specific candidate RGSU families

Members of the twenty gene clusters with the strongest evidence of l-DPR association (by p-value) were grouped into ‘regions’ wherever they fell within 1 Mbp of each other. For each gene cluster, the regions were pairwise aligned with as described above under “*gene-agnostic identification of long-duplication-prone-regions*”. The alignments were parsed and plotted using custom scripts making extensive use of R base::plot and data.table functions (see Code Availability). Three particularly informative cases that demonstrate general trends seen across these 20 (corresponding to cl_8856, cl_16606, cl_1888) were selected as the basis for discussion in the manuscript, the remainder are shown in Supplementary Data.

## Results

### Genes that associate most strongly with l-DPRs include many known pathogen interaction genes, and selection has affected the composition of candidate RGSUs to favour genes in arms races

Our l-DPR discovery pipeline identified 1,199 candidate l-DPRs (Supplementary Table 1) with lengths ranging between 5.5 and 1,123.598 Kbp (median length 33.600 Kbp). Annotated genes given a “high confidence” (HC) ranking by the annotation pipeline^22^ were assigned to 17,186 clusters comprising between 1 and 727 members each (Supplementary Tables 2—3), most of which (67.2%) were singleton clusters, and 458 of which qualified as members of the ‘arms race’ pool. Note that while HC genes require complete ORFs coupled with evidence from transcript and orthology information^30^, it is still possible that some HC annotations are in reality pseudogenes, gene fragments, or lowly expressed genes. Inspection of the orthology-derived descriptor terms^30^ for l-DPR-associated gene clusters reveals many terms relating to known pathogen-related functions in barley and other cereals, including within the top ten terms (Fig. 3; Supplementary Tables 4) ‘pathogenesis-related protein 1’^31^, ‘jacalin-like lectin’^32^, ‘receptor-like kinase’^33^, ‘jasmonate-induced protein’^34^, ‘thionin’ and ‘thionin-like peptide’^35,36^, and ‘leucine-rich repeat’^37^. Also present among the gene clusters’ common descriptors are ‘alpha/beta-hydrolase superfamily’^38^, whose members have broad but poorly-characterised functions in plants which do include hormone reception, and ‘Cortical cell-delineating protein’^39^, also poorly-studied, which is known to express in the cortical ground meristem of maize roots, accumulating near the region of fastest cellular elongation.

**Figure 3.**
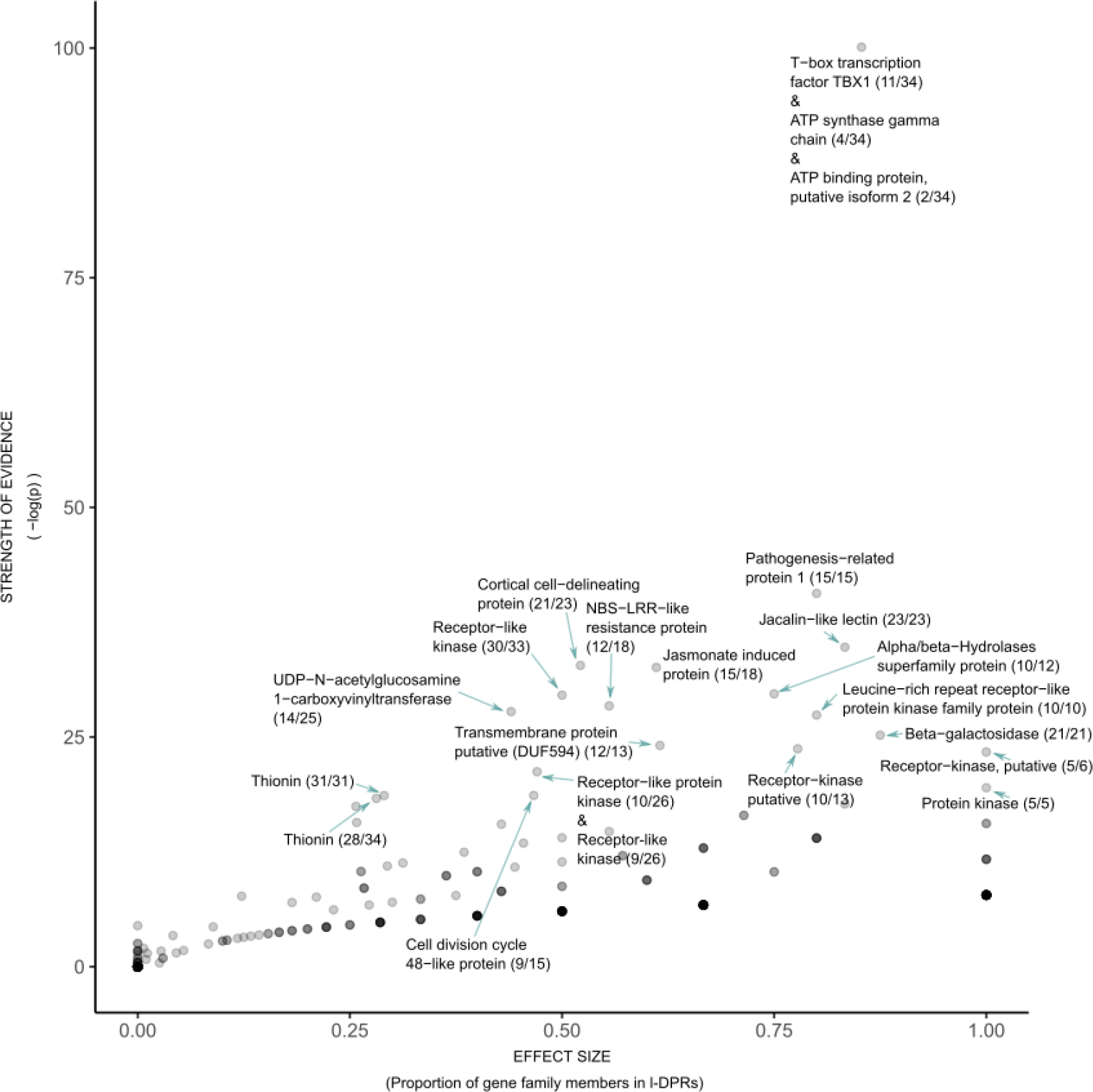
Effect size and strength of evidence for gene cluster association with l-DPRs, i.e. gene clusters whose members occur within l-DPRs, in multiple copies per l-DPR, significantly more frequently than random assignment could account for (points).Gene clusters with strong evidence of association with l-DPRs are marked with their most-common homology-based descriptors as provided by the MorexV3 annotation. In braces: (number of descriptors in gene cluster sharing most common descriptor/total number of gene cluster members). Subsequent analyses show that genes with strong evidence of l-DPR association are significantly more likely to be in the ‘arms-race’ pool (see results). Where the most common descriptor does not account for at least half the descriptors, further descriptors are listed. Note, the majority of unlisted descriptors differ only trivially from the most common capitalisation, hyphens, etc). The strongest evidence of association is given to gene cluster cl_16606, a cluster of very short proteins with mixed homology-based descriptors such as “T-box transcription factor” (a gene family found only in animals). Unlike other gene clusters discussed in this paper, these very short predicted proteins without strong evidence of expression are likely pseudogenes or annotation errors (see investigations in Supplementary Notes 2).

Most importantly, we were able to demonstrate with high confidence that gene clusters with greater evidence of association with l-DPRs are more likely to be in the potential arms-race pool (likelihood ratio test, p<0.003; Supplementary Table 5). Selection based on function is therefore implicated.

### The genomic structure surrounding candidate RGSUs suggest replication-inducers display migrate-and-expand dynamics

Examination of sequence alignment plots between regions containing l-DPR-associated gene clusters (e.g., figs. 4 & 5) suggest several common features. All three gene clusters discussed in the main text (figs. 4 & 5) suggest long-distance dispersal (explaining their presence at disparate genomic sites across multiple chromosomes), followed by tandem replication at the new landing site. Notably, however, the tandemly repeated units at each site are not similar. In the case of gene cluster A, (whose members are annotated as jasmonate-induced proteins, having roles in pathogen response signalling^22,34^)protein phylogeny and alignments (Supplementary Data) confirm; cluster members have been locally duplicated as part of completely distinct tandemly-arrayed motifs—one on chromosome 1H, and two on chromosome 3H. Interestingly, gene cluster A’s array on 1H displays alteration between two variants of the gene in an a-b-a-b-a-b arrangement, suggesting a single ancient tandem duplication event, followed by a pair of recent events that duplicated a larger segment containing the initial duplication to create a nested structure. Gene clusters cl_15653 (members annotated as thionin genes^22^), cl_14902 (annotated as pathogenesis-related proteins^22^), cl_15128 (annotated as cortical cell-delineating proteins), and many others all evidence the same phenomenon (Supplementary Figures S2; Supplementary Materials). The discovery of these tandem-repeat-structuredRGSUs on many different chromosomes therefore suggests they are associated with sequences that induce gene copying across long distances, and induce tandem replication at new transposition landing sites—effectively a transpose-then-replicate stratagem at the genome level, analogous to geographic dispersal and population expansion in ecology. While tandem arrays appear to be a dominant gene duplication mechanism within l-DPRs, duplicated l-DPR genes not falling within a tandemly repeated region tend to be associated with dense local expansions of LTR retrotransposons (recognisable by their canonical “percent-sign” appearance in alignment plots; see figs. 4 & 5; for more examples see gene clusters cl_1888 (members annotated as cell division cycle 48-like proteins), cl_15128 and cl_1274 (annotated as NBS-LRR-like resistance proteins); Supplementary Figures S2; Supplementary Materials).

**Figure 4.**
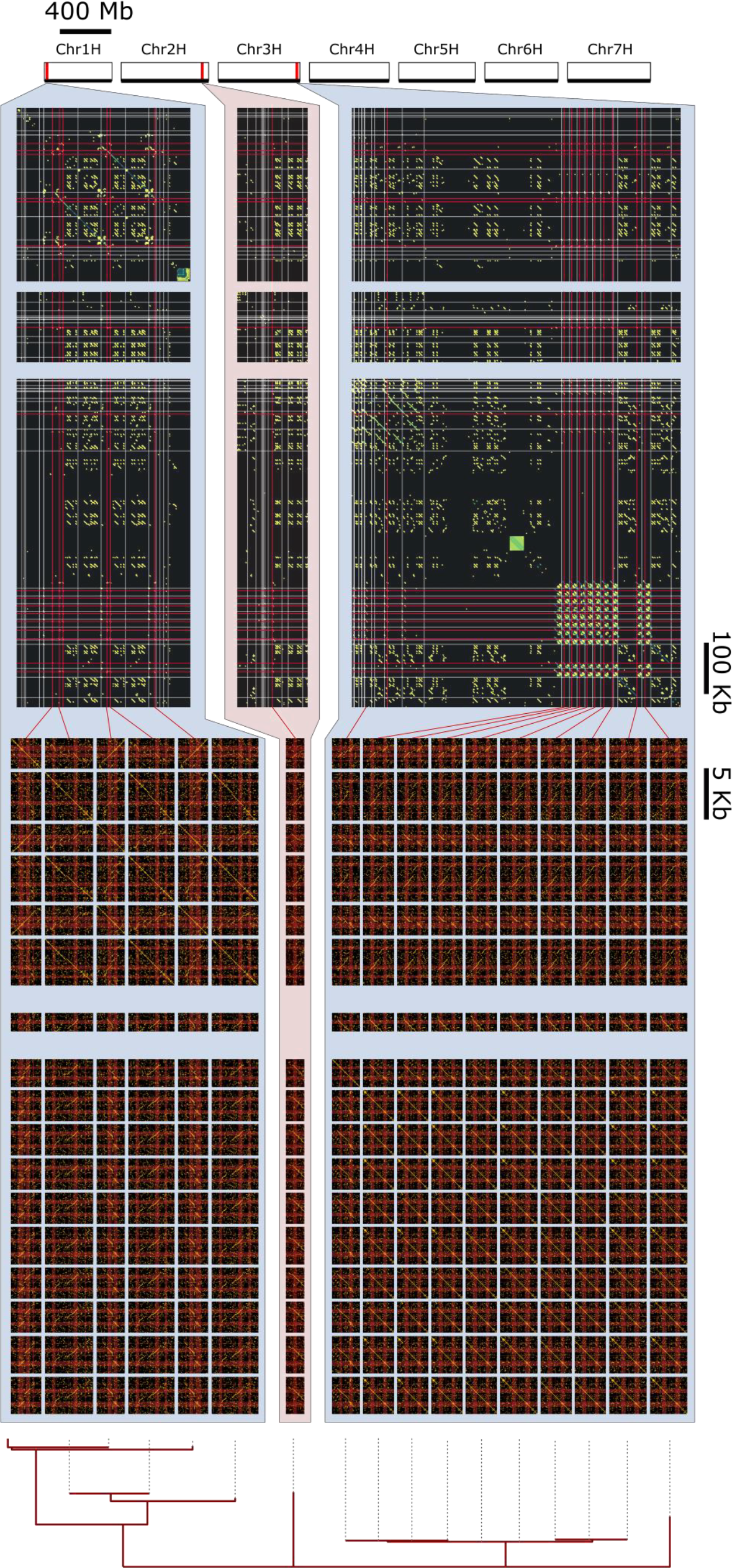
Evolutionary dynamics of a putative RGSU, containing gene cluster A (cl_8856). **Top:** Positions of gene cluster members on barley chromosomes 1H—7H; **Upper middle:** Alignment plots at the l-DPR scale, showing long alignments surrounding the gene cluster members.The positions of genes that are members of gene cluster A are indicated with red bars, white bars indicate genes from other gene clusters. Alignments are coloured according to their similarity, with darker colours having lower similarity. **Lower middle:** Alignment plots showing long alignments surrounding the cluster members; Alignment plots showing each member of gene cluster A with 1 Kbp either side of the coding region either side. Red strips indicate exons. **Bottom:** ML gene tree estimating the ancestral relationships between gene cluster cl_1888 genes based on protein alignment. Arrows, refer to text.

Gene cluster B is included to stimulate discussion on the possible utility specifically of tandem repeat induction as a means of creating gene redundancy. Close homology between spatially separated runs of repeat units clearly reveal the replication of whole subarrays of various sizes. Despite a lack of evidence for cl_16606 being an actively expressed RGSU (see Supplementary Notes 2), the l-DPRs with which it associates make apt examples to describe the mechanism of RGSUs proliferation. L-DPRs containing cl_16606 are discernible on at least five different chromosomes (fig. 5). The tandemly-duplicated unit is almost certainly also the unit that induces translocation, since it is shared between all l-DPRs involved. Furthermore, we can observe evidence that the duplication events can involve up to at least ~12 units at a time, resulting in increased runs of similarity between non-consecutive but equally-distant units owing to their recent common ancestry (yellow arrows in fig. 5). This occurs most strikingly within the chromosome 1H l-DPR shown in figure 5. In arrays on chromosome 5H and 6H, homology between neighbours increases in a graded manner towards one end of the array, suggesting unidirectional recent expansion.

**Figure 5.**
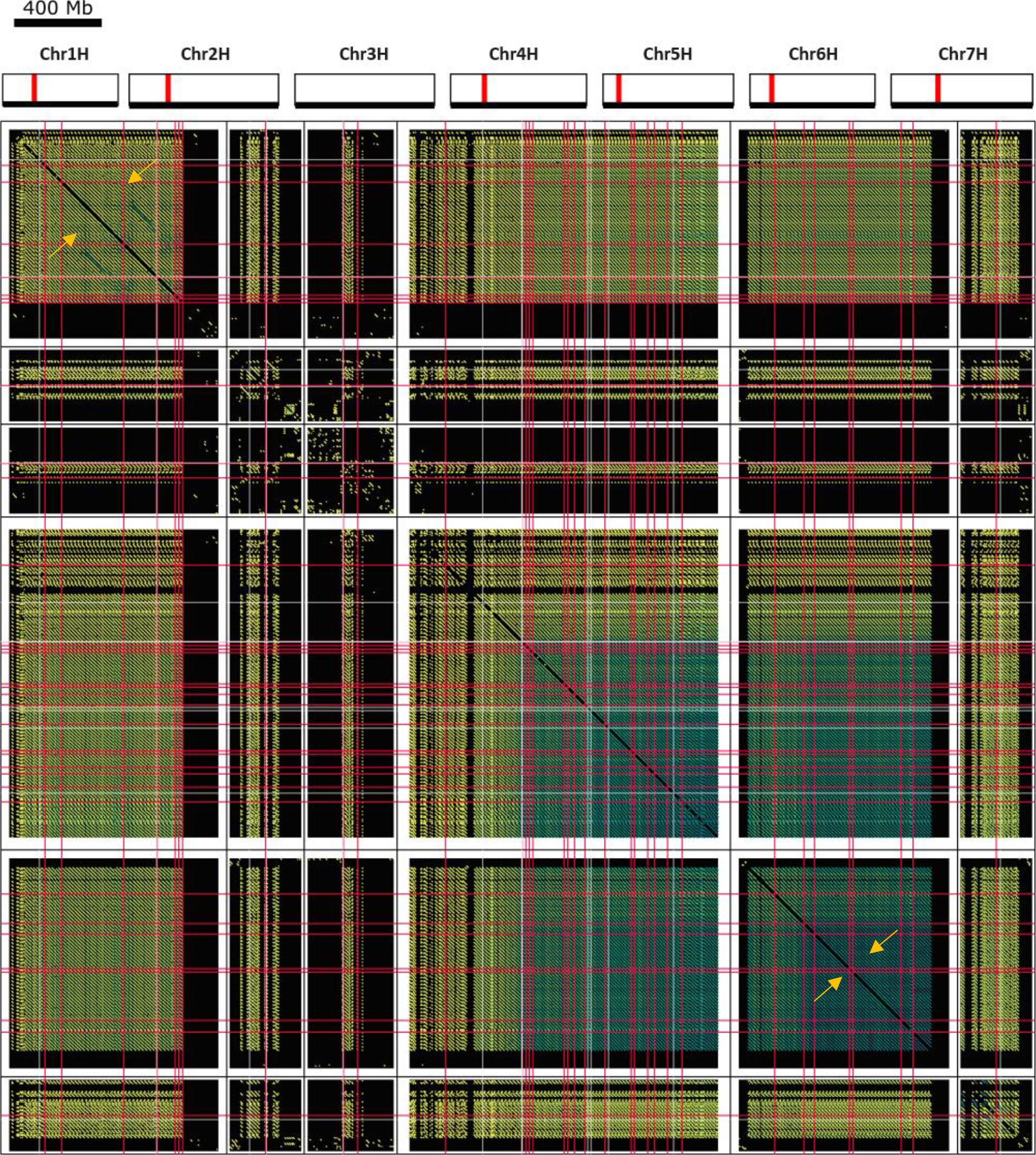
Putative RGSU, containing gene cluster B (cl_16606). Figure features follow the analogous panel in figure 4. Arrows, refer to text.

### Activity of genes in gene clusters A and B, and cl_1888 (Supplementary Figures S2)

There is ample evidence for active expression of gene clusters A and cl_1888. Evidence regarding cluster B’s function and expression is inconclusive. While it is not necessary to the analysis that every candidate RGSUs contain currently functional genes, expression data clearly indicate the activity of several genes identified in candidate RGSUs. The data are presented and discussed in more depth in Supplementary Notes 2, Supplementary Data, and Supplementary Table 6.

## Discussion

### Replicator-gene symbiosis hypothesis

The clear statistical evidence that genes with strong evidence of l-DPR association are significantly more likely to be in the arms race pool demonstrates thatselection has shaped the replicator-associated gene pool in favour of genes involved in plant/pathogen arms races, and is consistent with replicator-gene symbiosis (RGS) being a real phenomenon and an important evolutionary mechanism. A non-significant result would have effectively falsified the RGS hypothesis, at least for the case of arms race genes.

We used pathogen- and interspecific interaction-associated motifs to demonstrate evidence of selective influence based on function where that function was loosely “win arms races”. The observed overlap with these genes is unsurprising given the literature recording the tendency of genes with such functions to occur in large families replicated around particular loci, but the association between these genes and replication-prone DNA can now be asserted as a statistical fact and not just an intuitive hunch.

A few qualifications are nevertheless important. Not all l-DPR-associated genes are expected to be a result of selective advantage (fig. 1), and of those that are, not all are expected to be positively selected based on an arms-race functions specifically. L-DPRs as identified in this study are simply a test pool of regions that appear to be affected by replication-inducers, and do not represent anywhere near the total repetitive DNA content of the genome, which can be above 90%^40^. Under RGS hypothesis, the C-value enigma is,at least partially a result of replication-inducer *tolerance* because *some* replication-inducers are responsible for selectively beneficial outcomes at the organismal and lineage levels, and our result is consistent with this view. As Orgel and Crick wrote in 1980, *“What we would stress is that not all selfish DNA is likely to become useful. Much of it may have no specific function at all. It would be folly in such cases to hunt obsessively for one. To continue our analogy the idea that all human parasites have been selected by human beings for their own advantage*.*”* Only in the era of advanced genome assemblies may this difficulty be circumvented by examining associations of broad gene functional classesover many candidates so that evidence of positive selection becomes apparent at a statistical level.. Future research leveraging the power of pangenome datasets will likely prove invaluable to investigating the details of specific cases.

The results here do not definitively show that replication inducer tolerance is beneficial at the lineage level—there remains the possibility that lineages periodically succumb to genomic infestation by deleterious selfish replicators and selection simply makes the best of a bad situation. But this outlook struggles to explain the correlation of genome size with phylogeny, especially the observation that large and repetitive^41^ genomes have been, in groups such as the grasses (Poeaceae), the ferns (Monilophyta), and cone plants (Gymnospermata), maintained as the norm for hundreds of millions of years^42^. The occasional apomorphic small-genome taxon within these large-genome groups^42^ support the view that replicator suppression is possible, if it is selectively beneficial. And conversely, combined with findings such as those presented here, that replicator tolerance can also be favoured by selection.

### Disperse and expand

Dissimilarity between the tandemly replicated units that cause gene family expansion at disparate genomic locations (as seen in the case of gene clusters A, cl_1888, and others, see Results; fig. 4; Supplementary Figures S2; Supplementary Data) suggests that genes often become associated with replication *inducers* that do not themselves constitute the replicated unit. But what might these inducers be?

While this study is not primarily concerned with molecular mechanisms, observations suggest the importance of several well-understood mechanisms for gene duplication. The first of these is unequal crossing-over, UECO^43–45^ which, when it occurs between any two repeats that flank a locus, can result in gametes in which the entire segment between the two seed repeats is duplicated (fig. 6). It is important to note that, in agreement with the observations seen in fig. 4 and gene clusters including cl_1888 (Supplementary Figures S2), this process duplicates the entire region (including any genes withing the locus) between the two seed repeats. This initial duplication can then produce templates for further UECO events that efficiently lead to copy number variation. Previous studies indicated that certain ‘seed’ repeats of a few 100 bp in length and with ~90% sequence identity are sufficient as templates for UECO^44^. Repeats that drive gene duplication through UECO can be any type of repeated sequence (e.g. a transposable element, TE), which is particularly relevant in large TE-rich genomes. In barley and wheat, 50% of the genome is derived from only about 10 high-copy TE families^17,21^. This, means that there is a high probability that a given gene is flanked by TEs of the same family (and in the same orientation), providing suitable seeds for gene duplication (fig. 6). TE-rich genomes such as that of barley evolve with some base rate of quasi-random regional duplication, analogous to the base rate of other mutations such as SNPs.

**Figure 6.**
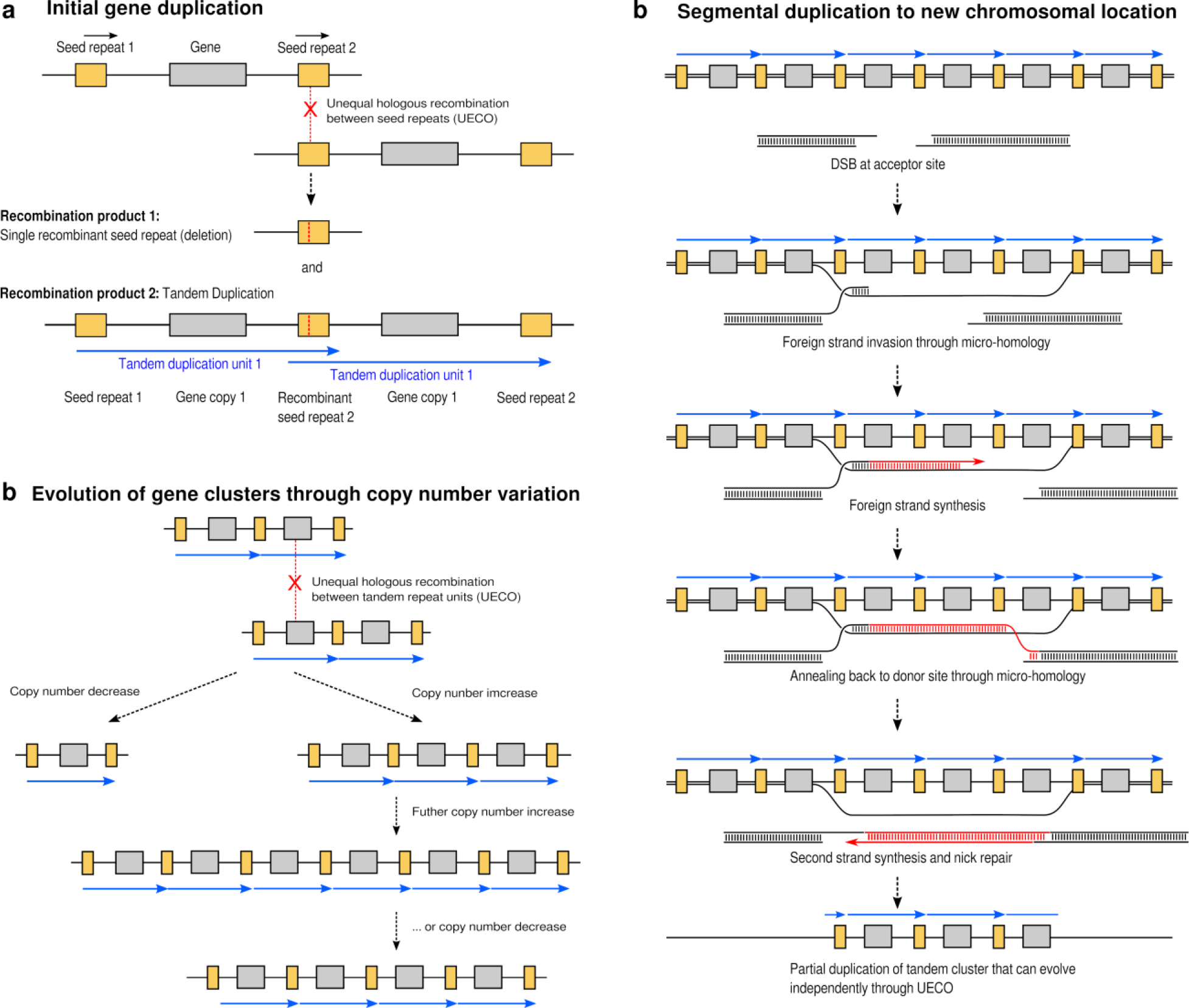
Known mechanisms for the origin and evolution of gene-duplicating l-DPR involving tandem repeats as observed in fig. 4 and cl_1888 (Supplementary Figures S2; see also Supplementary Data).

Gene cluster B (fig. 5) exemplifies an alternative (and, in our data, less common) system in which the dispersed unit does appear to constitute most of the locally-replicated unit, since the tandem repeat it induces is near-identical in all the chromosomal contexts discovered. The observed expansion of these arrays primarily from their ends, which also contain the highest density of annotated genes, is suggestive of an ‘experimental’ front, where successful organismal lineages produce new gene lineages, which are in turn selected based on their influence on fitness at the organism level.

### TEs as self-distributing gene-replication-inducers

The evidence for RGS hypothesis presented here suggests that the ‘selfish’ and ‘adaptive’ role of TEs in the genome could be reconciled as follows: TEs reproduce selfishly, but are tolerated in some lineages owing to the evolvability benefit they confer by creating opportunities for genetic diversity generation, via the mechanism of spreading homologous sequence around the genome to encourage processes such as UECO. The systematic tendencies of various TE families to insert at specific locations relative to genes^21^ may be related to this role. The ubiquity of repeat sequences supports the tantalising possibility that in genomes like barley, adaptive use of replication could constitute the ‘normal’ means of gene space evolution^46,47^. A broad assessment of frequencies of candidate RGSUs in numerous small plant genomes may illuminate whether there is a “critical mass” of repetitive sequences needed for RGSUs to evolve and thrive.

### Possible implications for agriculture

RGS hypothesis sees replicator tolerance as a gamble, where lineages that tolerate small negative effects in the short term tend to discover more beneficial variation that leads them to net increased reproductive success in the long term. This mode of evolution may be antithetical to the canonical process of domestication, in which traits such as yield and homogeneity are selected at the expense of genetic diversity and lineage-level resilience to pathogens and other dynamic threats^11,42^. Occasional observations have recently been made, including recently in maize^3,48^, suggesting excess DNA may be a meaningful contributor to undesirable agricultural traits, and thus there may exist potential applications to agriculture of discovering means to purge cultivated plants’ genomes of repetitive DNA.

## Code Availability

The code used to perform the analyses described is available at github.com/mtrw/RGS.

## Acknowlegements

We are grateful to Mohammed Pourkheirandish, Martin Mascher, and Alexandra Elbakyan for their invaluable comments, support, and contributions to this work.

