## Supplementary Data 2 for "Replicators, genes, and the C-value enigma: High-quality genome assembly of barley provides direct evidence that self-replicating DNA forms ‘cooperative’ associations with genes in arms races"

cl_16606

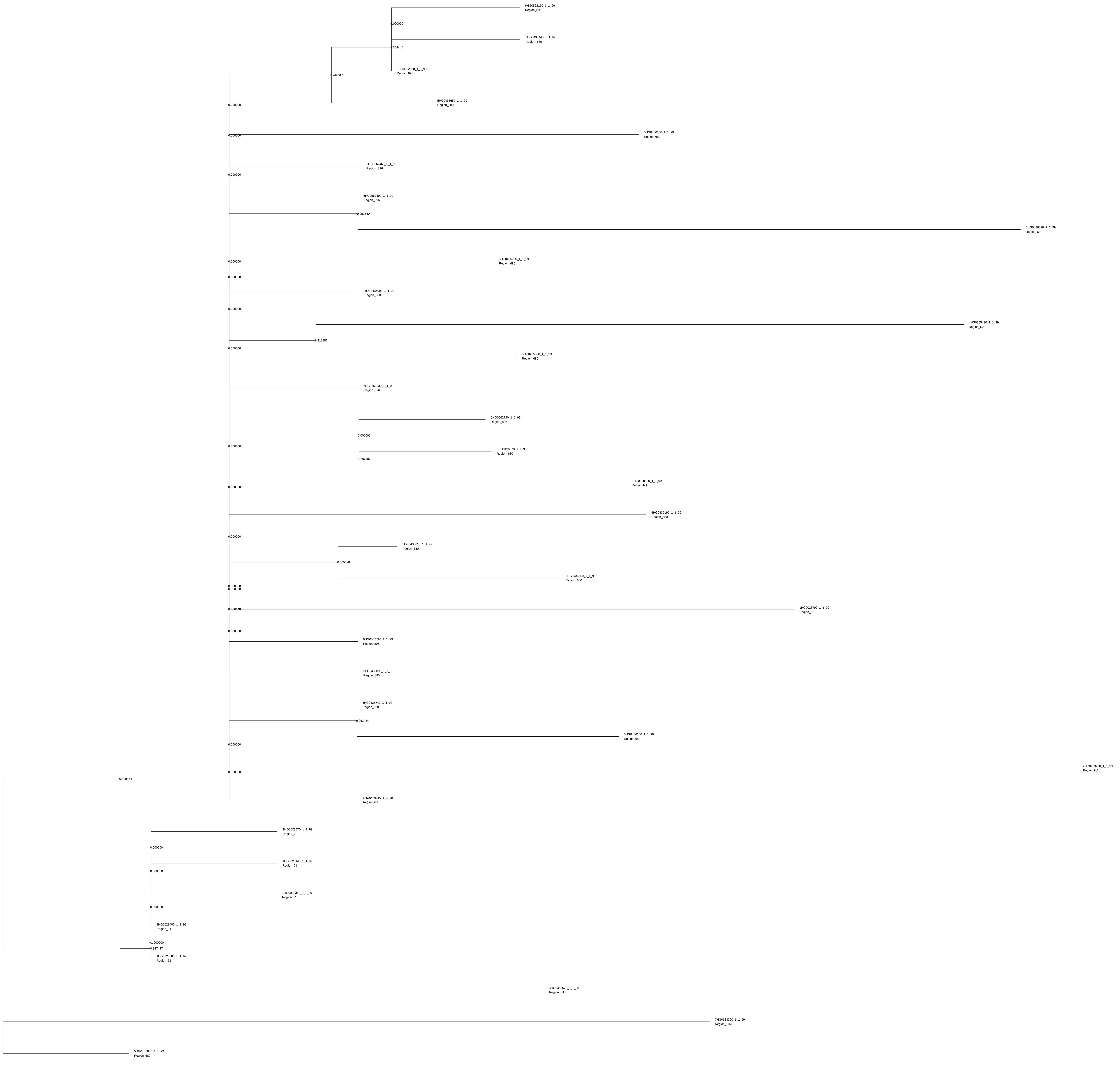

6HG0562150_1_1_99
Region_896
0.000000
5HG0435440_1_1_99
Region_680
0.354445
6HG0562080_1_1_99
Region_896
0.446847
5HG0435650_1_1_99
Region_680
0.000000
5HG0436250_1_1_99
Region_680
0.000000
6HG0562460_1_1_99
Region_896
0.000000
6HG0562490_1_1_99
Region_896
0.951540
5HG0436390_1_1_99
Region_680
0.000000
5HG0435760_1_1_99
Region_680
0.000000
5HG0436090_1_1_99
Region_680
0.000000
4HG0355490_1_1_99
Region_NA
0.412887
0.000000
5HG0435530_1_1_99
Region_680
6HG0562930_1_1_99
Region_896
6HG0562790_1_1_99
Region_896
0.000000
5HG0436070_1_1_99
Region_680
0.000000
0.507169
1HG0028850_1_1_99
Region_NA
0.000000
5HG0436190_1_1_99
Region_680
0.000000
5HG0435610_1_1_99
Region_680
0.505628
5HG0436050_1_1_99
Region_680
0.000000
0.000000
0.426148
1HG0028790_1_1_99
Region_62
0.000000
6HG0562710_1_1_99
Region_896
5HG0436000_1_1_99
Region_680
5HG0435740_1_1_99
Region_680
0.951434
5HG0436180_1_1_99
Region_680
0.000000
0.000000
2HG0133750_1_1_99
Region_NA
5HG0436210_1_1_99
Region_680
0.509574
1HG0028570_1_1_99
Region_62
0.000000
1HG0028440_1_1_99
Region_61
0.000000
1HG0028360_1_1_99
Region_61
0.000000
1HG0028400_1_1_99
Region_61
1.000000
0.337427
1HG0028380_1_1_99
Region_61
4HG0355570_1_1_99
Region_NA
7HG0683380_1_1_99
Region_1075
5HG0435850_1_1_99
Region_680

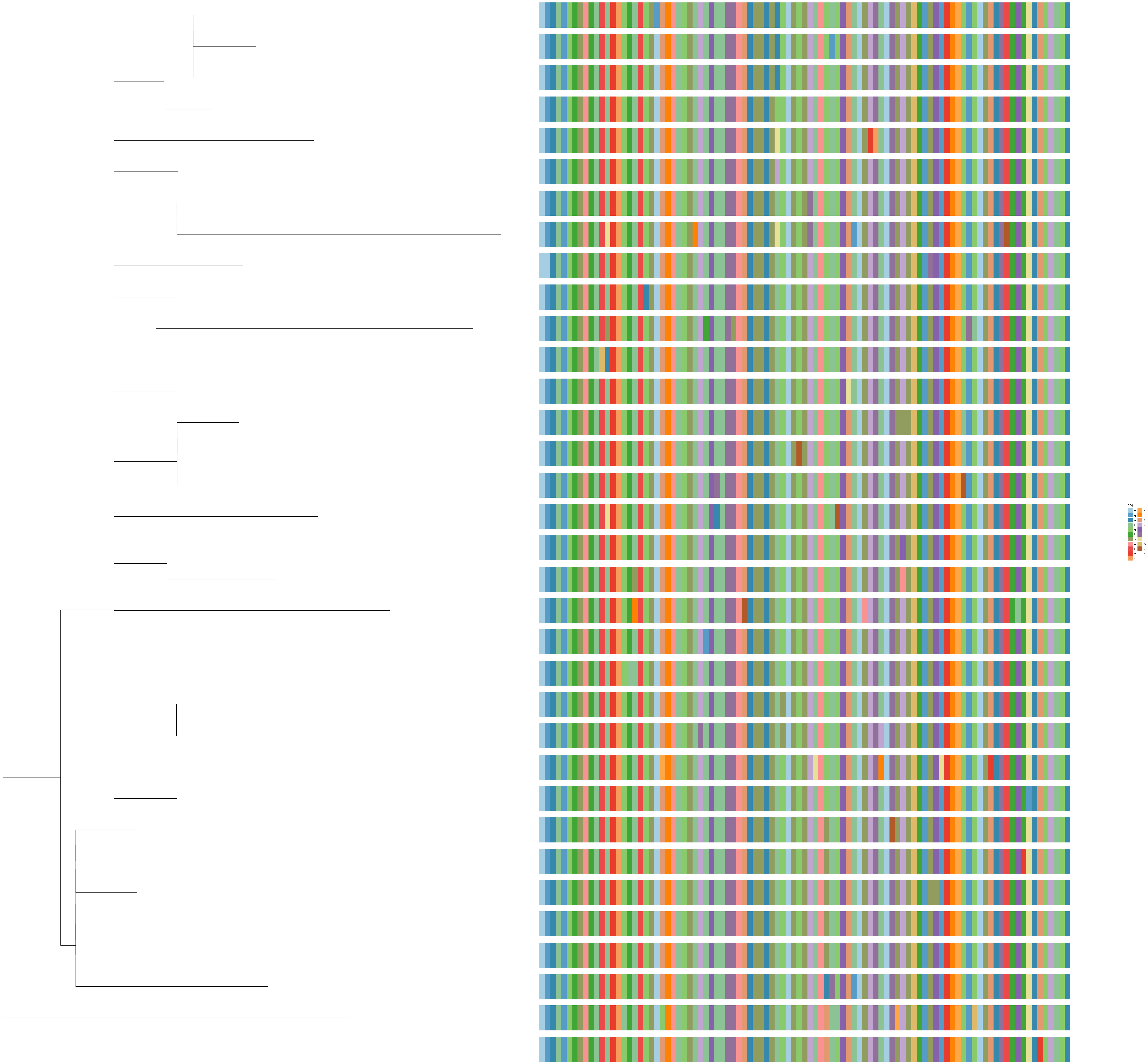
seq
e
q
c
r
g
k
s
a
i
n
f
y
w
d
p
t
l
h
m
v
–

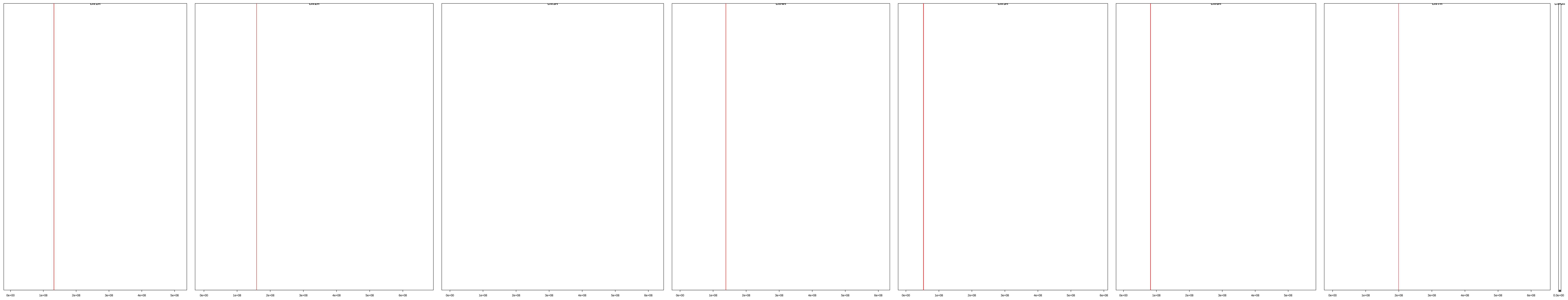

| | description | id | family | seqname | start | end |
|---|---|---|---|---|---|---|
| [1,] | cl_16606 | HORVU.MOREX.r3.1HG0028360.1 | HORVU.MOREX.r3.1HG0028360 | chr1H | 132085892 | 132088188 |
| [2,] | cl_16606 | HORVU.MOREX.r3.1HG0028380.1 | HORVU.MOREX.r3.1HG0028380 | chr1H | 132096518 | 132098814 |
| [3,] | cl_16606 | HORVU.MOREX.r3.1HG0028400.1 | HORVU.MOREX.r3.1HG0028400 | chr1H | 132106658 | 132108954 |
| [4,] | cl_16606 | HORVU.MOREX.r3.1HG0028440.1 | HORVU.MOREX.r3.1HG0028440 | chr1H | 132157388 | 132159684 |
| [5,] | cl_16606 | HORVU.MOREX.r3.1HG0028570.1 | HORVU.MOREX.r3.1HG0028570 | chr1H | 132251522 | 132253818 |
| [6,] | cl_16606 | HORVU.MOREX.r3.1HG0028790.1 | HORVU.MOREX.r3.1HG0028790 | chr1H | 132426611 | 132428907 |
| [7,] | cl_16606 | HORVU.MOREX.r3.1HG0028850.1 | HORVU.MOREX.r3.1HG0028850 | chr1H | 132473768 | 132476064 |
| [8,] | cl_16606 | HORVU.MOREX.r3.2HG0133750.1 | HORVU.MOREX.r3.2HG0133750 | chr2H | 158789295 | 158791591 |
| [9,] | cl_16606 | HORVU.MOREX.r3.4HG0355490.1 | HORVU.MOREX.r3.4HG0355490 | chr4H | 138506020 | 138508316 |
| [10,] | cl_16606 | HORVU.MOREX.r3.4HG0355570.1 | HORVU.MOREX.r3.4HG0355570 | chr4H | 138546573 | 138548869 |
| [11,] | cl_16606 | HORVU.MOREX.r3.5HG0435440.1 | HORVU.MOREX.r3.5HG0435440 | chr5H | 53039603 | 53041899 |
| [12,] | cl_16606 | HORVU.MOREX.r3.5HG0435530.1 | HORVU.MOREX.r3.5HG0435530 | chr5H | 53080936 | 53083232 |
| [13,] | cl_16606 | HORVU.MOREX.r3.5HG0435610.1 | HORVU.MOREX.r3.5HG0435610 | chr5H | 53118180 | 53120476 |
| [14,] | cl_16606 | HORVU.MOREX.r3.5HG0435650.1 | HORVU.MOREX.r3.5HG0435650 | chr5H | 53136802 | 53139098 |
| [15,] | cl_16606 | HORVU.MOREX.r3.5HG0435740.1 | HORVU.MOREX.r3.5HG0435740 | chr5H | 53174045 | 53176341 |
| [16,] | cl_16606 | HORVU.MOREX.r3.5HG0435760.1 | HORVU.MOREX.r3.5HG0435760 | chr5H | 53183355 | 53185651 |
| [17,] | cl_16606 | HORVU.MOREX.r3.5HG0435850.1 | HORVU.MOREX.r3.5HG0435850 | chr5H | 53257843 | 53260139 |
| [18,] | cl_16606 | HORVU.MOREX.r3.5HG0436000.1 | HORVU.MOREX.r3.5HG0436000 | chr5H | 53313700 | 53315996 |
| [19,] | cl_16606 | HORVU.MOREX.r3.5HG0436050.1 | HORVU.MOREX.r3.5HG0436050 | chr5H | 53343124 | 53345420 |
| [20,] | cl_16606 | HORVU.MOREX.r3.5HG0436070.1 | HORVU.MOREX.r3.5HG0436070 | chr5H | 53362422 | 53364718 |
| [21,] | cl_16606 | HORVU.MOREX.r3.5HG0436090.1 | HORVU.MOREX.r3.5HG0436090 | chr5H | 53372408 | 53374704 |
| [22,] | cl_16606 | HORVU.MOREX.r3.5HG0436160.1 | HORVU.MOREX.r3.5HG0436160 | chr5H | 53460651 | 53462947 |
| [23,] | cl_16606 | HORVU.MOREX.r3.5HG0436190.1 | HORVU.MOREX.r3.5HG0436190 | chr5H | 53470638 | 53472934 |
| [24,] | cl_16606 | HORVU.MOREX.r3.5HG0436210.1 | HORVU.MOREX.r3.5HG0436210 | chr5H | 53480610 | 53482906 |
| [25,] | cl_16606 | HORVU.MOREX.r3.5HG0436250.1 | HORVU.MOREX.r3.5HG0436250 | chr5H | 53490597 | 53492893 |
| [26,] | cl_16606 | HORVU.MOREX.r3.5HG0436390.1 | HORVU.MOREX.r3.5HG0436390 | chr5H | 53707733 | 53710029 |
| [27,] | cl_16606 | HORVU.MOREX.r3.6HG0562080.1 | HORVU.MOREX.r3.6HG0562080 | chr6H | 81974994 | 81978953 |
| [28,] | cl_16606 | HORVU.MOREX.r3.6HG0562150.1 | HORVU.MOREX.r3.6HG0562150 | chr6H | 82009252 | 82011548 |
| [29,] | cl_16606 | HORVU.MOREX.r3.6HG0562460.1 | HORVU.MOREX.r3.6HG0562460 | chr6H | 82145470 | 82147766 |
| [30,] | cl_16606 | HORVU.MOREX.r3.6HG0562490.1 | HORVU.MOREX.r3.6HG0562490 | chr6H | 82155303 | 82157599 |
| [31,] | cl_16606 | HORVU.MOREX.r3.6HG0562710.1 | HORVU.MOREX.r3.6HG0562710 | chr6H | 82253641 | 82255937 |
| [32,] | cl_16606 | HORVU.MOREX.r3.6HG0562790.1 | HORVU.MOREX.r3.6HG0562790 | chr6H | 82283146 | 82285442 |
| [33,] | cl_16606 | HORVU.MOREX.r3.6HG0562930.1 | HORVU.MOREX.r3.6HG0562930 | chr6H | 82366130 | 82368426 |
| [34,] | cl_16606 | HORVU.MOREX.r3.7HG0683360.1 | HORVU.MOREX.r3.7HG0683360 | chr7H | 199423259 | 199425555 |

cl_14902

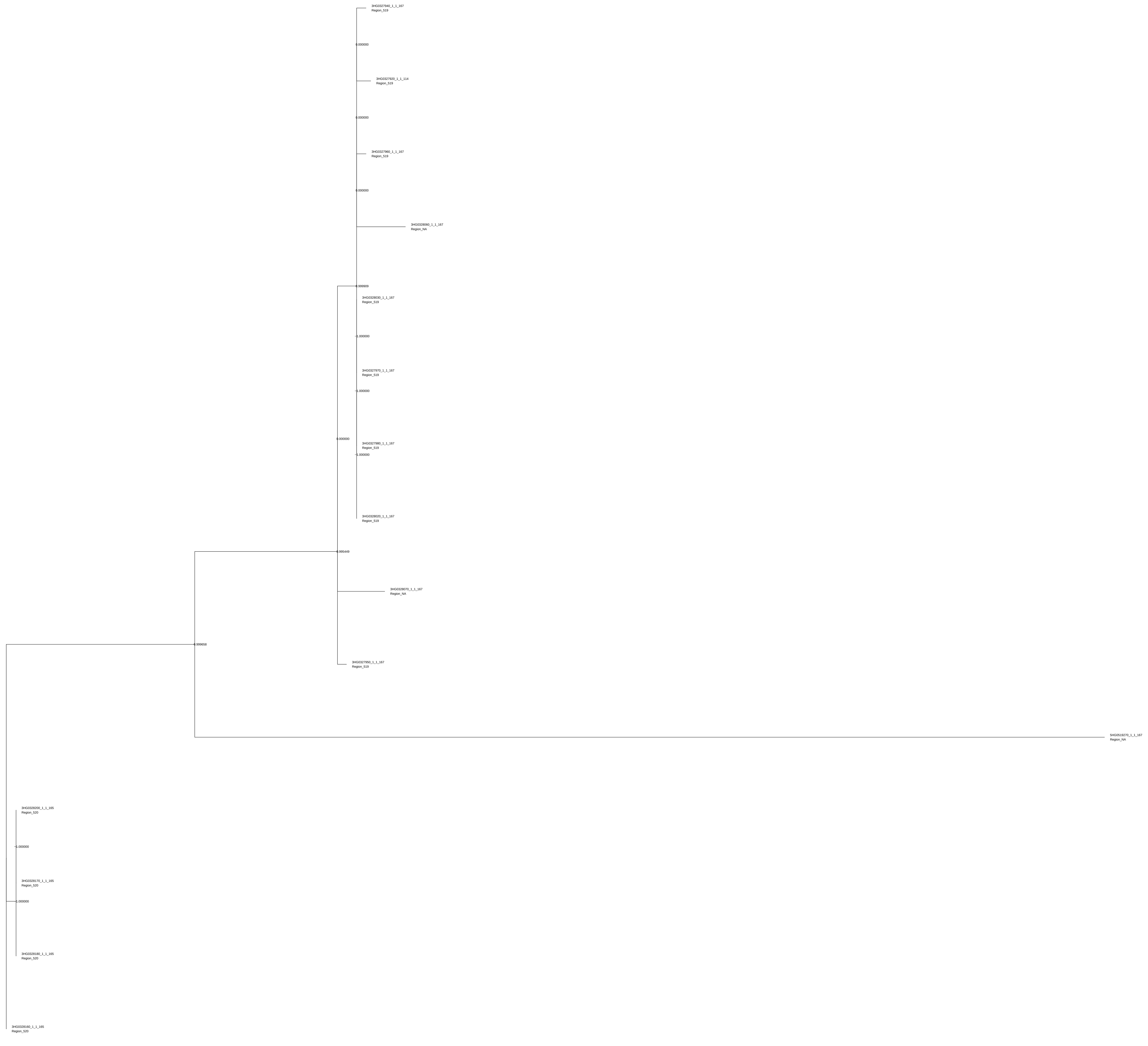
3HG0327940_1_1_167
Region_519
0.000000
3HG0327920_1_1_114
Region_519
0.000000
3HG0327960_1_1_167
Region_519
0.000000
3HG0328060_1_1_167
Region_NA
0.999909
3HG0328030_1_1_167
Region_519
1.000000
3HG0327970_1_1_167
Region_519
1.000000
0.000000
3HG0327980_1_1_167
Region_519
1.000000
3HG0328020_1_1_167
Region_519
0.995449
3HG0328070_1_1_167
Region_NA
0.999658
3HG0327950_1_1_167
Region_519
5HG0519270_1_1_167
Region_NA
3HG0328200_1_1_165
Region_520
1.000000
3HG0328170_1_1_165
Region_520
1.000000
3HG0328180_1_1_165
Region_520
3HG0328160_1_1_165
Region_520

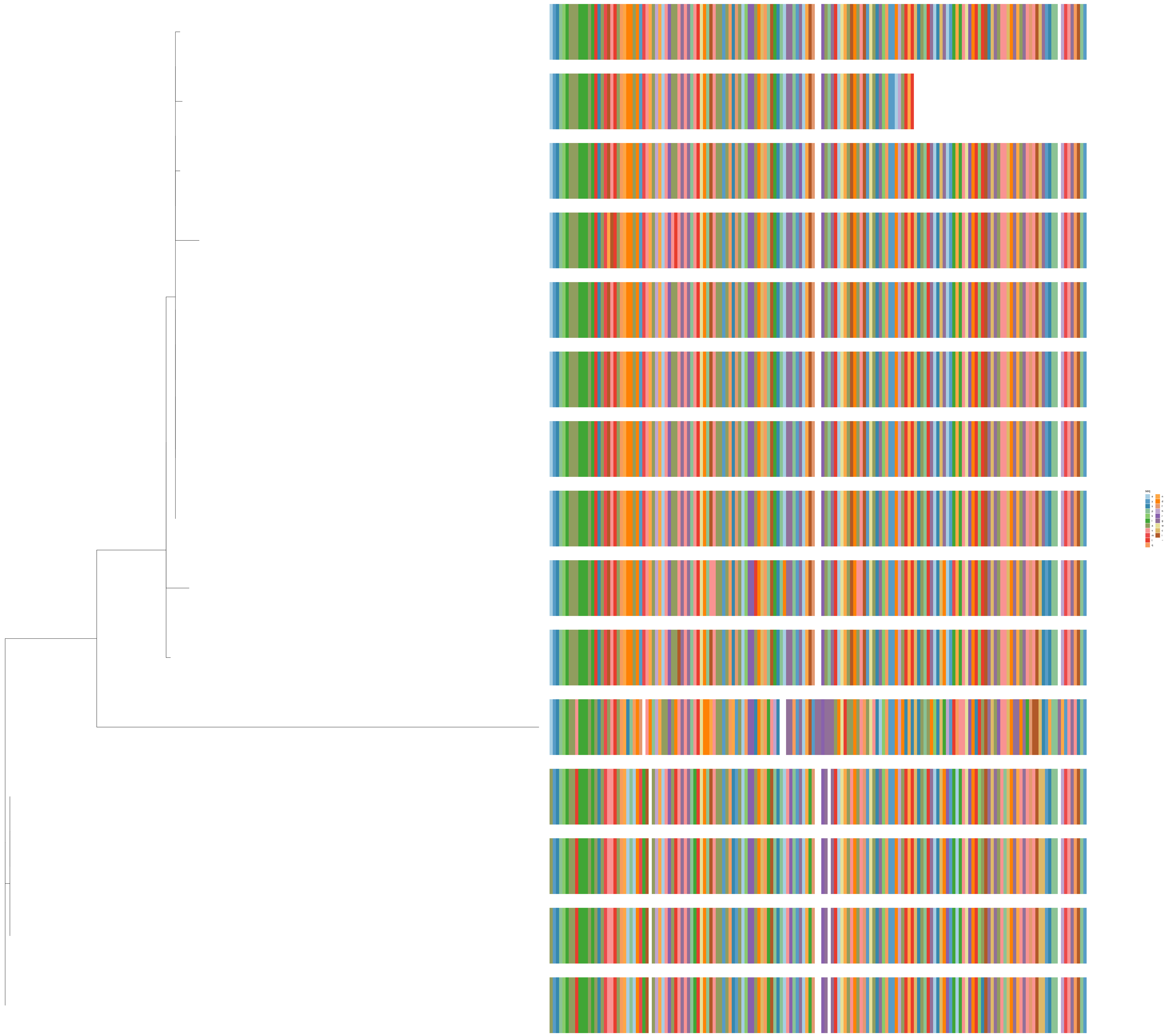
seq
e
y
s
p
k
l
a
v
m
t
q
n
d
f
h
r
g
w
c
i
-

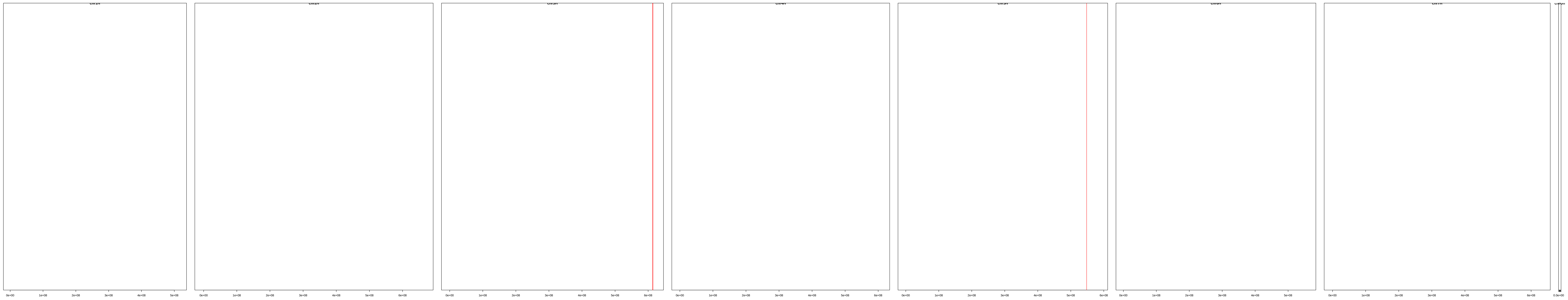

| | description | id | family | seqname | start | end |
|---|---|---|---|---|---|---|
| [1,] | cl_14902 | HORVU.MOREX.r3.3HG0327920.1 | HORVU.MOREX.r3.3HG0327920 | chr3H | 613790761 | 613793102 |
| [2,] | cl_14902 | HORVU.MOREX.r3.3HG0327940.1 | HORVU.MOREX.r3.3HG0327940 | chr3H | 613824925 | 613827425 |
| [3,] | cl_14902 | HORVU.MOREX.r3.3HG0327950.1 | HORVU.MOREX.r3.3HG0327950 | chr3H | 613843191 | 613845691 |
| [4,] | cl_14902 | HORVU.MOREX.r3.3HG0327960.1 | HORVU.MOREX.r3.3HG0327960 | chr3H | 613856384 | 613858884 |
| [5,] | cl_14902 | HORVU.MOREX.r3.3HG0327970.1 | HORVU.MOREX.r3.3HG0327970 | chr3H | 613860134 | 613862634 |
| [6,] | cl_14902 | HORVU.MOREX.r3.3HG0327980.1 | HORVU.MOREX.r3.3HG0327980 | chr3H | 613863884 | 613866384 |
| [7,] | cl_14902 | HORVU.MOREX.r3.3HG0328020.1 | HORVU.MOREX.r3.3HG0328020 | chr3H | 613890988 | 613893488 |
| [8,] | cl_14902 | HORVU.MOREX.r3.3HG0328030.1 | HORVU.MOREX.r3.3HG0328030 | chr3H | 613918093 | 613920593 |
| [9,] | cl_14902 | HORVU.MOREX.r3.3HG0328060.1 | HORVU.MOREX.r3.3HG0328060 | chr3H | 613961794 | 613964294 |
| [10,] | cl_14902 | HORVU.MOREX.r3.3HG0328070.1 | HORVU.MOREX.r3.3HG0328070 | chr3H | 613975102 | 613977602 |
| [11,] | cl_14902 | HORVU.MOREX.r3.3HG0328160.1 | HORVU.MOREX.r3.3HG0328160 | chr3H | 614255235 | 614257729 |
| [12,] | cl_14902 | HORVU.MOREX.r3.3HG0328170.1 | HORVU.MOREX.r3.3HG0328170 | chr3H | 614260430 | 614263155 |
| [13,] | cl_14902 | HORVU.MOREX.r3.3HG0328180.1 | HORVU.MOREX.r3.3HG0328180 | chr3H | 614306430 | 614308924 |
| [14,] | cl_14902 | HORVU.MOREX.r3.3HG0328200.1 | HORVU.MOREX.r3.3HG0328200 | chr3H | 614367396 | 614369890 |
| [15,] | cl_14902 | HORVU.MOREX.r3.5HG0519270.1 | HORVU.MOREX.r3.5HG0519270 | chr5H | 548021287 | 548024048 |

cl_10965

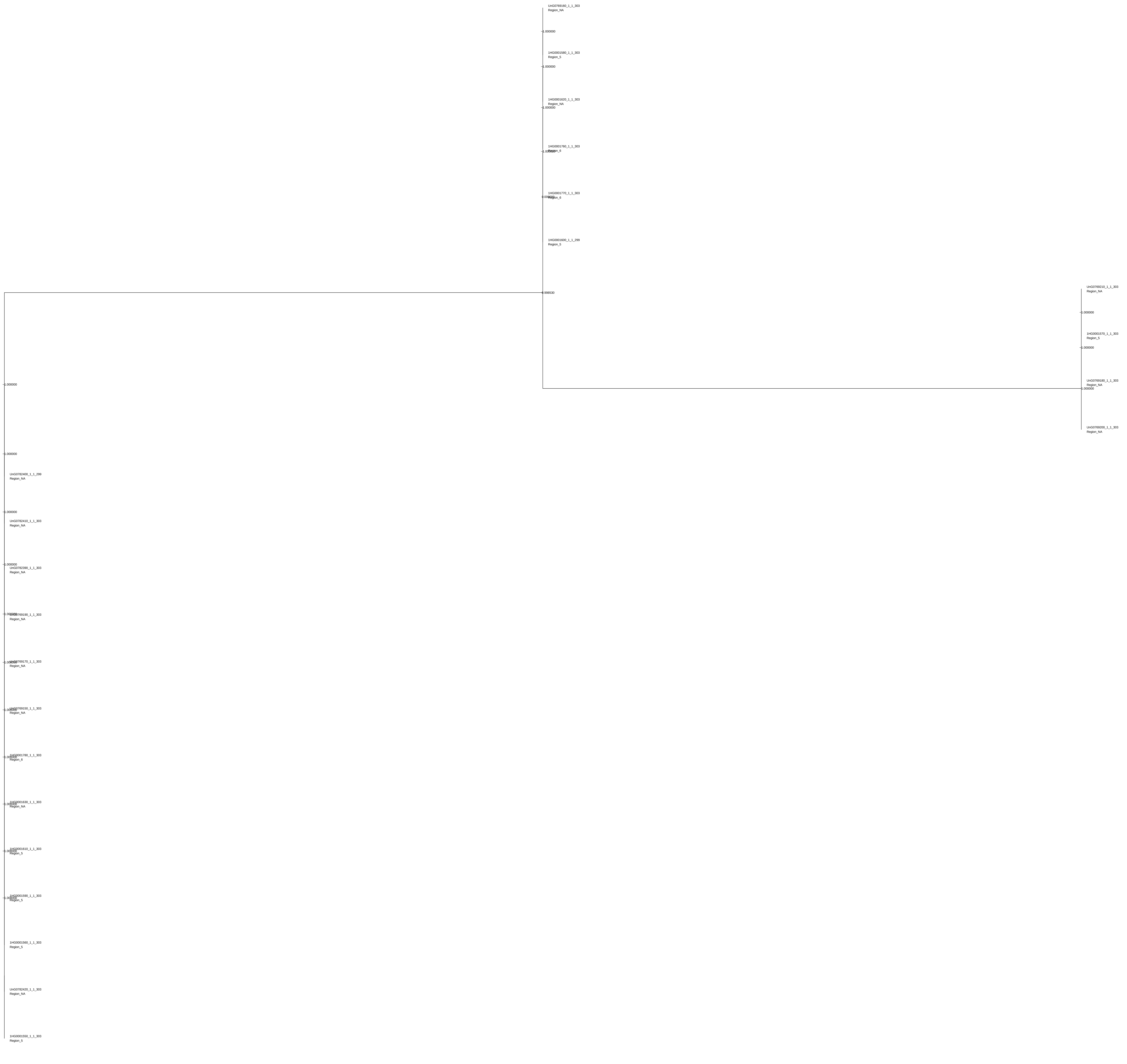

UnG0769160_1_1_303
Region_NA
1.000000
1HG0001580_1_1_303
Region_5
1.000000
1HG0001620_1_1_303
Region_NA
1.000000
1HG0001760_1_1_303
Region_6
1HG0001770_1_1_303
Region_6
1HG0001600_1_1_299
Region_5
0.998530
UnG0769210_1_1_303
Region_NA
1.000000
1HG0001570_1_1_303
Region_5
1.000000
UnG0769180_1_1_303
Region_NA
1.000000
UnG0769200_1_1_303
Region_NA
1.000000
1.000000
UnG0782400_1_1_299
Region_NA
1.000000
UnG0782410_1_1_303
Region_NA
1.000000
UnG0782390_1_1_303
Region_NA
Region_NA
Region_NA
Region_NA
Region_6
Region_NA
Region_5
Region_5
1HG0001560_1_1_303
Region_5
UnG0782420_1_1_303
Region_NA
1HG0001550_1_1_303
Region_5

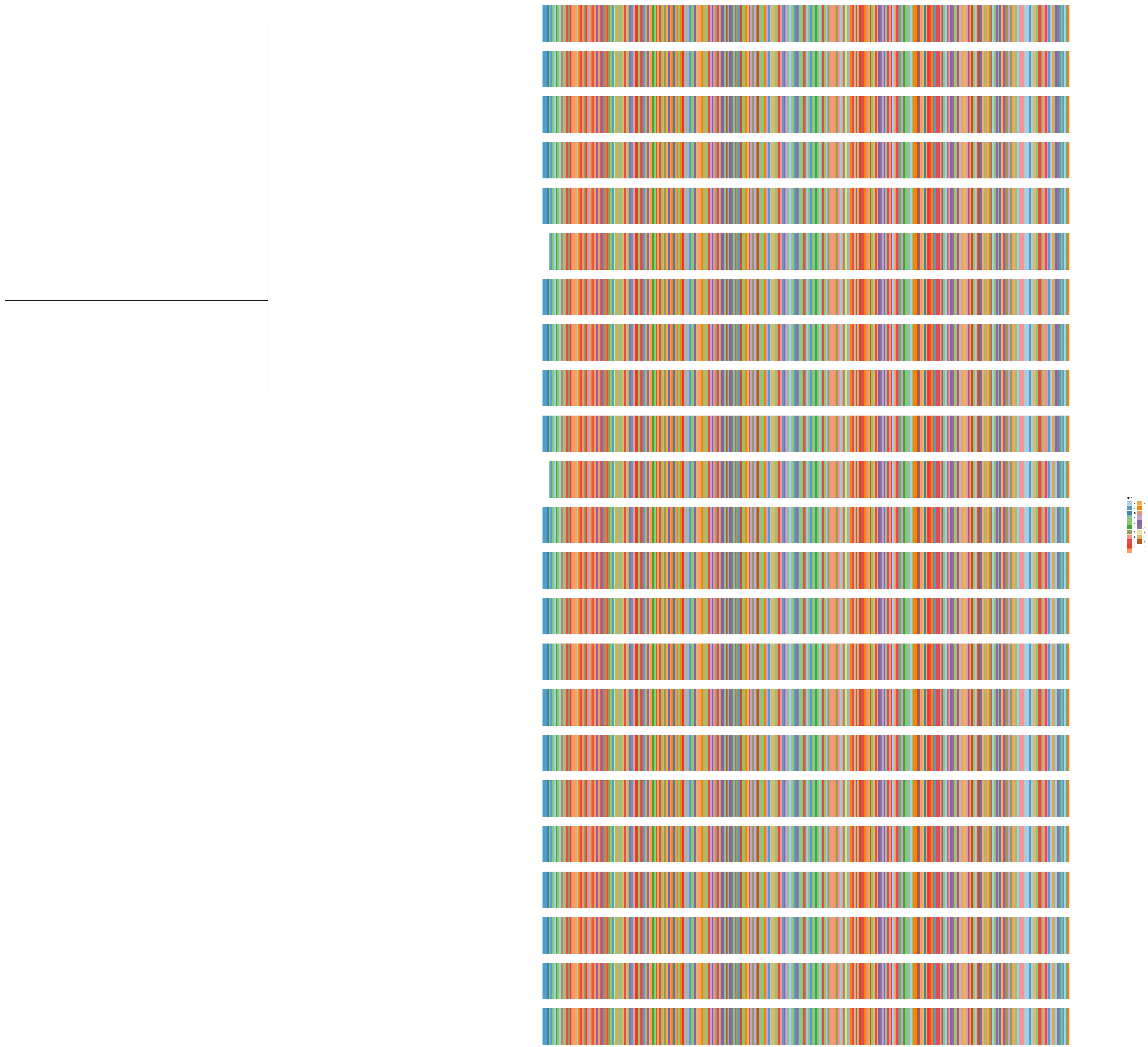
seq
a
v
m
k
g
w
p
d
s
q
i
n
e
r
l
t
y
c
f
h
–

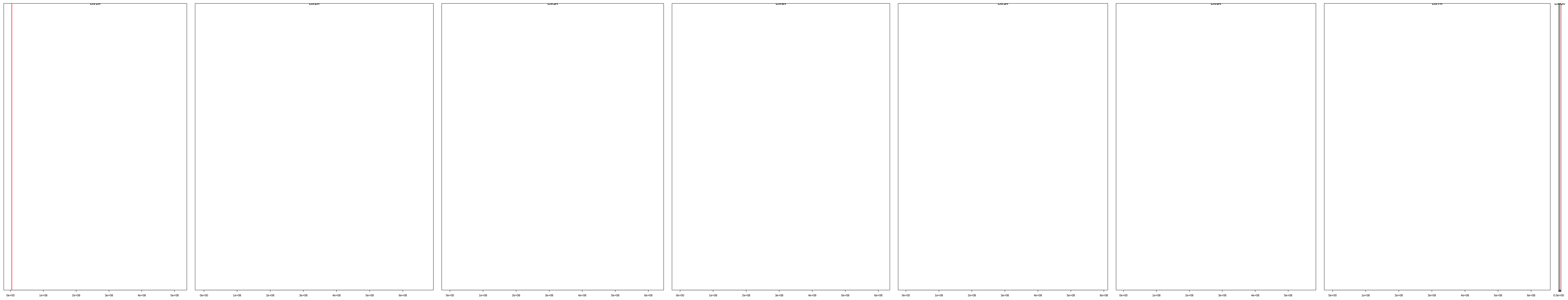

| | description | id | family | seqname | start | end |
|---|---|---|---|---|---|---|
| [1,] | cl_10965 | HORVU.MOREX.r3.1HG0001550.1 | HORVU.MOREX.r3.1HG0001550 | chr1H | 3689462 | 3692635 |
| [2,] | cl_10965 | HORVU.MOREX.r3.1HG0001560.1 | HORVU.MOREX.r3.1HG0001560 | chr1H | 3704388 | 3707834 |
| [3,] | cl_10965 | HORVU.MOREX.r3.1HG0001570.1 | HORVU.MOREX.r3.1HG0001570 | chr1H | 3718941 | 3722114 |
| [4,] | cl_10965 | HORVU.MOREX.r3.1HG0001580.1 | HORVU.MOREX.r3.1HG0001580 | chr1H | 3733860 | 3737033 |
| [5,] | cl_10965 | HORVU.MOREX.r3.1HG0001590.1 | HORVU.MOREX.r3.1HG0001590 | chr1H | 3747366 | 3750539 |
| [6,] | cl_10965 | HORVU.MOREX.r3.1HG0001600.1 | HORVU.MOREX.r3.1HG0001600 | chr1H | 3762109 | 3765086 |
| [7,] | cl_10965 | HORVU.MOREX.r3.1HG0001610.1 | HORVU.MOREX.r3.1HG0001610 | chr1H | 3776835 | 3780008 |
| [8,] | cl_10965 | HORVU.MOREX.r3.1HG0001620.1 | HORVU.MOREX.r3.1HG0001620 | chr1H | 3791417 | 3794590 |
| [9,] | cl_10965 | HORVU.MOREX.r3.1HG0001630.1 | HORVU.MOREX.r3.1HG0001630 | chr1H | 3811084 | 3814257 |
| [10,] | cl_10965 | HORVU.MOREX.r3.1HG0001760.1 | HORVU.MOREX.r3.1HG0001760 | chr1H | 4069036 | 4072209 |
| [11,] | cl_10965 | HORVU.MOREX.r3.1HG0001770.1 | HORVU.MOREX.r3.1HG0001770 | chr1H | 4083612 | 4086785 |
| [12,] | cl_10965 | HORVU.MOREX.r3.1HG0001780.1 | HORVU.MOREX.r3.1HG0001780 | chr1H | 4103282 | 4106455 |
| [13,] | cl_10965 | HORVU.MOREX.r3.UnG0769150.1 | HORVU.MOREX.r3.UnG0769150 | chrUn | 10537462 | 10540635 |
| [14,] | cl_10965 | HORVU.MOREX.r3.UnG0769160.1 | HORVU.MOREX.r3.UnG0769160 | chrUn | 10557132 | 10560305 |
| [15,] | cl_10965 | HORVU.MOREX.r3.UnG0769170.1 | HORVU.MOREX.r3.UnG0769170 | chrUn | 10571713 | 10574886 |
| [16,] | cl_10965 | HORVU.MOREX.r3.UnG0769180.1 | HORVU.MOREX.r3.UnG0769180 | chrUn | 10586635 | 10589808 |
| [17,] | cl_10965 | HORVU.MOREX.r3.UnG0769190.1 | HORVU.MOREX.r3.UnG0769190 | chrUn | 10601185 | 10604358 |
| [18,] | cl_10965 | HORVU.MOREX.r3.UnG0769200.1 | HORVU.MOREX.r3.UnG0769200 | chrUn | 10614691 | 10617864 |
| [19,] | cl_10965 | HORVU.MOREX.r3.UnG0769210.1 | HORVU.MOREX.r3.UnG0769210 | chrUn | 10629616 | 10632789 |
| [20,] | cl_10965 | HORVU.MOREX.r3.UnG0782390.1 | HORVU.MOREX.r3.UnG0782390 | chrUn | 15813349 | 15816861 |
| [21,] | cl_10965 | HORVU.MOREX.r3.UnG0782400.1 | HORVU.MOREX.r3.UnG0782400 | chrUn | 15828560 | 15831537 |
| [22,] | cl_10965 | HORVU.MOREX.r3.UnG0782410.1 | HORVU.MOREX.r3.UnG0782410 | chrUn | 15843416 | 15846589 |
| [23,] | cl_10965 | HORVU.MOREX.r3.UnG0782420.1 | HORVU.MOREX.r3.UnG0782420 | chrUn | 15861421 | 15864594 |

cl_15128

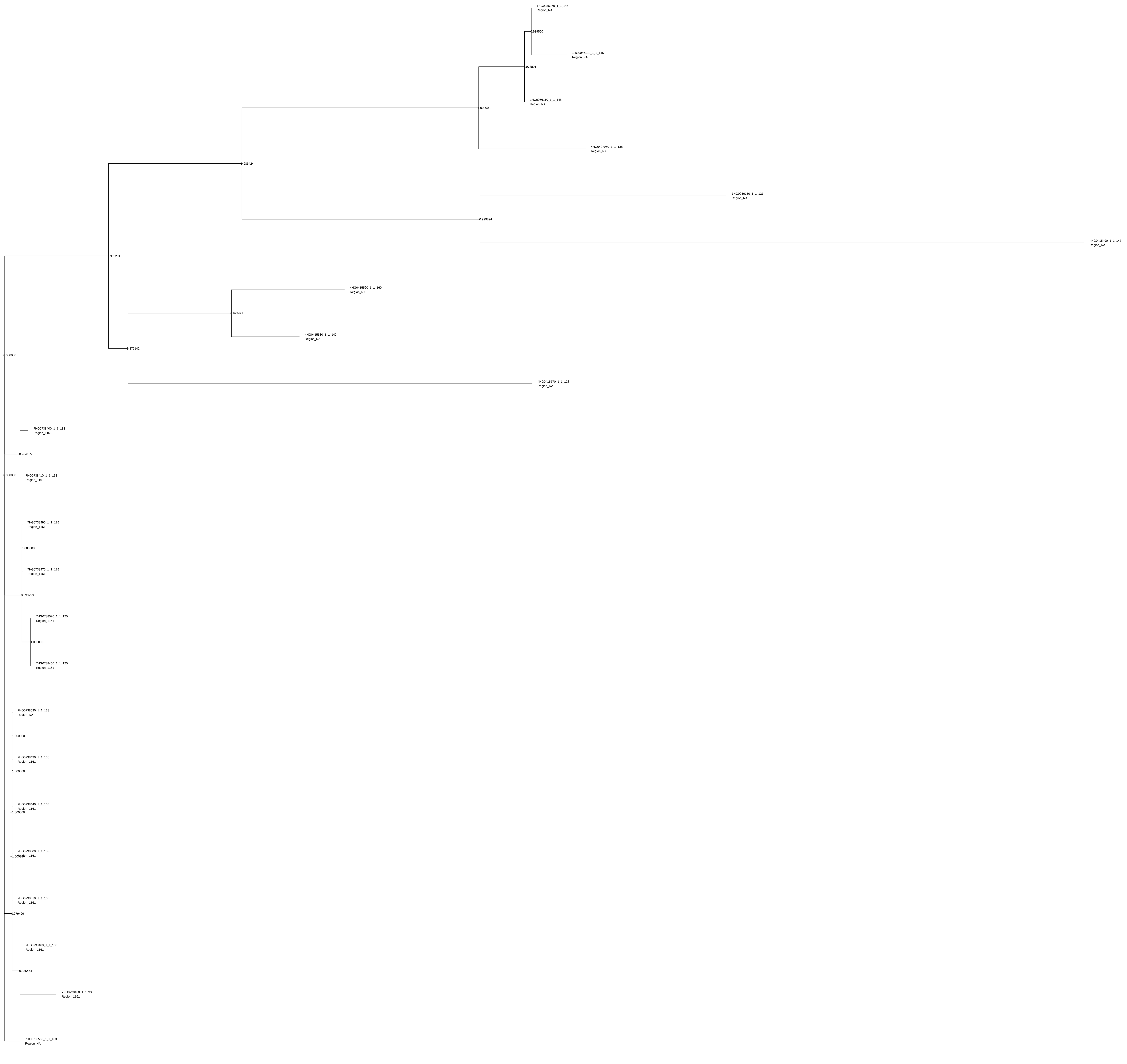

1HG0056070_1_1_145
Region_NA
0.939550
1HG0056130_1_1_145
Region_NA
0.973801
1HG0056110_1_1_145
Region_NA
1.000000
4HG0407950_1_1_138
Region_NA
0.986424
1HG0056150_1_1_121
Region_NA
0.999894
4HG0415490_1_1_147
Region_NA
0.999291
4HG0415520_1_1_160
Region_NA
0.999471
4HG0415530_1_1_140
Region_NA
0.372142
0.000000
4HG0415570_1_1_128
Region_NA
7HG0738400_1_1_133
Region_1161
0.984185
0.000000
7HG0738410_1_1_133
Region_1161
7HG0738490_1_1_125
Region_1161
1.000000
7HG0738470_1_1_125
Region_1161
0.999759
7HG0738520_1_1_125
Region_1161
1.000000
7HG0738450_1_1_125
Region_1161
7HG0738530_1_1_133
Region_NA
1.000000
7HG0738430_1_1_133
Region_1161
1.000000
7HG0738440_1_1_133
Region_1161
1.000000
7HG0738500_1_1_133
Region_1161
1.000000
7HG0738510_1_1_133
Region_1161
0.979499
7HG0738460_1_1_133
Region_1161
0.335474
7HG0738480_1_1_93
Region_1161
7HG0738560_1_1_133
Region_NA

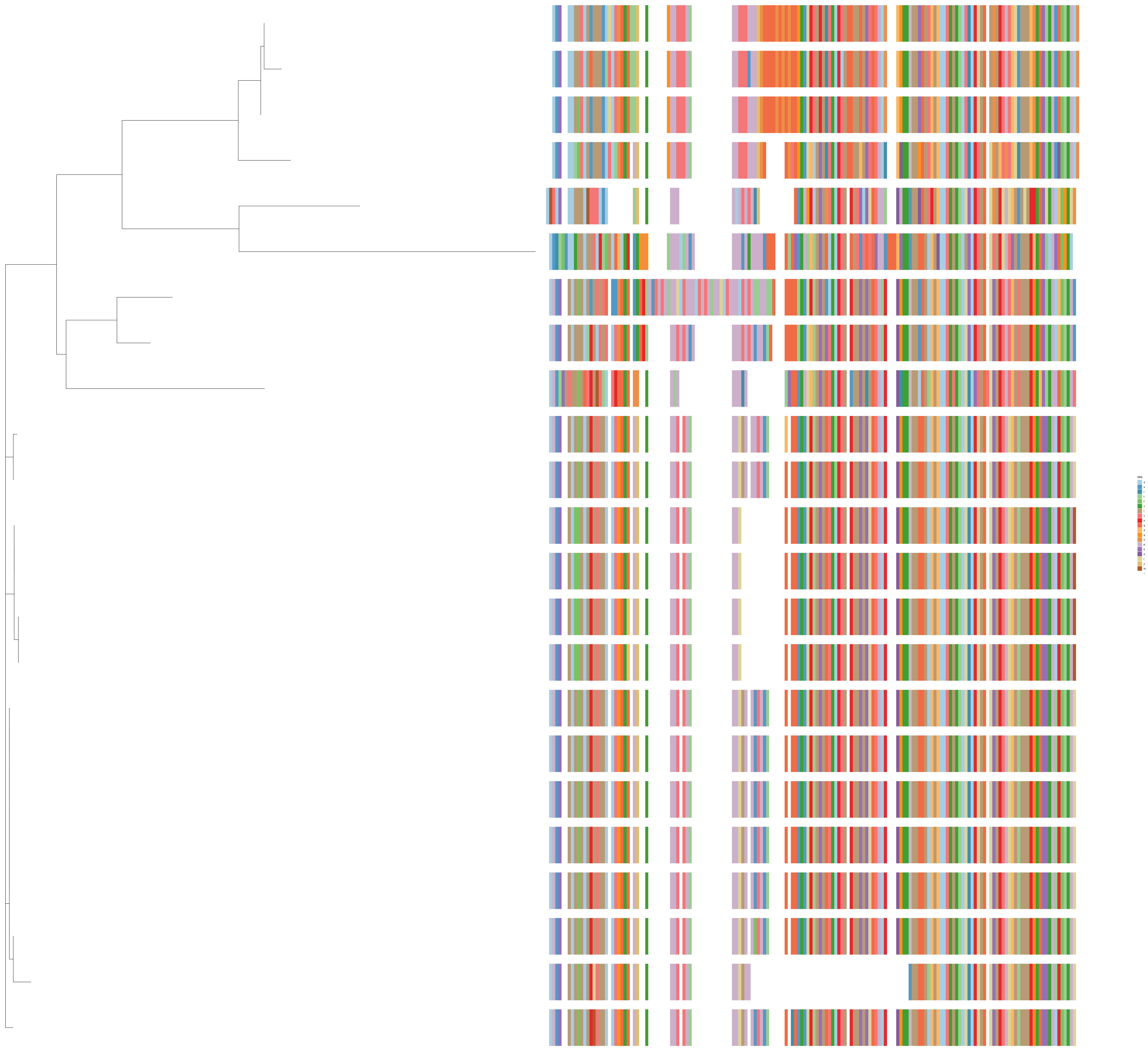

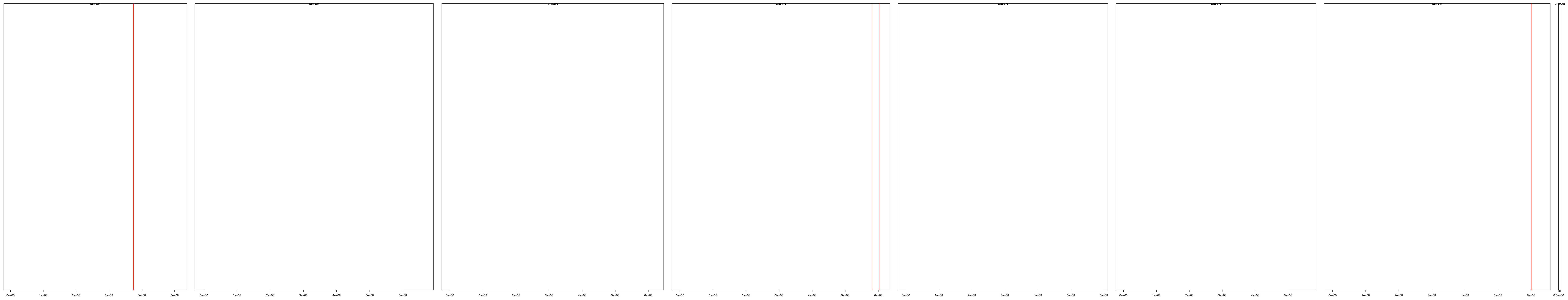

| | description | id | family | seqname | start | end |
|---|---|---|---|---|---|---|
| [1,] | cl_15128 | HORVU.MOREX.r3.1HG0056070.1 | HORVU.MOREX.r3.1HG0056070 | chr1H | 374370543 | 374372977 |
| [2,] | cl_15128 | HORVU.MOREX.r3.1HG0056110.1 | HORVU.MOREX.r3.1HG0056110 | chr1H | 374419433 | 374422181 |
| [3,] | cl_15128 | HORVU.MOREX.r3.1HG0056130.1 | HORVU.MOREX.r3.1HG0056130 | chr1H | 374527496 | 374529930 |
| [4,] | cl_15128 | HORVU.MOREX.r3.1HG0056150.1 | HORVU.MOREX.r3.1HG0056150 | chr1H | 374660896 | 374663513 |
| [5,] | cl_15128 | HORVU.MOREX.r3.4HG0407950.1 | HORVU.MOREX.r3.4HG0407950 | chr4H | 580898656 | 580901317 |
| [6,] | cl_15128 | HORVU.MOREX.r3.4HG0415490.1 | HORVU.MOREX.r3.4HG0415490 | chr4H | 602227319 | 602230180 |
| [7,] | cl_15128 | HORVU.MOREX.r3.4HG0415520.1 | HORVU.MOREX.r3.4HG0415520 | chr4H | 602280811 | 602283612 |
| [8,] | cl_15128 | HORVU.MOREX.r3.4HG0415530.1 | HORVU.MOREX.r3.4HG0415530 | chr4H | 602299174 | 602301928 |
| [9,] | cl_15128 | HORVU.MOREX.r3.4HG0415570.1 | HORVU.MOREX.r3.4HG0415570 | chr4H | 602415817 | 602418567 |
| [10,] | cl_15128 | HORVU.MOREX.r3.7HG0738400.1 | HORVU.MOREX.r3.7HG0738400 | chr7H | 599817370 | 599820074 |
| [11,] | cl_15128 | HORVU.MOREX.r3.7HG0738410.1 | HORVU.MOREX.r3.7HG0738410 | chr7H | 599829527 | 599832234 |
| [12,] | cl_15128 | HORVU.MOREX.r3.7HG0738430.1 | HORVU.MOREX.r3.7HG0738430 | chr7H | 599946631 | 599949332 |
| [13,] | cl_15128 | HORVU.MOREX.r3.7HG0738440.1 | HORVU.MOREX.r3.7HG0738440 | chr7H | 599983351 | 599985975 |
| [14,] | cl_15128 | HORVU.MOREX.r3.7HG0738450.1 | HORVU.MOREX.r3.7HG0738450 | chr7H | 599992534 | 599995301 |
| [15,] | cl_15128 | HORVU.MOREX.r3.7HG0738460.1 | HORVU.MOREX.r3.7HG0738460 | chr7H | 600026812 | 600029210 |
| [16,] | cl_15128 | HORVU.MOREX.r3.7HG0738470.1 | HORVU.MOREX.r3.7HG0738470 | chr7H | 600057515 | 600059889 |
| [17,] | cl_15128 | HORVU.MOREX.r3.7HG0738480.1 | HORVU.MOREX.r3.7HG0738480 | chr7H | 600083430 | 600085793 |
| [18,] | cl_15128 | HORVU.MOREX.r3.7HG0738490.1 | HORVU.MOREX.r3.7HG0738490 | chr7H | 600091330 | 600093704 |
| [19,] | cl_15128 | HORVU.MOREX.r3.7HG0738500.1 | HORVU.MOREX.r3.7HG0738500 | chr7H | 600116956 | 600119354 |
| [20,] | cl_15128 | HORVU.MOREX.r3.7HG0738510.1 | HORVU.MOREX.r3.7HG0738510 | chr7H | 600153608 | 600156006 |
| [21,] | cl_15128 | HORVU.MOREX.r3.7HG0738520.1 | HORVU.MOREX.r3.7HG0738520 | chr7H | 600162950 | 600165324 |
| [22,] | cl_15128 | HORVU.MOREX.r3.7HG0738530.1 | HORVU.MOREX.r3.7HG0738530 | chr7H | 600189502 | 600192080 |
| [23,] | cl_15128 | HORVU.MOREX.r3.7HG0738560.1 | HORVU.MOREX.r3.7HG0738560 | chr7H | 600343728 | 600346487 |

cl_8856

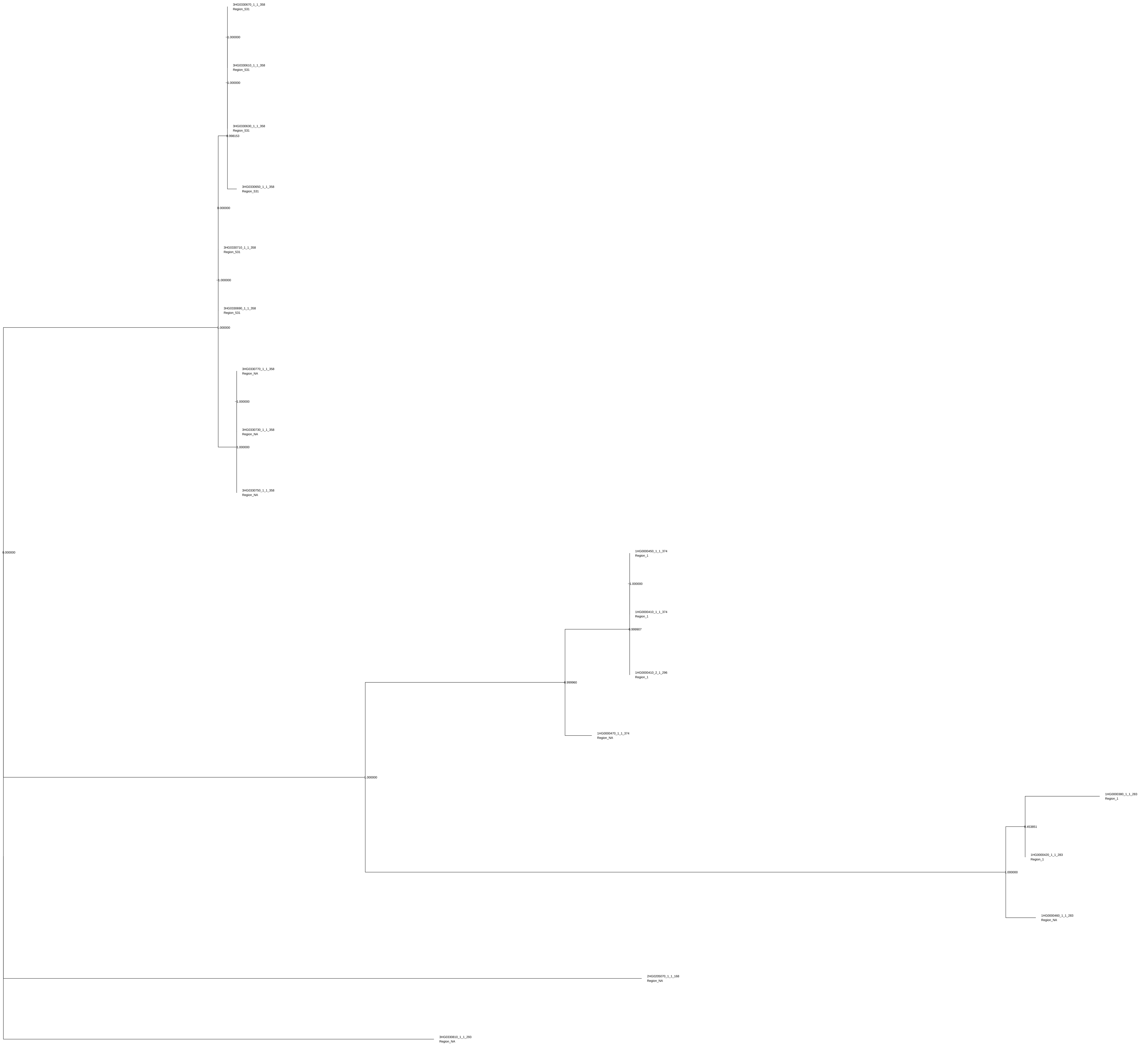
3HG0330670_1_1_358
Region_531
1.000000
3HG0330610_1_1_358
Region_531
1.000000
3HG0330630_1_1_358
Region_531
0.998153
3HG0330650_1_1_358
Region_531
0.000000
3HG0330710_1_1_358
Region_531
1.000000
3HG0330690_1_1_358
Region_531
1.000000
3HG0330770_1_1_358
Region_NA
1.000000
3HG0330730_1_1_358
Region_NA
1.000000
3HG0330750_1_1_358
Region_NA
0.000000
1HG0000450_1_1_374
Region_1
1.000000
1HG0000410_1_1_374
Region_1
0.999907
1HG0000410_2_1_296
Region_1
0.999960
1HG0000470_1_1_374
Region_NA
1.000000
1HG0000380_1_1_283
Region_1
0.453851
1HG0000420_1_1_283
Region_1
1.000000
1HG0000460_1_1_283
Region_NA
2HG0205070_1_1_168
Region_NA
3HG0330810_1_1_293
Region_NA

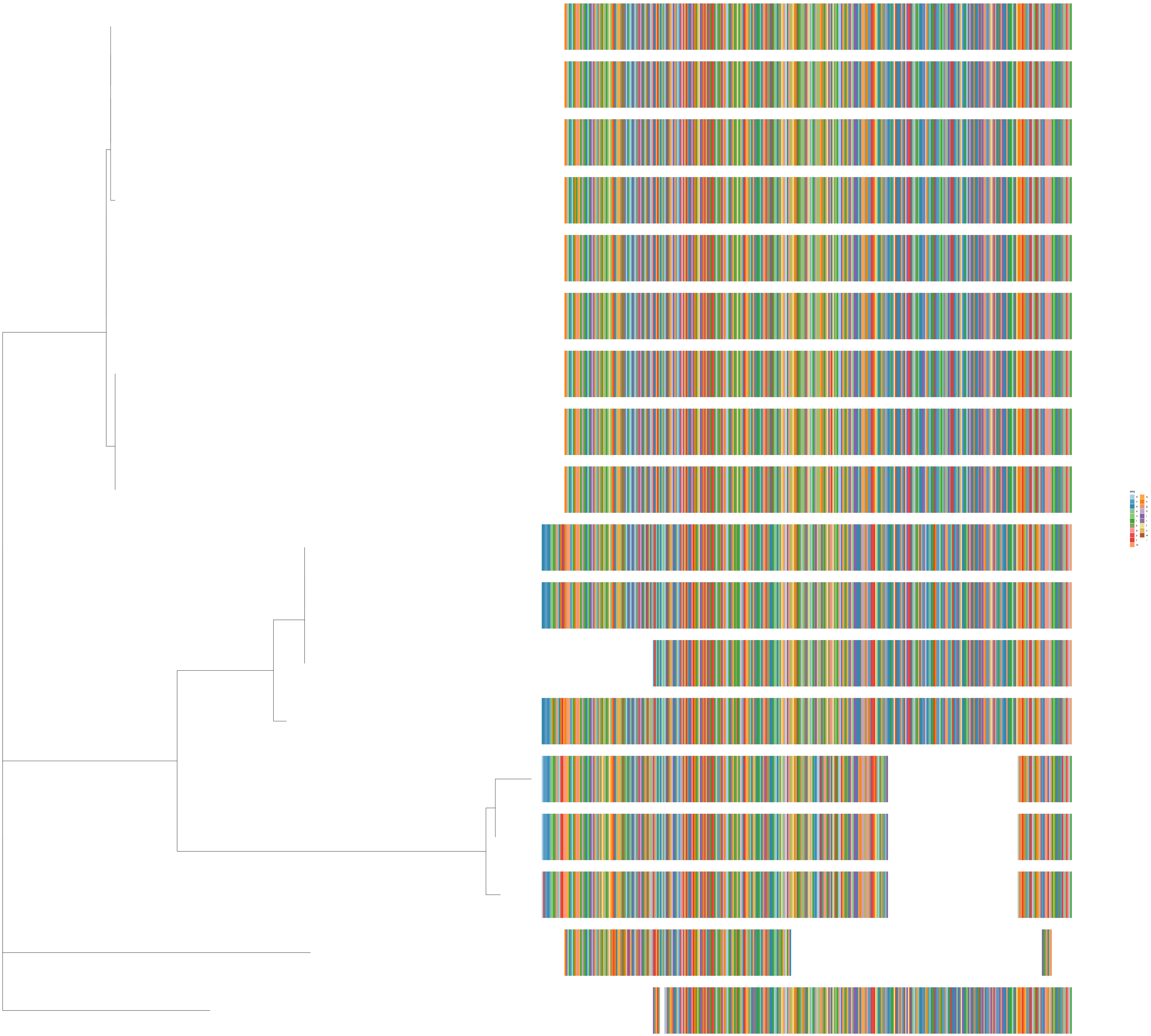
seq
d
v
a
e
n
l
k
p
y
t
m
q
s
g
h
i
r
f
c
w
−

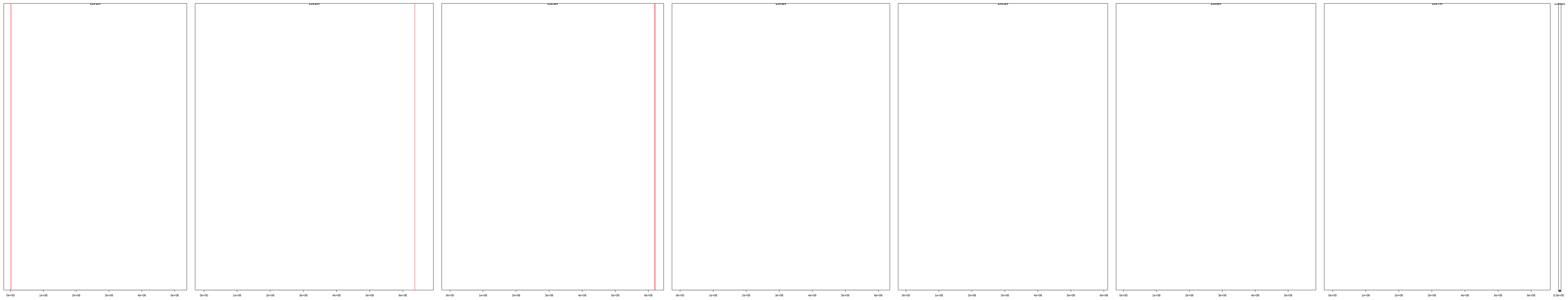

| | description | id | family | seqname | start | end |
|---|---|---|---|---|---|---|
| **[1,]** | cl_8856 | HORVU.MOREX.r3.1HG0000380.1 | HORVU.MOREX.r3.1HG0000380 | chr1H | 1179899 | 1186074 |
| **[2,]** | cl_8856 | HORVU.MOREX.r3.1HG0000410.1 | HORVU.MOREX.r3.1HG0000410 | chr1H | 1307204 | 1311237 |
| **[3,]** | cl_8856 | HORVU.MOREX.r3.1HG0000410.2 | HORVU.MOREX.r3.1HG0000410 | chr1H | 1307650 | 1311237 |
| **[4,]** | cl_8856 | HORVU.MOREX.r3.1HG0000420.1 | HORVU.MOREX.r3.1HG0000420 | chr1H | 1316064 | 1322214 |
| **[5,]** | cl_8856 | HORVU.MOREX.r3.1HG0000450.1 | HORVU.MOREX.r3.1HG0000450 | chr1H | 1443203 | 1446966 |
| **[6,]** | cl_8856 | HORVU.MOREX.r3.1HG0000460.1 | HORVU.MOREX.r3.1HG0000460 | chr1H | 1452026 | 1458443 |
| **[7,]** | cl_8856 | HORVU.MOREX.r3.1HG0000470.1 | HORVU.MOREX.r3.1HG0000470 | chr1H | 1473564 | 1477642 |
| **[8,]** | cl_8856 | HORVU.MOREX.r3.2HG0205070.1 | HORVU.MOREX.r3.2HG0205070 | chr2H | 636226973 | 636229476 |
| **[9,]** | cl_8856 | HORVU.MOREX.r3.3HG0330610.1 | HORVU.MOREX.r3.3HG0330610 | chr3H | 619607558 | 619612517 |
| **[10,]** | cl_8856 | HORVU.MOREX.r3.3HG0330630.1 | HORVU.MOREX.r3.3HG0330630 | chr3H | 619631330 | 619636319 |
| **[11,]** | cl_8856 | HORVU.MOREX.r3.3HG0330650.1 | HORVU.MOREX.r3.3HG0330650 | chr3H | 619702159 | 619706333 |
| **[12,]** | cl_8856 | HORVU.MOREX.r3.3HG0330670.1 | HORVU.MOREX.r3.3HG0330670 | chr3H | 619725949 | 619730123 |
| **[13,]** | cl_8856 | HORVU.MOREX.r3.3HG0330690.1 | HORVU.MOREX.r3.3HG0330690 | chr3H | 619750607 | 619755581 |
| **[14,]** | cl_8856 | HORVU.MOREX.r3.3HG0330710.1 | HORVU.MOREX.r3.3HG0330710 | chr3H | 619773752 | 619777926 |
| **[15,]** | cl_8856 | HORVU.MOREX.r3.3HG0330730.1 | HORVU.MOREX.r3.3HG0330730 | chr3H | 619795753 | 619799927 |
| **[16,]** | cl_8856 | HORVU.MOREX.r3.3HG0330750.1 | HORVU.MOREX.r3.3HG0330750 | chr3H | 619817763 | 619821937 |
| **[17,]** | cl_8856 | HORVU.MOREX.r3.3HG0330770.1 | HORVU.MOREX.r3.3HG0330770 | chr3H | 619839771 | 619843945 |
| **[18,]** | cl_8856 | HORVU.MOREX.r3.3HG0330810.1 | HORVU.MOREX.r3.3HG0330810 | chr3H | 620343985 | 620347727 |

cl_6074

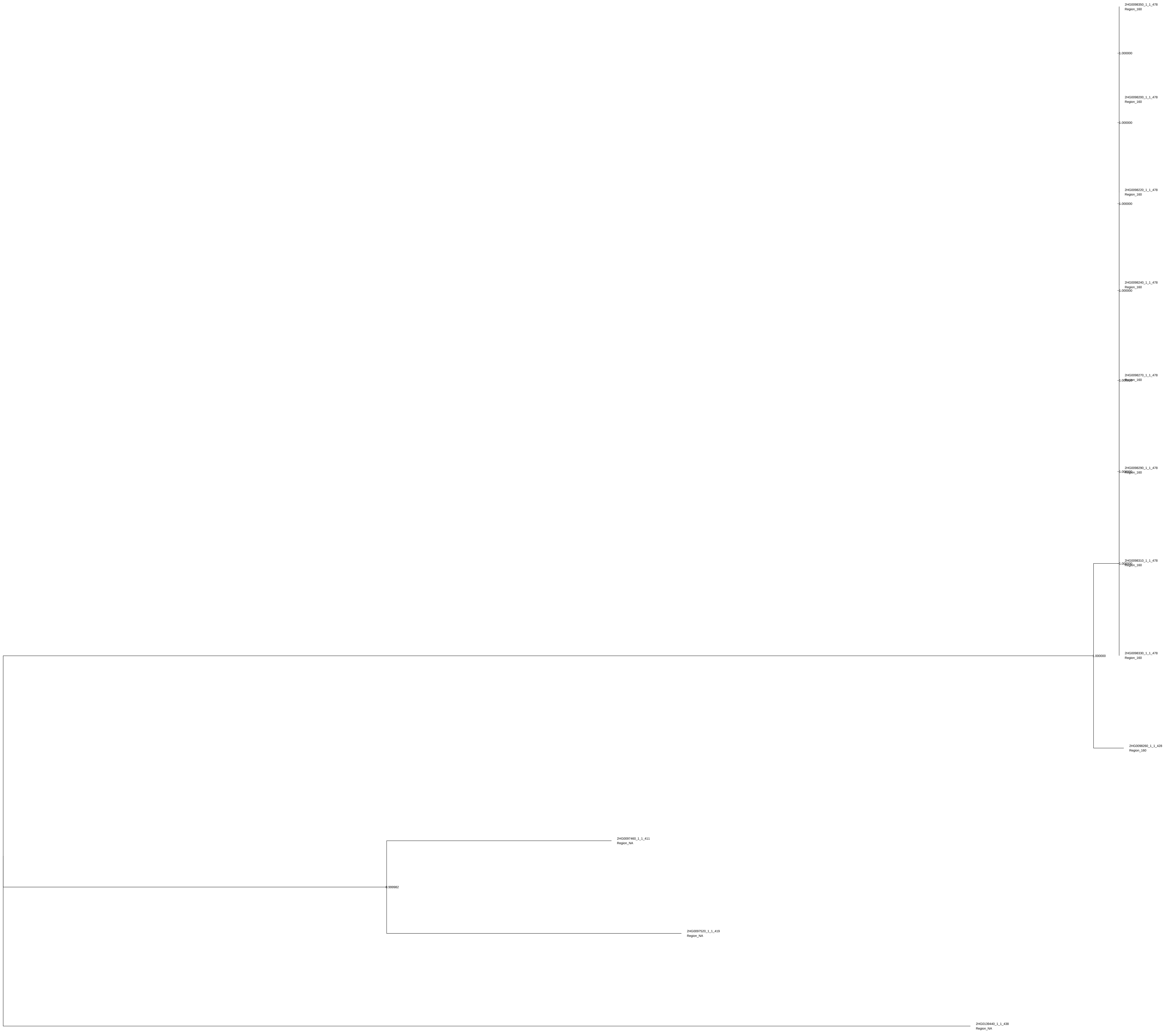
2HG0098350_1_1_478
Region_160
1.000000
2HG0098200_1_1_478
Region_160
1.000000
2HG0098220_1_1_478
Region_160
1.000000
2HG0098240_1_1_478
Region_160
1.000000
2HG0098270_1_1_478
Region_160
1.000000
2HG0098290_1_1_478
Region_160
1.000000
2HG0098310_1_1_478
Region_160
1.000000
2HG0098330_1_1_478
Region_160
1.000000
2HG0098260_1_1_428
Region_160
2HG0097460_1_1_411
Region_NA
0.999982
2HG0097520_1_1_419
Region_NA
2HG0139440_1_1_438
Region_NA

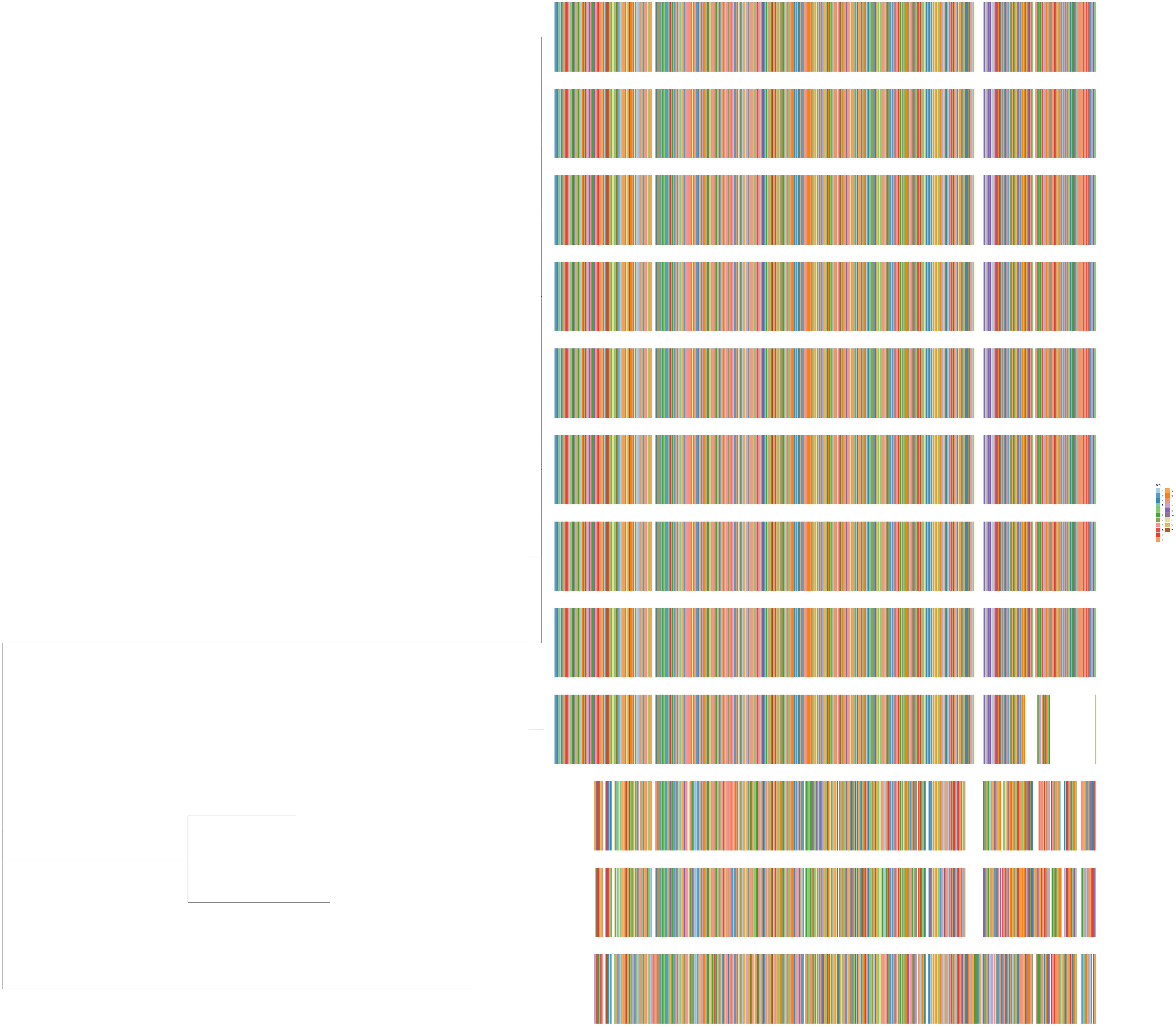
seq
l
v
n
f
d
t
r
a
c
k
i
p
g
s
h
q
m
e
y
w
−

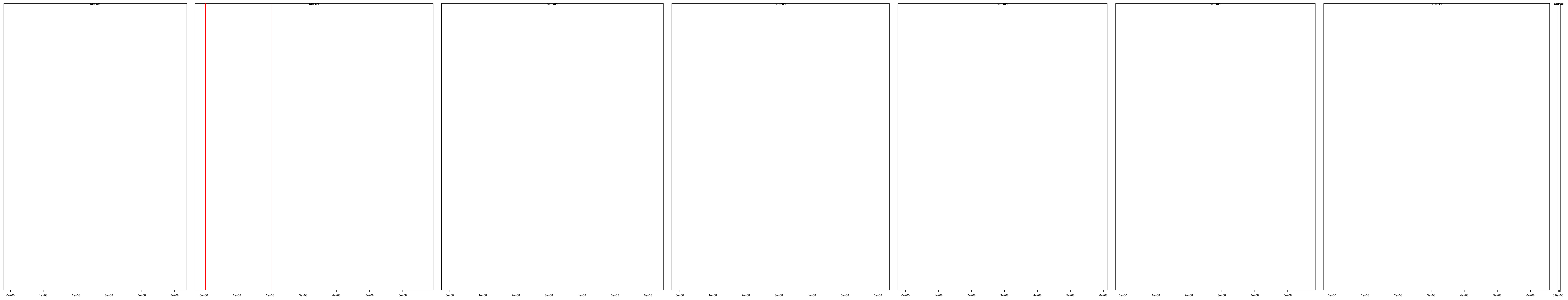

| | description | id | family | seqname | start | end |
|---|---|---|---|---|---|---|
| [1,] | cl_6074 | HORVU.MOREX.r3.2HG0097460.1 | HORVU.MOREX.r3.2HG0097460 | chr2H | 4737593 | 4741254 |
| [2,] | cl_6074 | HORVU.MOREX.r3.2HG0097520.1 | HORVU.MOREX.r3.2HG0097520 | chr2H | 5019971 | 5023227 |
| [3,] | cl_6074 | HORVU.MOREX.r3.2HG0098200.1 | HORVU.MOREX.r3.2HG0098200 | chr2H | 5898893 | 5902326 |
| [4,] | cl_6074 | HORVU.MOREX.r3.2HG0098220.1 | HORVU.MOREX.r3.2HG0098220 | chr2H | 5929561 | 5932994 |
| [5,] | cl_6074 | HORVU.MOREX.r3.2HG0098240.1 | HORVU.MOREX.r3.2HG0098240 | chr2H | 5960224 | 5963657 |
| [6,] | cl_6074 | HORVU.MOREX.r3.2HG0098260.1 | HORVU.MOREX.r3.2HG0098260 | chr2H | 5990889 | 5994172 |
| [7,] | cl_6074 | HORVU.MOREX.r3.2HG0098270.1 | HORVU.MOREX.r3.2HG0098270 | chr2H | 6004805 | 6008238 |
| [8,] | cl_6074 | HORVU.MOREX.r3.2HG0098290.1 | HORVU.MOREX.r3.2HG0098290 | chr2H | 6035467 | 6038900 |
| [9,] | cl_6074 | HORVU.MOREX.r3.2HG0098310.1 | HORVU.MOREX.r3.2HG0098310 | chr2H | 6066094 | 6069527 |
| [10,] | cl_6074 | HORVU.MOREX.r3.2HG0098330.1 | HORVU.MOREX.r3.2HG0098330 | chr2H | 6096729 | 6100162 |
| [11,] | cl_6074 | HORVU.MOREX.r3.2HG0098350.1 | HORVU.MOREX.r3.2HG0098350 | chr2H | 6127386 | 6130819 |
| [12,] | cl_6074 | HORVU.MOREX.r3.2HG0139440.1 | HORVU.MOREX.r3.2HG0139440 | chr2H | 202872713 | 202876095 |

cl_2850

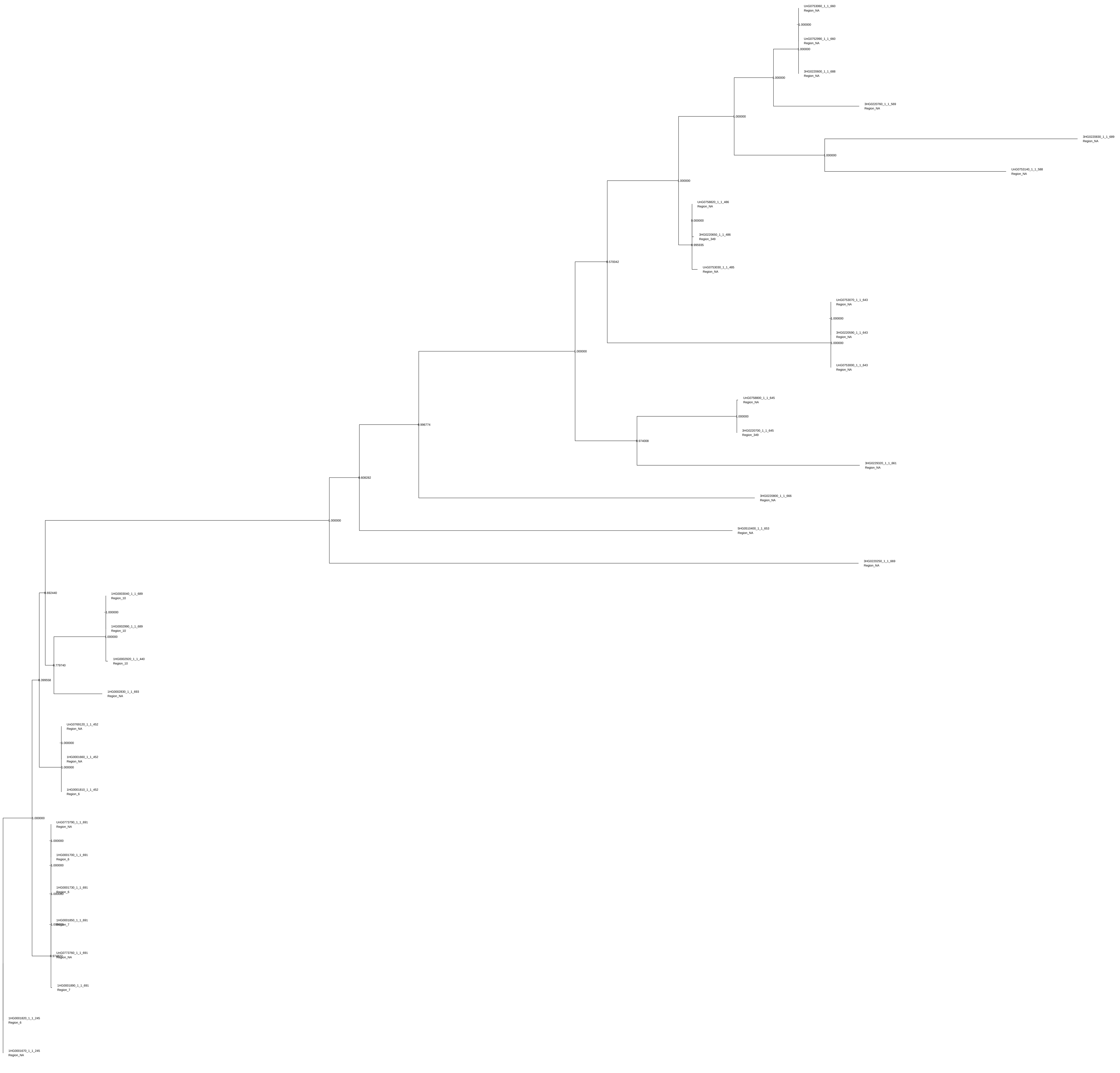

UnG0753060_1_1_660
Region_NA
1.000000
UnG0752990_1_1_660
Region_NA
1.000000
3HG0220600_1_1_688
Region_NA
1.000000
3HG0220760_1_1_569
Region_NA
1.000000
3HG0220830_1_1_689
Region_NA
1.000000
UnG0753140_1_1_588
Region_NA
1.000000
UnG0758820_1_1_486
Region_NA
0.000000
3HG0220650_1_1_486
Region_349
0.995935
UnG0753030_1_1_485
Region_NA
0.570042
UnG0753070_1_1_643
Region_NA
1.000000
3HG0220590_1_1_643
Region_NA
1.000000
UnG0753000_1_1_643
Region_NA
1.000000
UnG0758800_1_1_645
Region_NA
1.000000
3HG0220700_1_1_645
Region_349
0.974008
3HG0229320_1_1_661
Region_NA
0.996774
3HG0220800_1_1_666
Region_NA
0.608282
5HG0510400_1_1_653
Region_NA
1.000000
3HG0220250_1_1_669
Region_NA
0.692440
1HG0003040_1_1_689
Region_10
1.000000
1HG0002990_1_1_689
Region_10
1.000000
1HG0002920_1_1_440
Region_10
0.779740
0.399558
1HG0002830_1_1_893
Region_NA
UnG0769120_1_1_452
Region_NA
1.000000
1HG0001660_1_1_452
Region_NA
1.000000
1HG0001810_1_1_452
Region_6
1.000000
UnG0773790_1_1_691
Region_NA
1.000000
1HG0001700_1_1_691
Region_6
1.000000
1HG0001730_1_1_691
Region_6
1.000000
1HG0001850_1_1_691
Region_7
1.000000
UnG0773760_1_1_691
Region_NA
0.978022
1HG0001890_1_1_691
Region_7
1HG0001820_1_1_245
Region_6
1HG0001670_1_1_245
Region_NA

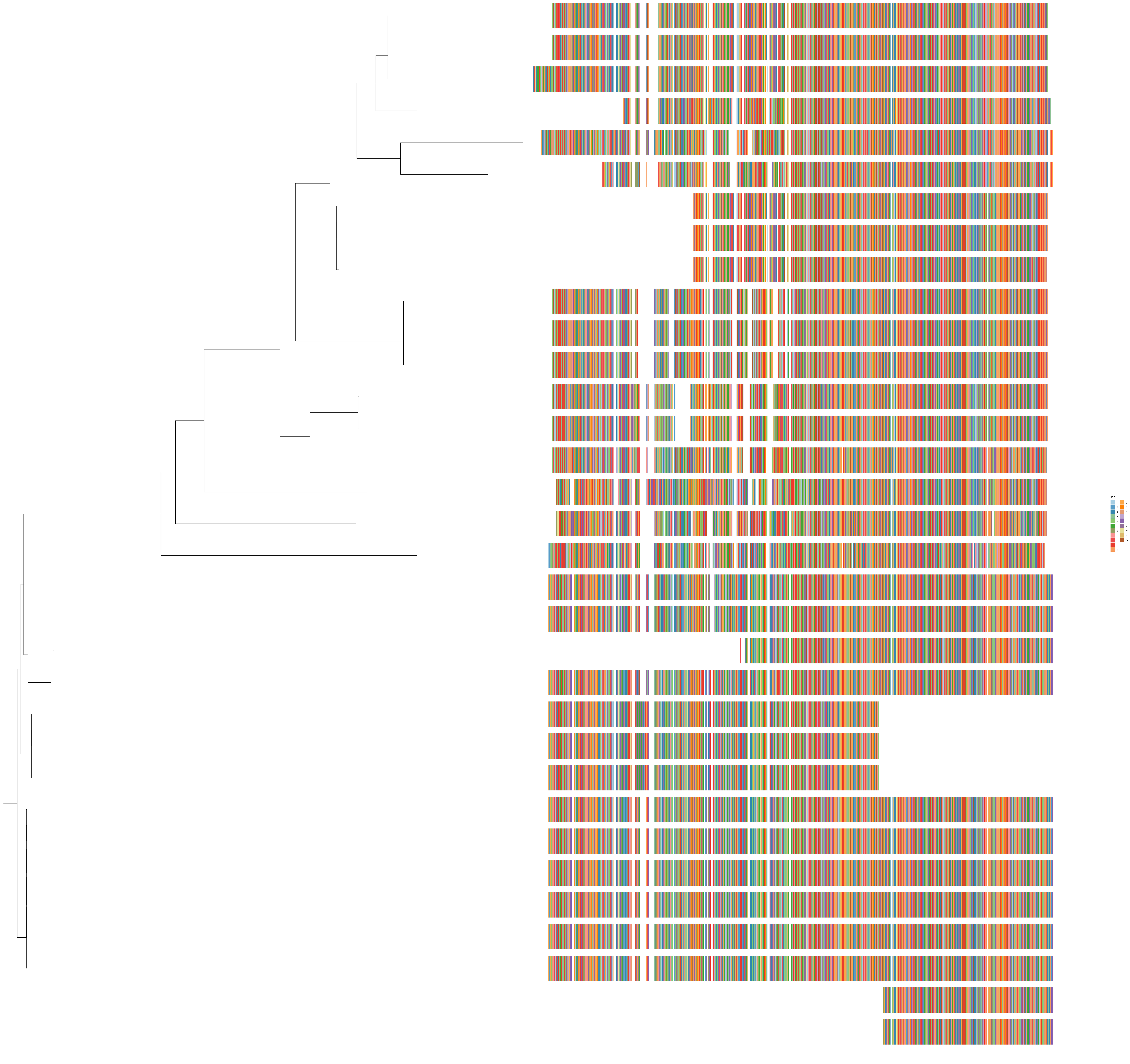
seq
t
n
s
v
a
f
p
r
i
l
e
g
c
h
q
d
y
w
k
m
–

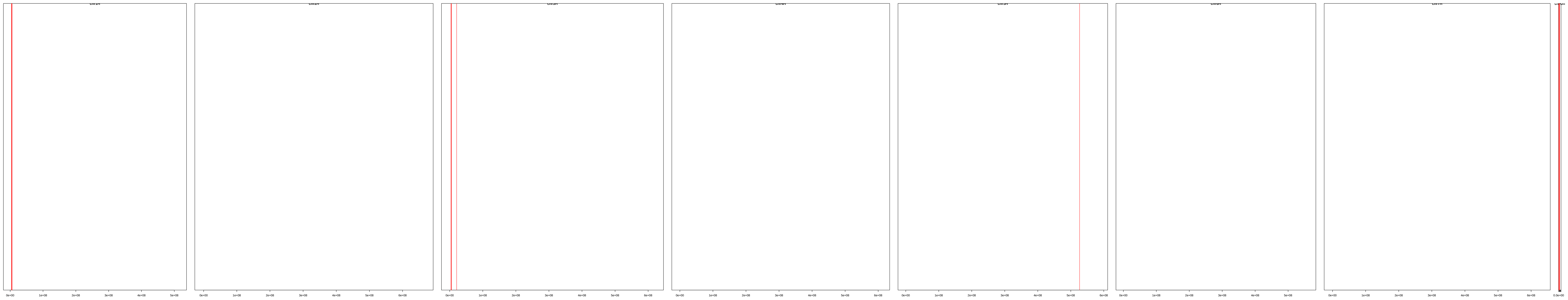

| | description | id | family | seqname | start | end |
|---|---|---|---|---|---|---|
| [1,] | cl_2850 | HORVU.MOREX.r3.1HG0001660.1 | HORVU.MOREX.r3.1HG0001660 | chr1H | 3828914 | 3834421 |
| [2,] | cl_2850 | HORVU.MOREX.r3.1HG0001670.1 | HORVU.MOREX.r3.1HG0001670 | chr1H | 3832434 | 3835168 |
| [3,] | cl_2850 | HORVU.MOREX.r3.1HG0001700.1 | HORVU.MOREX.r3.1HG0001700 | chr1H | 3917699 | 3922273 |
| [4,] | cl_2850 | HORVU.MOREX.r3.1HG0001730.1 | HORVU.MOREX.r3.1HG0001730 | chr1H | 4009580 | 4014154 |
| [5,] | cl_2850 | HORVU.MOREX.r3.1HG0001810.1 | HORVU.MOREX.r3.1HG0001810 | chr1H | 4121142 | 4126649 |
| [6,] | cl_2850 | HORVU.MOREX.r3.1HG0001820.1 | HORVU.MOREX.r3.1HG0001820 | chr1H | 4124662 | 4127396 |
| [7,] | cl_2850 | HORVU.MOREX.r3.1HG0001850.1 | HORVU.MOREX.r3.1HG0001850 | chr1H | 4209887 | 4214461 |
| [8,] | cl_2850 | HORVU.MOREX.r3.1HG0001890.1 | HORVU.MOREX.r3.1HG0001890 | chr1H | 4301769 | 4306343 |
| [9,] | cl_2850 | HORVU.MOREX.r3.1HG0002830.1 | HORVU.MOREX.r3.1HG0002830 | chr1H | 5690018 | 5694589 |
| [10,] | cl_2850 | HORVU.MOREX.r3.1HG0002920.1 | HORVU.MOREX.r3.1HG0002920 | chr1H | 5753051 | 5756507 |
| [11,] | cl_2850 | HORVU.MOREX.r3.1HG0002990.1 | HORVU.MOREX.r3.1HG0002990 | chr1H | 5793787 | 5798378 |
| [12,] | cl_2850 | HORVU.MOREX.r3.1HG0003040.1 | HORVU.MOREX.r3.1HG0003040 | chr1H | 5835660 | 5840251 |
| [13,] | cl_2850 | HORVU.MOREX.r3.3HG0220250.1 | HORVU.MOREX.r3.3HG0220250 | chr3H | 3939823 | 3944209 |
| [14,] | cl_2850 | HORVU.MOREX.r3.3HG0220590.1 | HORVU.MOREX.r3.3HG0220590 | chr3H | 4502870 | 4507013 |
| [15,] | cl_2850 | HORVU.MOREX.r3.3HG0220600.1 | HORVU.MOREX.r3.3HG0220600 | chr3H | 4531838 | 4536086 |
| [16,] | cl_2850 | HORVU.MOREX.r3.3HG0220650.1 | HORVU.MOREX.r3.3HG0220650 | chr3H | 4724779 | 4779093 |
| [17,] | cl_2850 | HORVU.MOREX.r3.3HG0220700.1 | HORVU.MOREX.r3.3HG0220700 | chr3H | 4895078 | 4899266 |
| [18,] | cl_2850 | HORVU.MOREX.r3.3HG0220760.1 | HORVU.MOREX.r3.3HG0220760 | chr3H | 4992912 | 4996828 |
| [19,] | cl_2850 | HORVU.MOREX.r3.3HG0220800.1 | HORVU.MOREX.r3.3HG0220800 | chr3H | 5070444 | 5074645 |
| [20,] | cl_2850 | HORVU.MOREX.r3.3HG0220830.1 | HORVU.MOREX.r3.3HG0220830 | chr3H | 5090724 | 5095118 |
| [21,] | cl_2850 | HORVU.MOREX.r3.3HG0229320.1 | HORVU.MOREX.r3.3HG0229320 | chr3H | 21068579 | 21072895 |
| [22,] | cl_2850 | HORVU.MOREX.r3.5HG0510400.1 | HORVU.MOREX.r3.5HG0510400 | chr5H | 526707938 | 526712065 |
| [23,] | cl_2850 | HORVU.MOREX.r3.UnG0752990.1 | HORVU.MOREX.r3.UnG0752990 | chrUn | 104767 | 108931 |
| [24,] | cl_2850 | HORVU.MOREX.r3.UnG0753000.1 | HORVU.MOREX.r3.UnG0753000 | chrUn | 133837 | 137980 |
| [25,] | cl_2850 | HORVU.MOREX.r3.UnG0753030.1 | HORVU.MOREX.r3.UnG0753030 | chrUn | 171928 | 175567 |
| [26,] | cl_2850 | HORVU.MOREX.r3.UnG0753060.1 | HORVU.MOREX.r3.UnG0753060 | chrUn | 364201 | 368365 |
| [27,] | cl_2850 | HORVU.MOREX.r3.UnG0753070.1 | HORVU.MOREX.r3.UnG0753070 | chrUn | 386176 | 390319 |
| [28,] | cl_2850 | HORVU.MOREX.r3.UnG0753140.1 | HORVU.MOREX.r3.UnG0753140 | chrUn | 472472 | 481747 |
| [29,] | cl_2850 | HORVU.MOREX.r3.UnG0758800.1 | HORVU.MOREX.r3.UnG0758800 | chrUn | 5047794 | 5051982 |
| [30,] | cl_2850 | HORVU.MOREX.r3.UnG0758820.1 | HORVU.MOREX.r3.UnG0758820 | chrUn | 5167946 | 5222243 |
| [31,] | cl_2850 | HORVU.MOREX.r3.UnG0769120.1 | HORVU.MOREX.r3.UnG0769120 | chrUn | 10517273 | 10522780 |
| [32,] | cl_2850 | HORVU.MOREX.r3.UnG0773760.1 | HORVU.MOREX.r3.UnG0773760 | chrUn | 12674258 | 12678832 |
| [33,] | cl_2850 | HORVU.MOREX.r3.UnG0773790.1 | HORVU.MOREX.r3.UnG0773790 | chrUn | 12762902 | 12767476 |

cl_1274

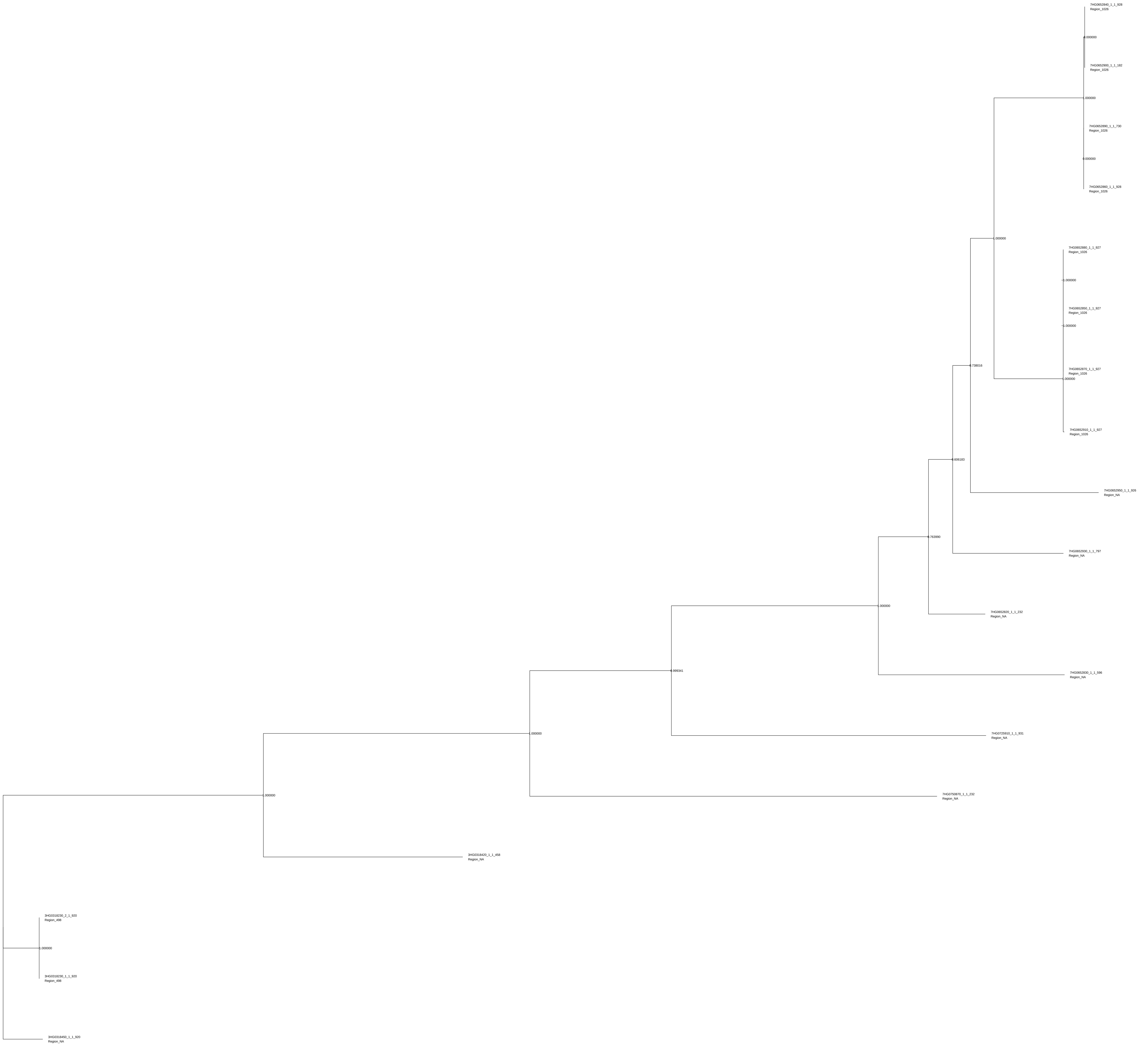

7HG0652840_1_1_928
Region_1026
0.000000
7HG0652900_1_1_182
Region_1026
1.000000
7HG0652890_1_1_730
Region_1026
0.000000
7HG0652860_1_1_928
Region_1026
1.000000
7HG0652880_1_1_927
Region_1026
1.000000
7HG0652850_1_1_927
Region_1026
1.000000
0.738016
7HG0652870_1_1_927
Region_1026
1.000000
7HG0652910_1_1_927
Region_1026
0.606183
7HG0652950_1_1_926
Region_NA
0.763990
7HG0652930_1_1_797
Region_NA
1.000000
7HG0652820_1_1_232
Region_NA
0.999341
7HG0652830_1_1_596
Region_NA
1.000000
7HG0725910_1_1_931
Region_NA
1.000000
7HG0750870_1_1_232
Region_NA
3HG0318420_1_1_458
Region_NA
3HG0318230_2_1_920
Region_498
1.000000
3HG0318230_1_1_920
Region_498
3HG0318450_1_1_920
Region_NA

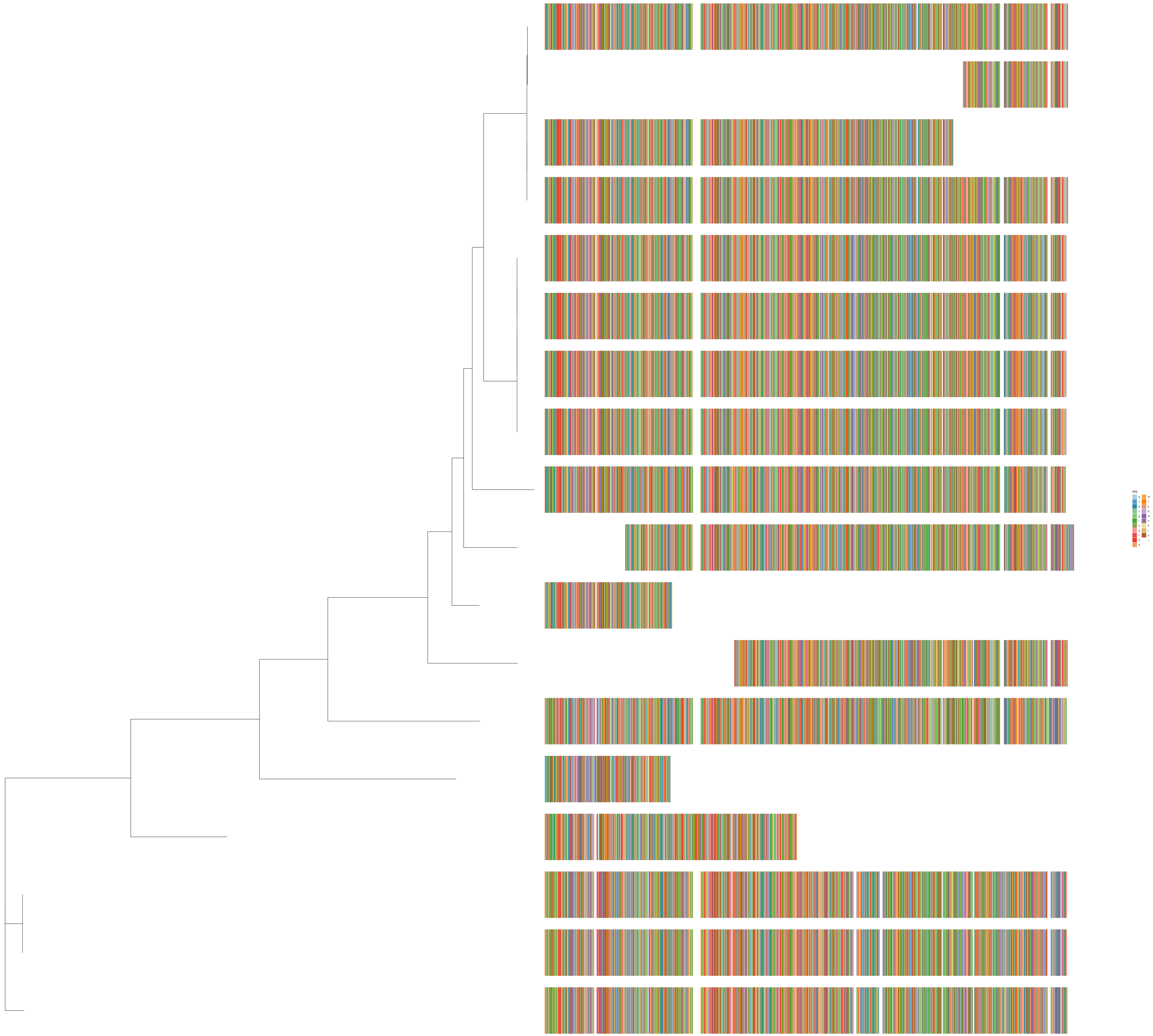
seq
q
v
a
t
g
l
s
p
r
k
e
m
f
y
d
w
n
h
i
c
–

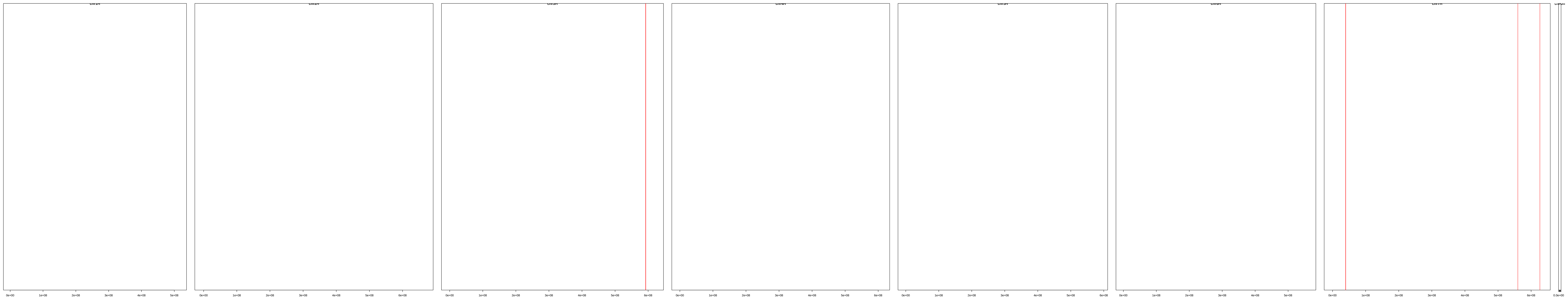

| | description | id | family | seqname | start | end |
|---|---|---|---|---|---|---|
| [1,] | cl_1274 | HORVU.MOREX.r3.3HG0318230.1 | HORVU.MOREX.r3.3HG0318230 | chr3H | 592334259 | 592340426 |
| [2,] | cl_1274 | HORVU.MOREX.r3.3HG0318230.2 | HORVU.MOREX.r3.3HG0318230 | chr3H | 592334261 | 592340426 |
| [3,] | cl_1274 | HORVU.MOREX.r3.3HG0318420.1 | HORVU.MOREX.r3.3HG0318420 | chr3H | 592700827 | 592704468 |
| [4,] | cl_1274 | HORVU.MOREX.r3.3HG0318450.1 | HORVU.MOREX.r3.3HG0318450 | chr3H | 592713399 | 592719570 |
| [5,] | cl_1274 | HORVU.MOREX.r3.7HG0652820.1 | HORVU.MOREX.r3.7HG0652820 | chr7H | 39557062 | 39559757 |
| [6,] | cl_1274 | HORVU.MOREX.r3.7HG0652830.1 | HORVU.MOREX.r3.7HG0652830 | chr7H | 39559633 | 39563420 |
| [7,] | cl_1274 | HORVU.MOREX.r3.7HG0652840.1 | HORVU.MOREX.r3.7HG0652840 | chr7H | 39581427 | 39586556 |
| [8,] | cl_1274 | HORVU.MOREX.r3.7HG0652850.1 | HORVU.MOREX.r3.7HG0652850 | chr7H | 39597025 | 39602159 |
| [9,] | cl_1274 | HORVU.MOREX.r3.7HG0652860.1 | HORVU.MOREX.r3.7HG0652860 | chr7H | 39633147 | 39639517 |
| [10,] | cl_1274 | HORVU.MOREX.r3.7HG0652870.1 | HORVU.MOREX.r3.7HG0652870 | chr7H | 39649540 | 39654674 |
| [11,] | cl_1274 | HORVU.MOREX.r3.7HG0652880.1 | HORVU.MOREX.r3.7HG0652880 | chr7H | 39664808 | 39669942 |
| [12,] | cl_1274 | HORVU.MOREX.r3.7HG0652890.1 | HORVU.MOREX.r3.7HG0652890 | chr7H | 39701728 | 39706263 |
| [13,] | cl_1274 | HORVU.MOREX.r3.7HG0652900.1 | HORVU.MOREX.r3.7HG0652900 | chr7H | 39704312 | 39706857 |
| [14,] | cl_1274 | HORVU.MOREX.r3.7HG0652910.1 | HORVU.MOREX.r3.7HG0652910 | chr7H | 39717327 | 39722461 |
| [15,] | cl_1274 | HORVU.MOREX.r3.7HG0652930.1 | HORVU.MOREX.r3.7HG0652930 | chr7H | 39753417 | 39759885 |
| [16,] | cl_1274 | HORVU.MOREX.r3.7HG0652950.1 | HORVU.MOREX.r3.7HG0652950 | chr7H | 39782179 | 39787293 |
| [17,] | cl_1274 | HORVU.MOREX.r3.7HG0725910.1 | HORVU.MOREX.r3.7HG0725910 | chr7H | 559839856 | 559845002 |
| [18,] | cl_1274 | HORVU.MOREX.r3.7HG0750870.1 | HORVU.MOREX.r3.7HG0750870 | chr7H | 626502964 | 626505659 |

cl_16769

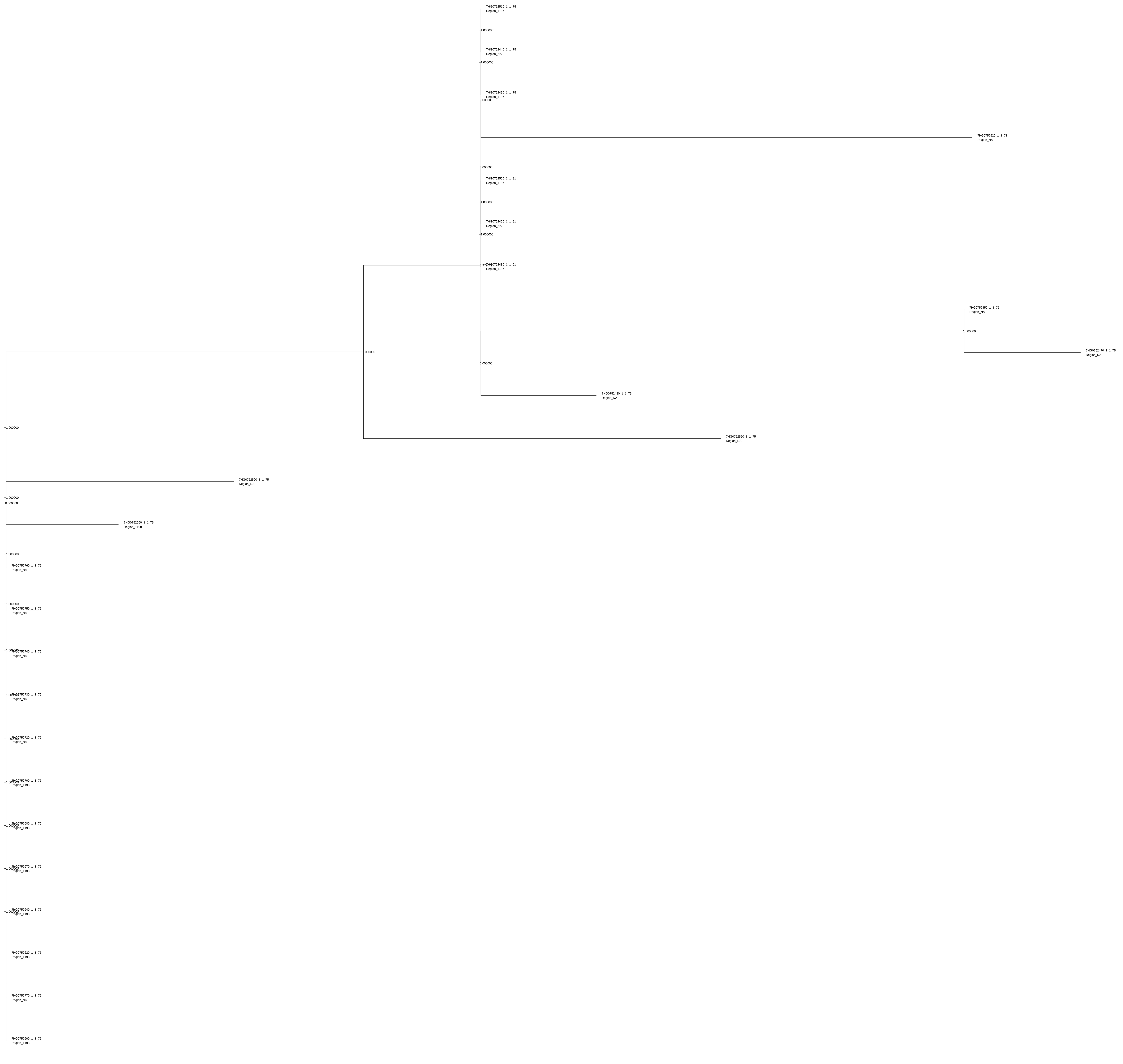
7HG0752510_1_1_75
Region_1197
1.000000
7HG0752440_1_1_75
Region_NA
1.000000
7HG0752490_1_1_75
Region_1197
0.000000
7HG0752520_1_1_71
Region_NA
0.000000
7HG0752500_1_1_91
Region_1197
1.000000
7HG0752460_1_1_91
Region_NA
1.000000
7HG0752480_1_1_91
Region_1197
7HG0752450_1_1_75
Region_NA
1.000000
7HG0752470_1_1_75
Region_NA
1.000000
0.000000
7HG0752430_1_1_75
Region_NA
1.000000
7HG0752550_1_1_75
Region_NA
7HG0752590_1_1_75
Region_NA
1.000000
0.000000
7HG0752660_1_1_75
Region_1198
1.000000
7HG0752760_1_1_75
Region_NA
1.000000
7HG0752750_1_1_75
Region_NA
7HG0752740_1_1_75
Region_NA
7HG0752730_1_1_75
Region_NA
7HG0752720_1_1_75
Region_NA
7HG0752700_1_1_75
Region_1198
7HG0752680_1_1_75
Region_1198
7HG0752670_1_1_75
Region_1198
7HG0752640_1_1_75
Region_1198
7HG0752620_1_1_75
Region_1198
7HG0752770_1_1_75
Region_NA
7HG0752600_1_1_75
Region_1198

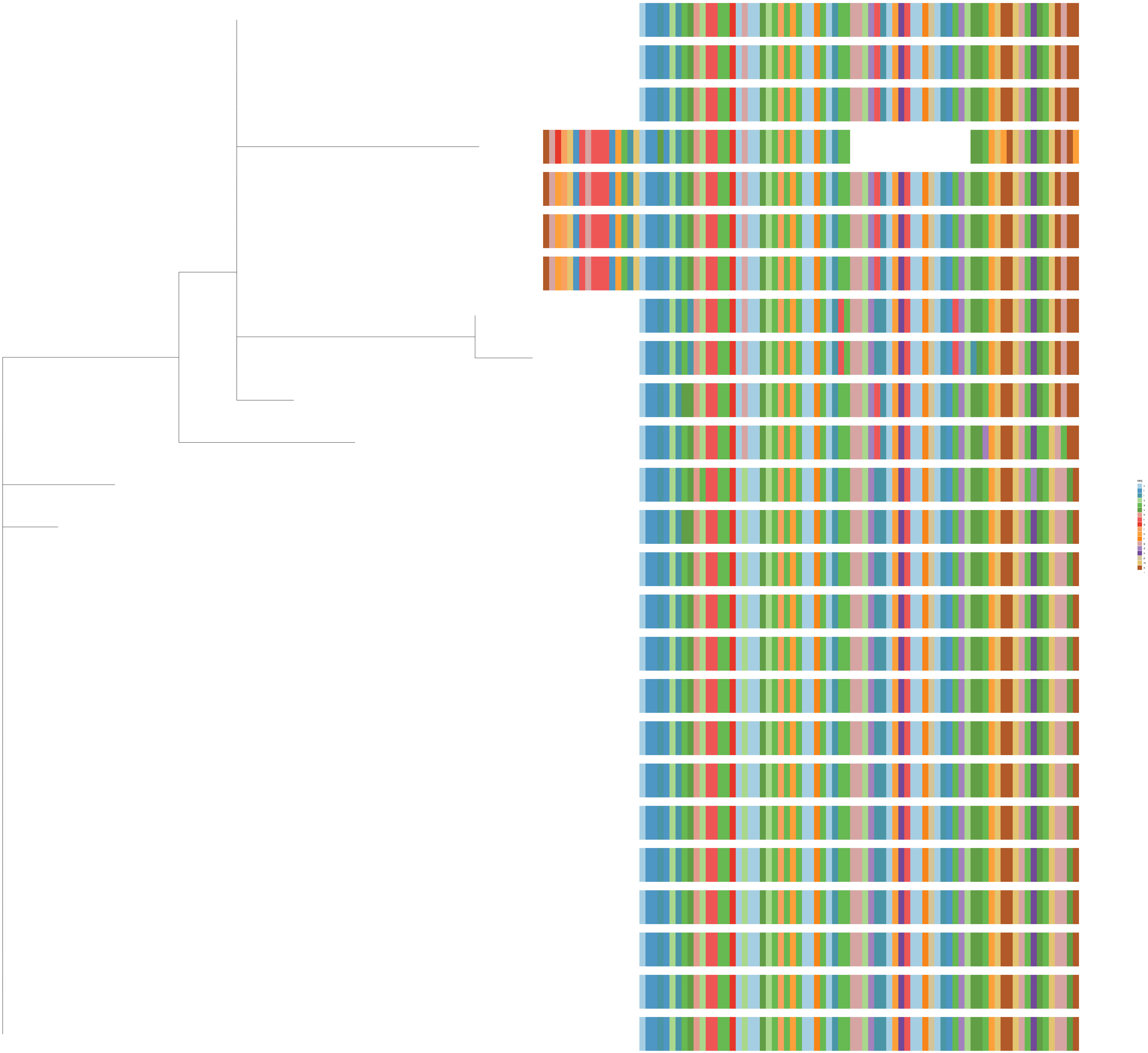

seq
c
l
i
s
a
v
h
t
q
r
k
f
g
d
n
p
m
e
–

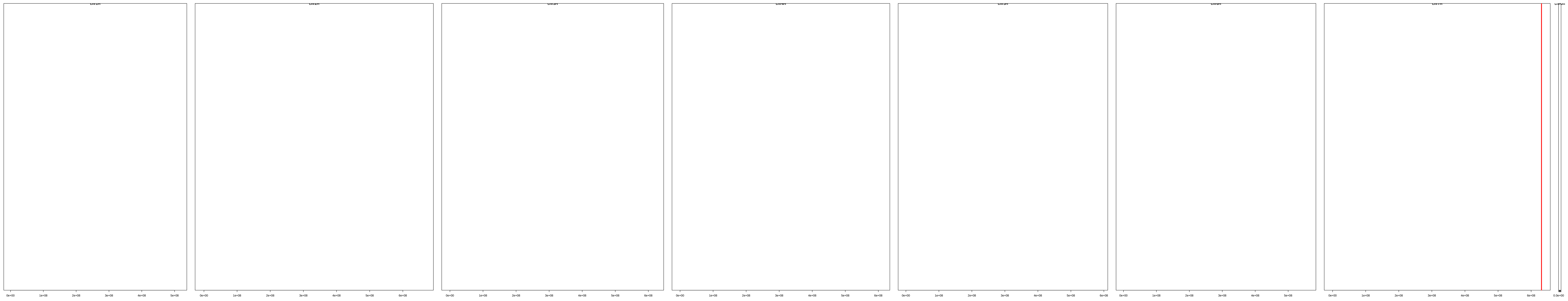

| | description | id | family | seqname | start | end |
|---|---|---|---|---|---|---|
| [1,] | cl_16769 | HORVU.MOREX.r3.7HG0752430.1 | HORVU.MOREX.r3.7HG0752430 | chr7H | 630643342 | 630645808 |
| [2,] | cl_16769 | HORVU.MOREX.r3.7HG0752440.1 | HORVU.MOREX.r3.7HG0752440 | chr7H | 630665968 | 630668434 |
| [3,] | cl_16769 | HORVU.MOREX.r3.7HG0752450.1 | HORVU.MOREX.r3.7HG0752450 | chr7H | 630689801 | 630692267 |
| [4,] | cl_16769 | HORVU.MOREX.r3.7HG0752460.1 | HORVU.MOREX.r3.7HG0752460 | chr7H | 630740534 | 630743425 |
| [5,] | cl_16769 | HORVU.MOREX.r3.7HG0752470.1 | HORVU.MOREX.r3.7HG0752470 | chr7H | 630753507 | 630755973 |
| [6,] | cl_16769 | HORVU.MOREX.r3.7HG0752480.1 | HORVU.MOREX.r3.7HG0752480 | chr7H | 630800377 | 630803255 |
| [7,] | cl_16769 | HORVU.MOREX.r3.7HG0752490.1 | HORVU.MOREX.r3.7HG0752490 | chr7H | 630822909 | 630825375 |
| [8,] | cl_16769 | HORVU.MOREX.r3.7HG0752500.1 | HORVU.MOREX.r3.7HG0752500 | chr7H | 630844741 | 630847661 |
| [9,] | cl_16769 | HORVU.MOREX.r3.7HG0752510.1 | HORVU.MOREX.r3.7HG0752510 | chr7H | 630867244 | 630869710 |
| [10,] | cl_16769 | HORVU.MOREX.r3.7HG0752520.1 | HORVU.MOREX.r3.7HG0752520 | chr7H | 630887643 | 630890157 |
| [11,] | cl_16769 | HORVU.MOREX.r3.7HG0752550.1 | HORVU.MOREX.r3.7HG0752550 | chr7H | 631061044 | 631063510 |
| [12,] | cl_16769 | HORVU.MOREX.r3.7HG0752590.1 | HORVU.MOREX.r3.7HG0752590 | chr7H | 631417631 | 631420097 |
| [13,] | cl_16769 | HORVU.MOREX.r3.7HG0752600.1 | HORVU.MOREX.r3.7HG0752600 | chr7H | 631441829 | 631444294 |
| [14,] | cl_16769 | HORVU.MOREX.r3.7HG0752620.1 | HORVU.MOREX.r3.7HG0752620 | chr7H | 631513601 | 631516067 |
| [15,] | cl_16769 | HORVU.MOREX.r3.7HG0752640.1 | HORVU.MOREX.r3.7HG0752640 | chr7H | 631541085 | 631543551 |
| [16,] | cl_16769 | HORVU.MOREX.r3.7HG0752660.1 | HORVU.MOREX.r3.7HG0752660 | chr7H | 631601042 | 631603508 |
| [17,] | cl_16769 | HORVU.MOREX.r3.7HG0752670.1 | HORVU.MOREX.r3.7HG0752670 | chr7H | 631624195 | 631626661 |
| [18,] | cl_16769 | HORVU.MOREX.r3.7HG0752680.1 | HORVU.MOREX.r3.7HG0752680 | chr7H | 631659111 | 631661577 |
| [19,] | cl_16769 | HORVU.MOREX.r3.7HG0752700.1 | HORVU.MOREX.r3.7HG0752700 | chr7H | 631741055 | 631743521 |
| [20,] | cl_16769 | HORVU.MOREX.r3.7HG0752720.1 | HORVU.MOREX.r3.7HG0752720 | chr7H | 631786560 | 631789026 |
| [21,] | cl_16769 | HORVU.MOREX.r3.7HG0752730.1 | HORVU.MOREX.r3.7HG0752730 | chr7H | 631799161 | 631801627 |
| [22,] | cl_16769 | HORVU.MOREX.r3.7HG0752740.1 | HORVU.MOREX.r3.7HG0752740 | chr7H | 631824319 | 631826785 |
| [23,] | cl_16769 | HORVU.MOREX.r3.7HG0752750.1 | HORVU.MOREX.r3.7HG0752750 | chr7H | 631862031 | 631864497 |
| [24,] | cl_16769 | HORVU.MOREX.r3.7HG0752760.1 | HORVU.MOREX.r3.7HG0752760 | chr7H | 631923583 | 631926049 |
| [25,] | cl_16769 | HORVU.MOREX.r3.7HG0752770.1 | HORVU.MOREX.r3.7HG0752770 | chr7H | 631961339 | 631963805 |

cl_1105

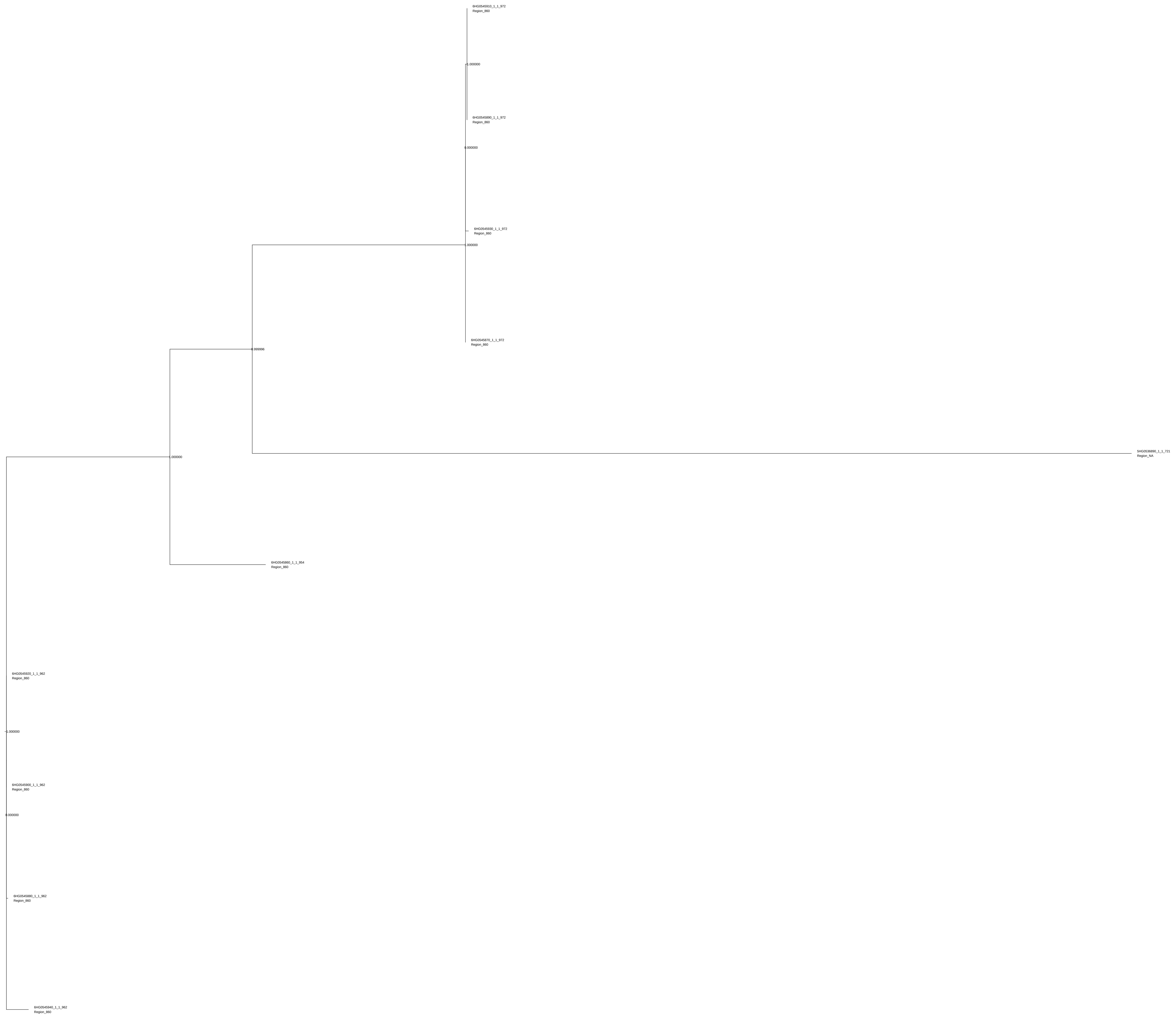
6HG0545910_1_1_972
Region_860
1.000000
6HG0545890_1_1_972
Region_860
0.000000
6HG0545930_1_1_972
Region_860
1.000000
6HG0545870_1_1_972
Region_860
0.999996
5HG0536890_1_1_721
Region_NA
1.000000
6HG0545860_1_1_954
Region_860
6HG0545920_1_1_962
Region_860
1.000000
6HG0545900_1_1_962
Region_860
0.000000
6HG0545880_1_1_962
Region_860
6HG0545940_1_1_962
Region_860

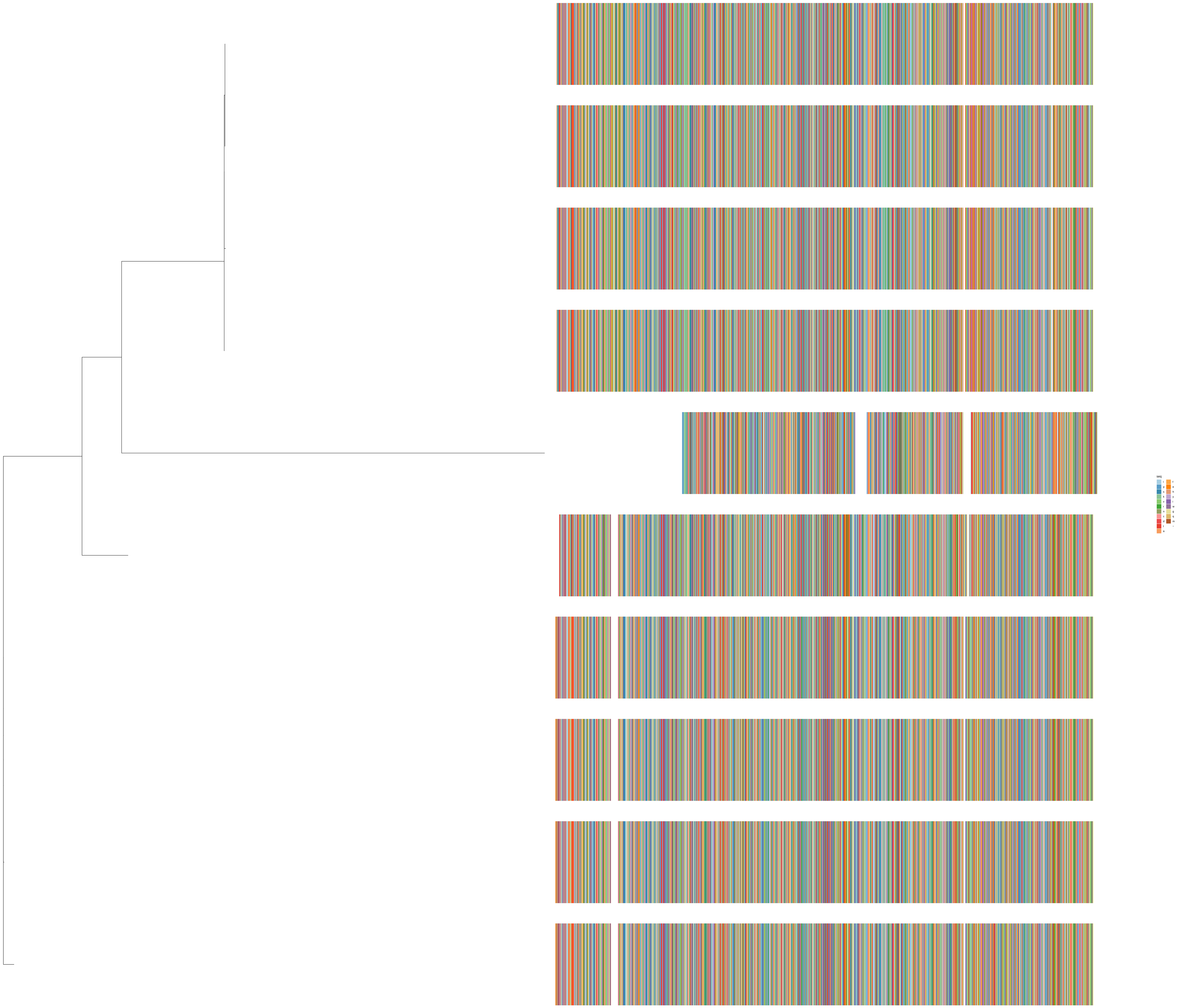
seq

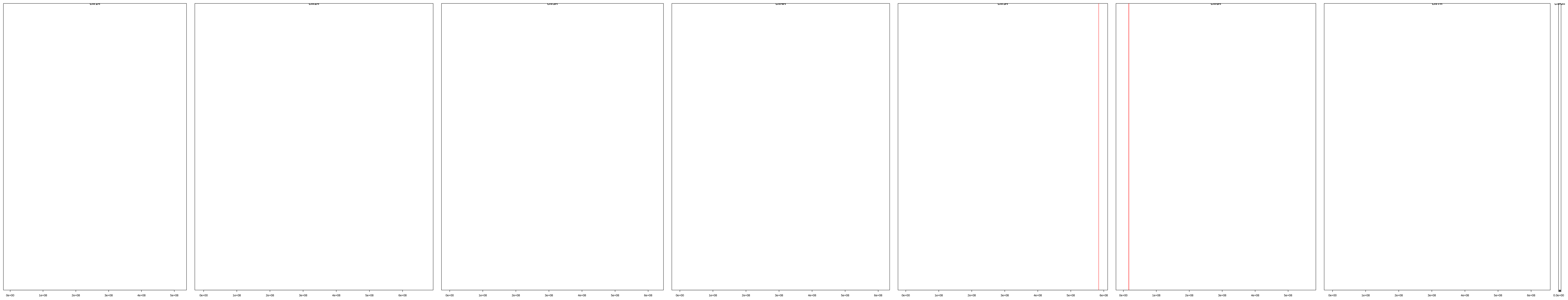

| | description | id | family | seqname | start | end |
|---|---|---|---|---|---|---|
| [1,] | cl_1105 | HORVU.MOREX.r3.5HG0536890.1 | HORVU.MOREX.r3.5HG0536890 | chr5H | 584585493 | 584589655 |
| [2,] | cl_1105 | HORVU.MOREX.r3.6HG0545860.1 | HORVU.MOREX.r3.6HG0545860 | chr6H | 16573283 | 16578144 |
| [3,] | cl_1105 | HORVU.MOREX.r3.6HG0545870.1 | HORVU.MOREX.r3.6HG0545870 | chr6H | 16580913 | 16585828 |
| [4,] | cl_1105 | HORVU.MOREX.r3.6HG0545880.1 | HORVU.MOREX.r3.6HG0545880 | chr6H | 16593223 | 16598108 |
| [5,] | cl_1105 | HORVU.MOREX.r3.6HG0545890.1 | HORVU.MOREX.r3.6HG0545890 | chr6H | 16600868 | 16605783 |
| [6,] | cl_1105 | HORVU.MOREX.r3.6HG0545900.1 | HORVU.MOREX.r3.6HG0545900 | chr6H | 16613153 | 16618038 |
| [7,] | cl_1105 | HORVU.MOREX.r3.6HG0545910.1 | HORVU.MOREX.r3.6HG0545910 | chr6H | 16620798 | 16625713 |
| [8,] | cl_1105 | HORVU.MOREX.r3.6HG0545920.1 | HORVU.MOREX.r3.6HG0545920 | chr6H | 16633085 | 16637970 |
| [9,] | cl_1105 | HORVU.MOREX.r3.6HG0545930.1 | HORVU.MOREX.r3.6HG0545930 | chr6H | 16640739 | 16645654 |
| [10,] | cl_1105 | HORVU.MOREX.r3.6HG0545940.1 | HORVU.MOREX.r3.6HG0545940 | chr6H | 16653053 | 16657938 |

cl_1797

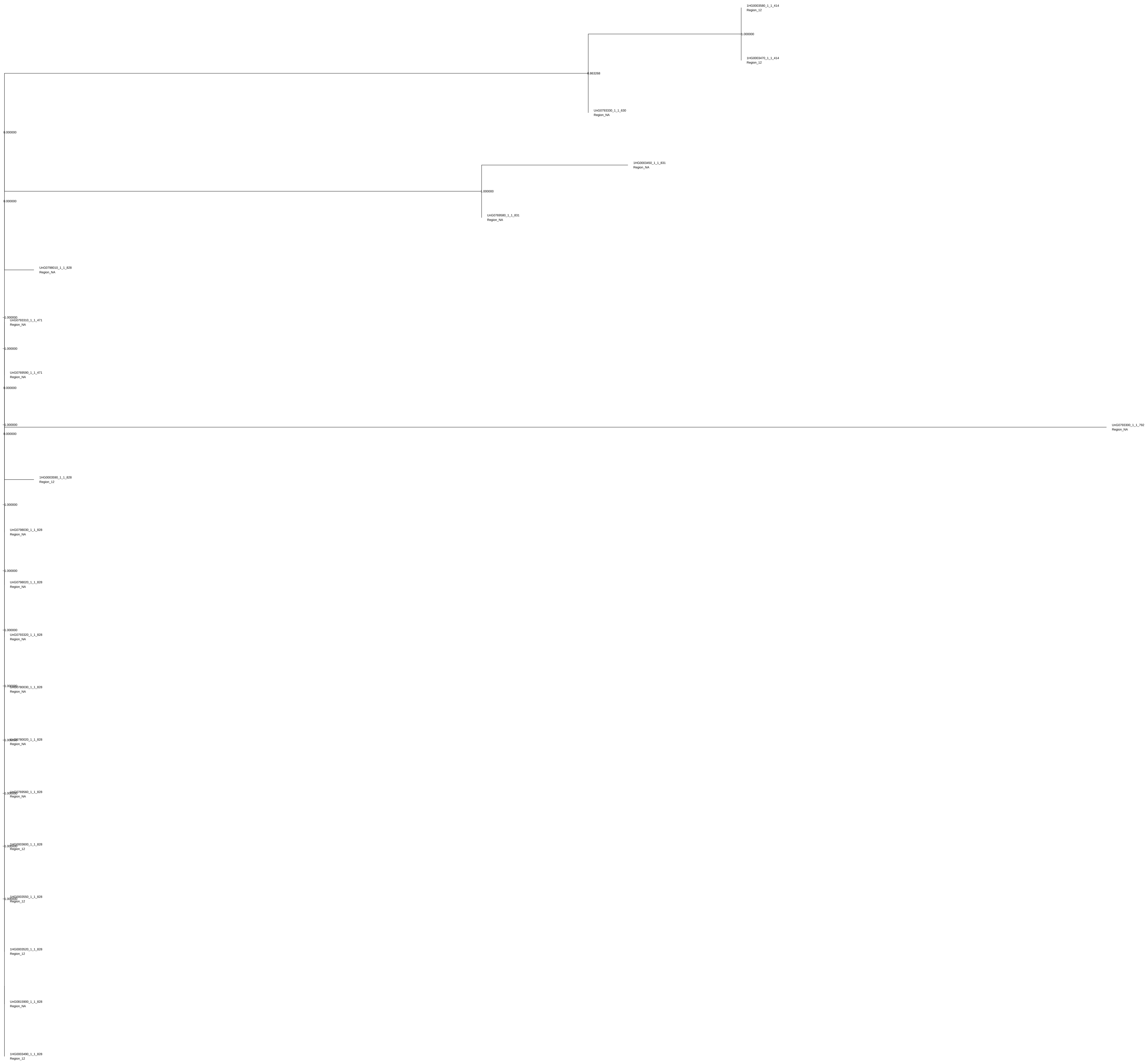

1HG0003580_1_1_414
Region_12
1.000000
1HG0003470_1_1_414
Region_12
0.863268
UnG0793330_1_1_630
Region_NA
0.000000
1HG0003450_1_1_831
Region_NA
1.000000
0.000000
UnG0769580_1_1_831
Region_NA
UnG0798010_1_1_828
Region_NA
1.000000
UnG0793310_1_1_471
Region_NA
1.000000
UnG0769590_1_1_471
Region_NA
0.000000
1.000000
UnG0793300_1_1_792
Region_NA
0.000000
1HG0003590_1_1_828
Region_12
1.000000
UnG0798030_1_1_828
Region_NA
1.000000
UnG0798020_1_1_828
Region_NA
1.000000
UnG0793320_1_1_828
Region_NA
1.000000
UnG0780030_1_1_828
Region_NA
1.000000
UnG0780020_1_1_828
Region_NA
1.000000
UnG0769560_1_1_828
Region_NA
1.000000
1HG0003600_1_1_828
Region_12
1.000000
1HG0003550_1_1_828
Region_12
1HG0003520_1_1_828
Region_12
UnG0815900_1_1_828
Region_NA
1HG0003490_1_1_828
Region_12

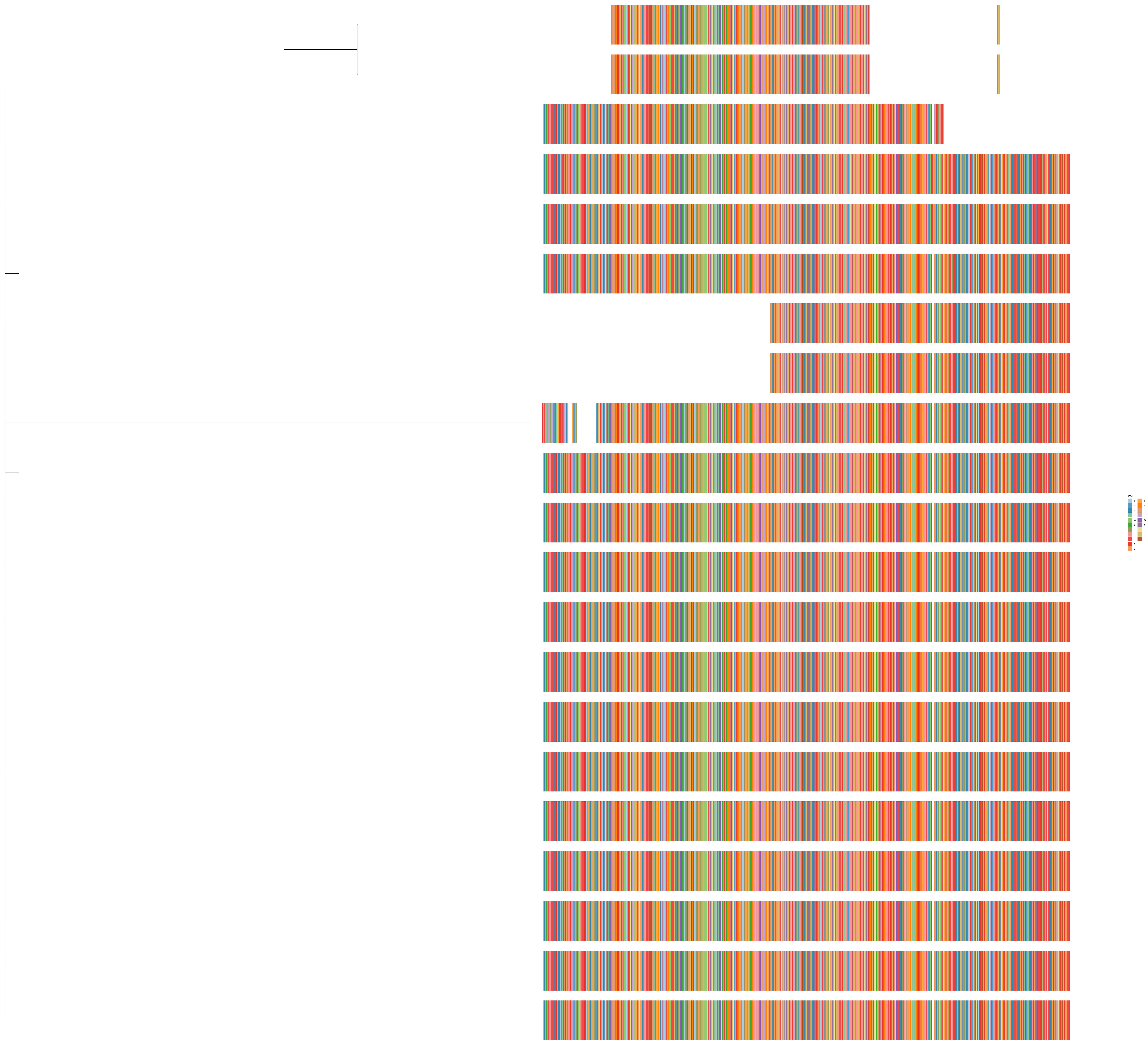
seq
d
r
v
s
w
q
p
l
a
g
t
e
y
i
h
m
k
f
n
c
-

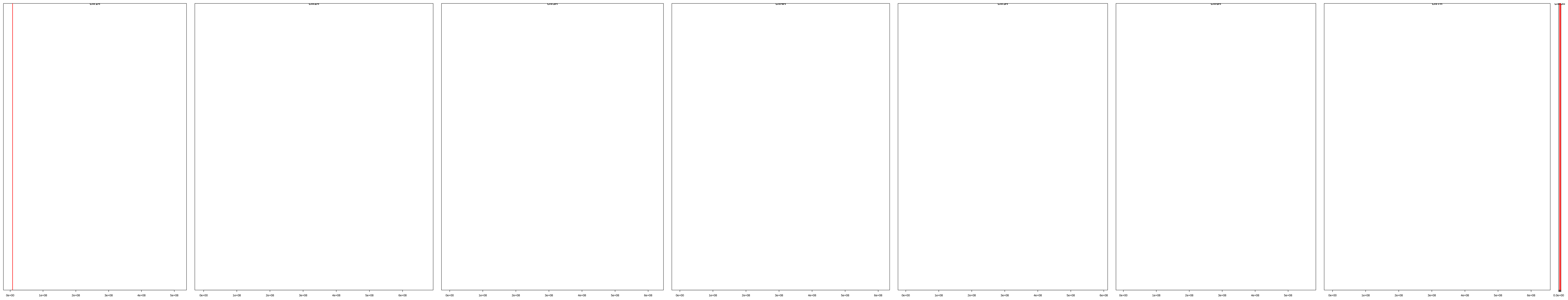

| | description | id | family | seqname | start | end |
|---|---|---|---|---|---|---|
| [1,] | cl_1797 | HORVU.MOREX.r3.1HG0003450.1 | HORVU.MOREX.r3.1HG0003450 | chr1H | 7096113 | 7102052 |
| [2,] | cl_1797 | HORVU.MOREX.r3.1HG0003470.1 | HORVU.MOREX.r3.1HG0003470 | chr1H | 7181710 | 7185886 |
| [3,] | cl_1797 | HORVU.MOREX.r3.1HG0003490.1 | HORVU.MOREX.r3.1HG0003490 | chr1H | 7251051 | 7256659 |
| [4,] | cl_1797 | HORVU.MOREX.r3.1HG0003520.1 | HORVU.MOREX.r3.1HG0003520 | chr1H | 7284171 | 7289779 |
| [5,] | cl_1797 | HORVU.MOREX.r3.1HG0003550.1 | HORVU.MOREX.r3.1HG0003550 | chr1H | 7300728 | 7306336 |
| [6,] | cl_1797 | HORVU.MOREX.r3.1HG0003580.1 | HORVU.MOREX.r3.1HG0003580 | chr1H | 7367378 | 7371554 |
| [7,] | cl_1797 | HORVU.MOREX.r3.1HG0003590.1 | HORVU.MOREX.r3.1HG0003590 | chr1H | 7459640 | 7465248 |
| [8,] | cl_1797 | HORVU.MOREX.r3.1HG0003600.1 | HORVU.MOREX.r3.1HG0003600 | chr1H | 7476205 | 7481813 |
| [9,] | cl_1797 | HORVU.MOREX.r3.UnG0769560.1 | HORVU.MOREX.r3.UnG0769560 | chrUn | 10790034 | 10795644 |
| [10,] | cl_1797 | HORVU.MOREX.r3.UnG0769580.1 | HORVU.MOREX.r3.UnG0769580 | chrUn | 10904805 | 10910464 |
| [11,] | cl_1797 | HORVU.MOREX.r3.UnG0769590.1 | HORVU.MOREX.r3.UnG0769590 | chrUn | 10921417 | 10925116 |
| [12,] | cl_1797 | HORVU.MOREX.r3.UnG0780020.1 | HORVU.MOREX.r3.UnG0780020 | chrUn | 14990774 | 14996382 |
| [13,] | cl_1797 | HORVU.MOREX.r3.UnG0780030.1 | HORVU.MOREX.r3.UnG0780030 | chrUn | 15057016 | 15062624 |
| [14,] | cl_1797 | HORVU.MOREX.r3.UnG0793300.1 | HORVU.MOREX.r3.UnG0793300 | chrUn | 20372338 | 20377836 |
| [15,] | cl_1797 | HORVU.MOREX.r3.UnG0793310.1 | HORVU.MOREX.r3.UnG0793310 | chrUn | 20390696 | 20394395 |
| [16,] | cl_1797 | HORVU.MOREX.r3.UnG0793320.1 | HORVU.MOREX.r3.UnG0793320 | chrUn | 20405346 | 20410954 |
| [17,] | cl_1797 | HORVU.MOREX.r3.UnG0793330.1 | HORVU.MOREX.r3.UnG0793330 | chrUn | 20421900 | 20426914 |
| [18,] | cl_1797 | HORVU.MOREX.r3.UnG0798010.1 | HORVU.MOREX.r3.UnG0798010 | chrUn | 22185371 | 22190979 |
| [19,] | cl_1797 | HORVU.MOREX.r3.UnG0798020.1 | HORVU.MOREX.r3.UnG0798020 | chrUn | 22201927 | 22207535 |
| [20,] | cl_1797 | HORVU.MOREX.r3.UnG0798030.1 | HORVU.MOREX.r3.UnG0798030 | chrUn | 22235047 | 22240655 |
| [21,] | cl_1797 | HORVU.MOREX.r3.UnG0815900.1 | HORVU.MOREX.r3.UnG0815900 | chrUn | 28762251 | 28767859 |

cl_1895

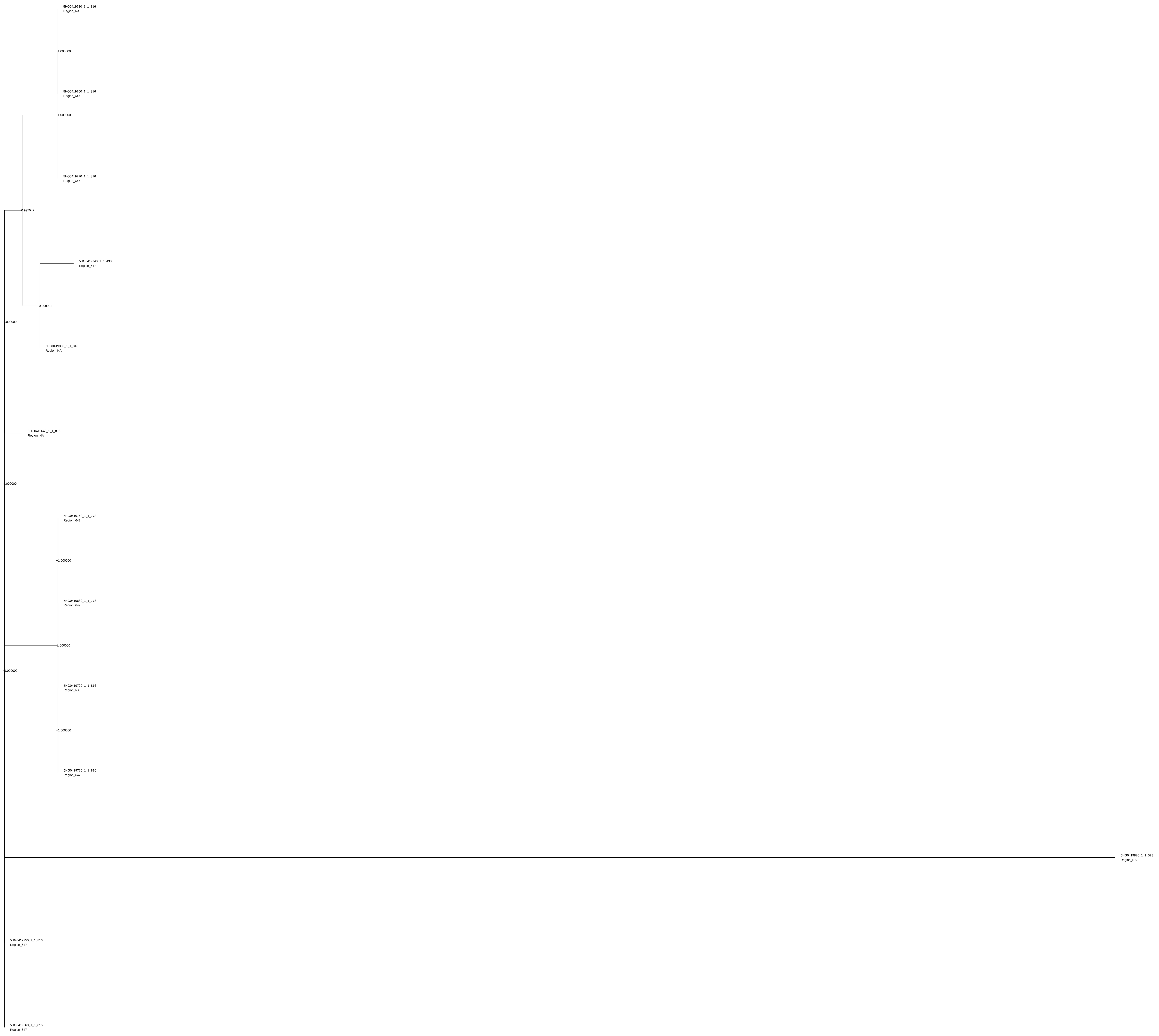

5HG0419780_1_1_816
Region_NA
1.000000
5HG0419700_1_1_816
Region_647
1.000000
5HG0419770_1_1_816
Region_647
0.997542
5HG0419740_1_1_438
Region_647
0.998901
0.000000
5HG0419800_1_1_816
Region_NA
5HG0419640_1_1_816
Region_NA
0.000000
5HG0419760_1_1_778
Region_647
1.000000
5HG0419680_1_1_778
Region_647
1.000000
1.000000
5HG0419790_1_1_816
Region_NA
1.000000
5HG0419720_1_1_816
Region_647
5HG0419820_1_1_573
Region_NA
5HG0419750_1_1_816
Region_647
5HG0419660_1_1_816
Region_647

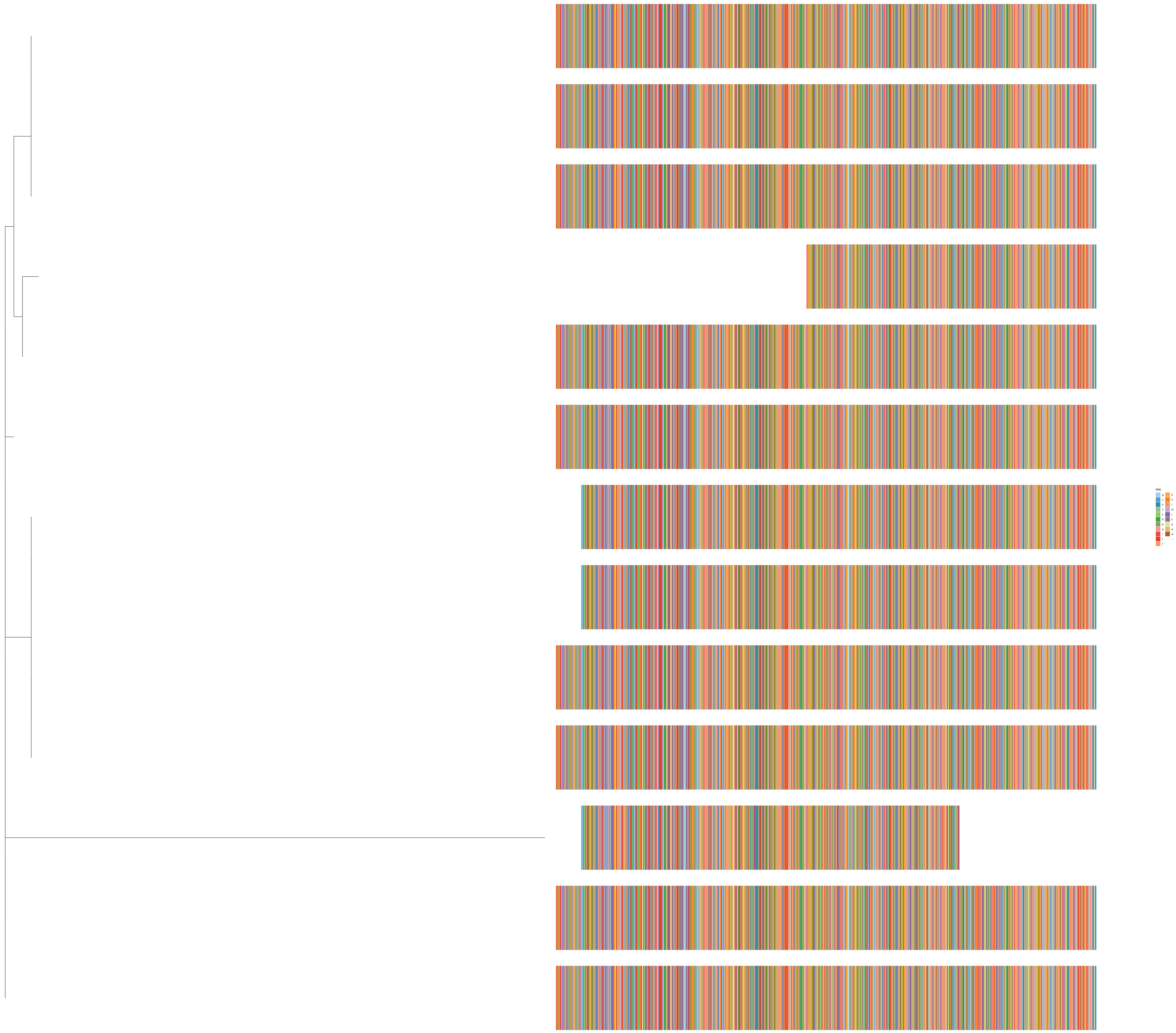
seq
g
v
n
c
s
d
q
a
i
t
f
k
h
l
m
r
y
p
e
w
−

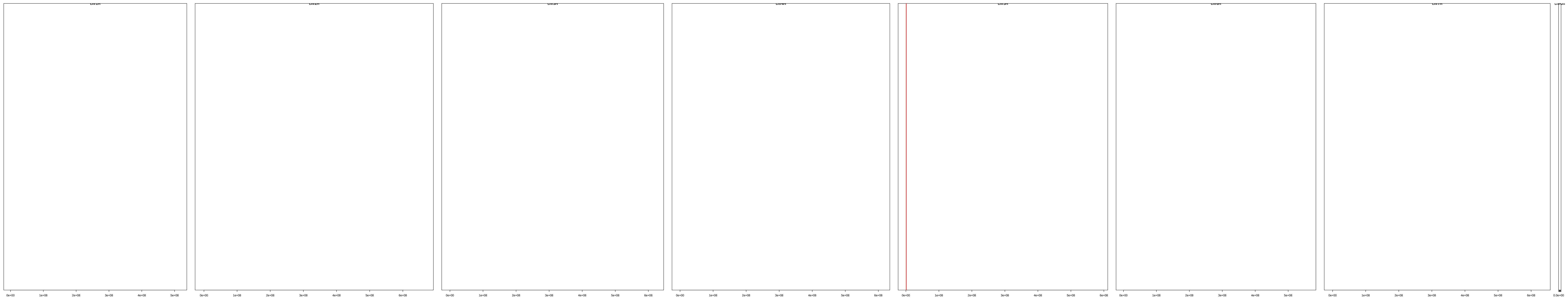

| | description | id | family | seqname | start | end |
|---|---|---|---|---|---|---|
| [1,] | cl_1895 | HORVU.MOREX.r3.5HG0419640.1 | HORVU.MOREX.r3.5HG0419640 | chr5H | 786014 | 790461 |
| [2,] | cl_1895 | HORVU.MOREX.r3.5HG0419660.1 | HORVU.MOREX.r3.5HG0419660 | chr5H | 819442 | 823889 |
| [3,] | cl_1895 | HORVU.MOREX.r3.5HG0419680.1 | HORVU.MOREX.r3.5HG0419680 | chr5H | 888419 | 893081 |
| [4,] | cl_1895 | HORVU.MOREX.r3.5HG0419700.1 | HORVU.MOREX.r3.5HG0419700 | chr5H | 925009 | 929456 |
| [5,] | cl_1895 | HORVU.MOREX.r3.5HG0419720.1 | HORVU.MOREX.r3.5HG0419720 | chr5H | 958455 | 962902 |
| [6,] | cl_1895 | HORVU.MOREX.r3.5HG0419740.1 | HORVU.MOREX.r3.5HG0419740 | chr5H | 995982 | 999295 |
| [7,] | cl_1895 | HORVU.MOREX.r3.5HG0419750.1 | HORVU.MOREX.r3.5HG0419750 | chr5H | 1030770 | 1035217 |
| [8,] | cl_1895 | HORVU.MOREX.r3.5HG0419760.1 | HORVU.MOREX.r3.5HG0419760 | chr5H | 1101274 | 1139392 |
| [9,] | cl_1895 | HORVU.MOREX.r3.5HG0419770.1 | HORVU.MOREX.r3.5HG0419770 | chr5H | 1101474 | 1105921 |
| [10,] | cl_1895 | HORVU.MOREX.r3.5HG0419780.1 | HORVU.MOREX.r3.5HG0419780 | chr5H | 1171271 | 1175718 |
| [11,] | cl_1895 | HORVU.MOREX.r3.5HG0419790.1 | HORVU.MOREX.r3.5HG0419790 | chr5H | 1204714 | 1209161 |
| [12,] | cl_1895 | HORVU.MOREX.r3.5HG0419800.1 | HORVU.MOREX.r3.5HG0419800 | chr5H | 1240861 | 1245722 |
| [13,] | cl_1895 | HORVU.MOREX.r3.5HG0419820.1 | HORVU.MOREX.r3.5HG0419820 | chr5H | 1277146 | 1280864 |

cl_805

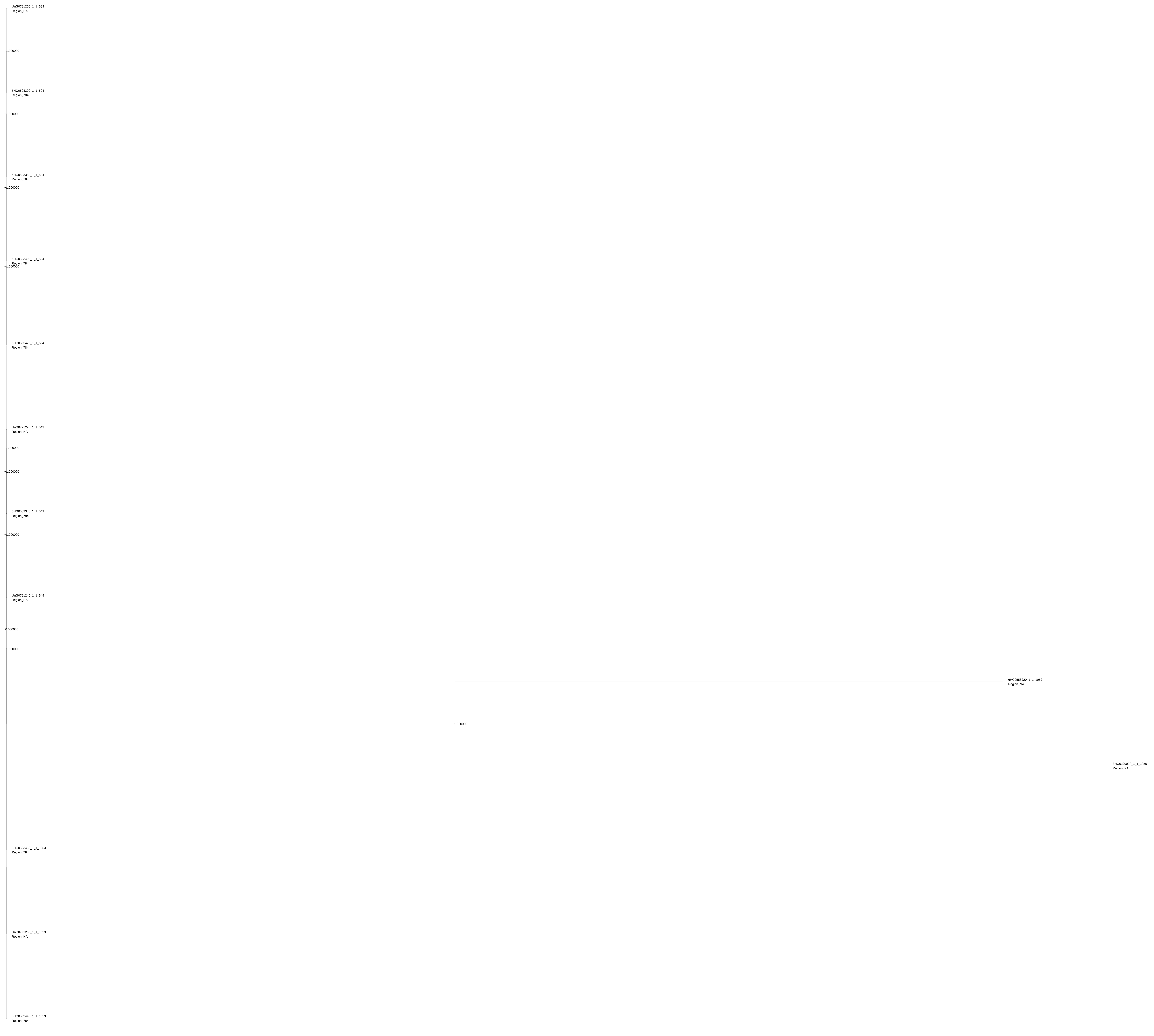

UnG0791200_1_1_594
Region_NA
1.000000
5HG0503300_1_1_594
Region_784
1.000000
5HG0503380_1_1_594
Region_784
1.000000
5HG0503400_1_1_594
Region_784
1.000000
5HG0503420_1_1_594
Region_784
UnG0791290_1_1_549
Region_NA
1.000000
1.000000
5HG0503340_1_1_549
Region_784
1.000000
UnG0791240_1_1_549
Region_NA
0.000000
1.000000
6HG0558220_1_1_1052
Region_NA
1.000000
3HG0229090_1_1_1056
Region_NA
5HG0503450_1_1_1053
Region_784
UnG0791250_1_1_1053
Region_NA
5HG0503440_1_1_1053
Region_784

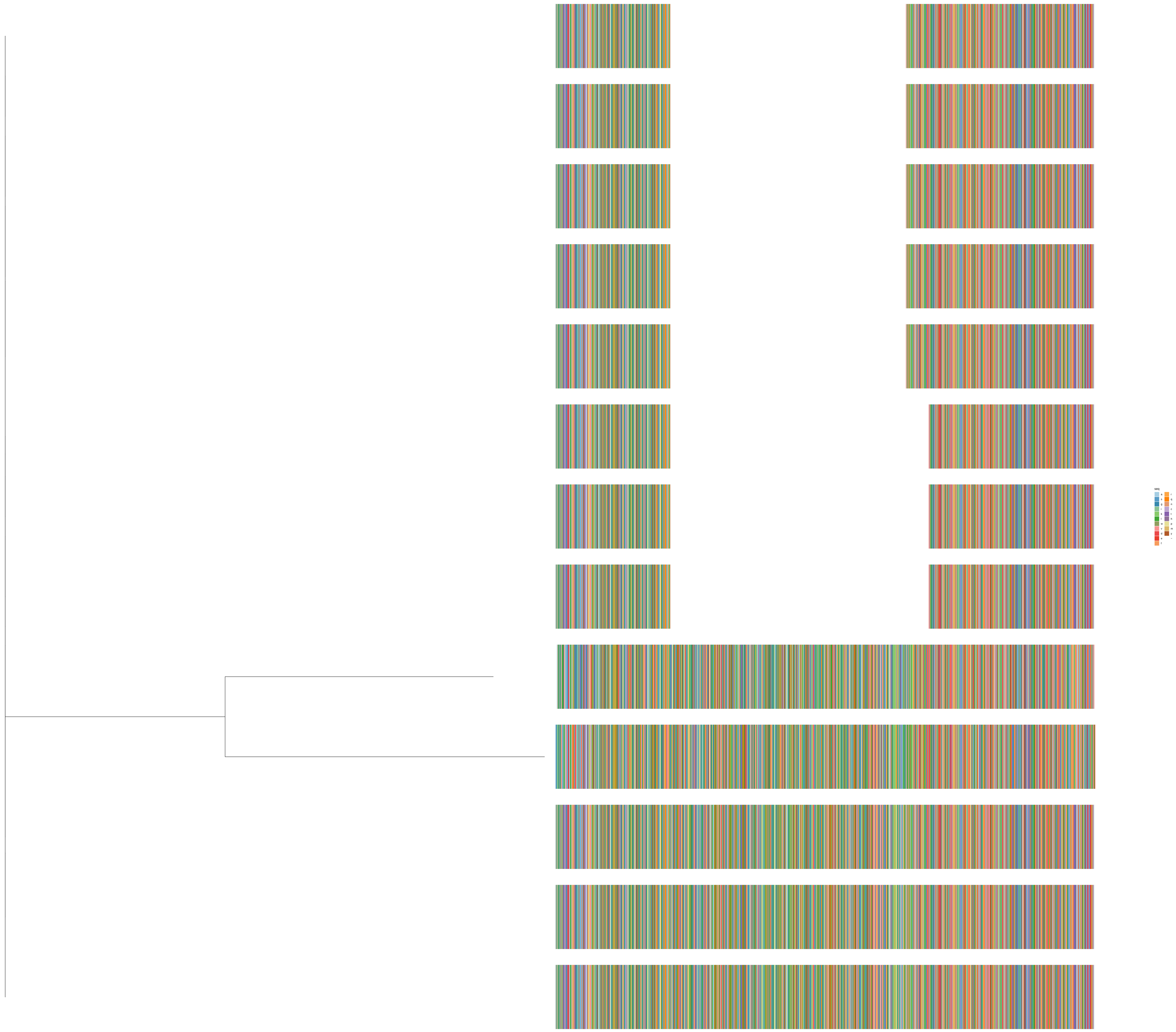

seq
a
s
g
i
k
l
w
v
d
e
f
r
q
n
c
t
h
p
m
y
−

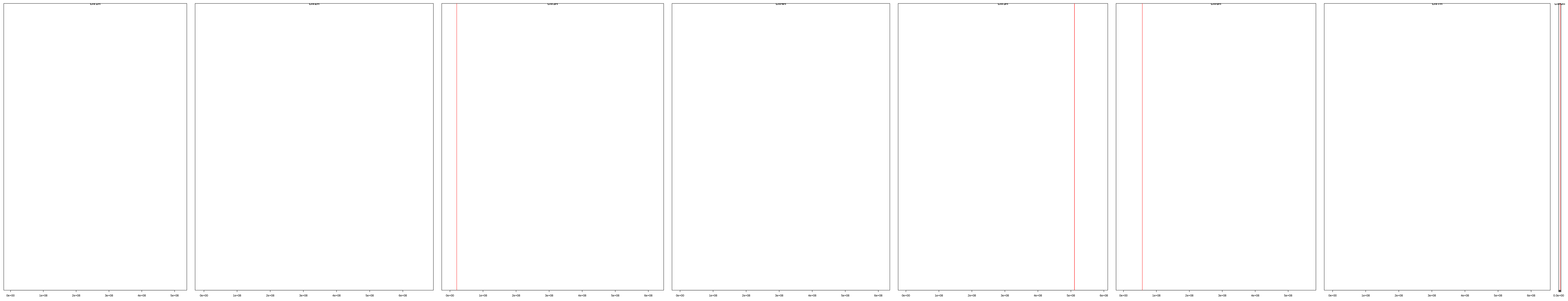

| | description | id | family | seqname | start | end |
|---|---|---|---|---|---|---|
| [1,] | cl_805 | HORVU.MOREX.r3.3HG0229090.1 | HORVU.MOREX.r3.3HG0229090 | chr3H | 20530522 | 20536004 |
| [2,] | cl_805 | HORVU.MOREX.r3.5HG0503300.1 | HORVU.MOREX.r3.5HG0503300 | chr5H | 510896929 | 510902293 |
| [3,] | cl_805 | HORVU.MOREX.r3.5HG0503340.1 | HORVU.MOREX.r3.5HG0503340 | chr5H | 510912001 | 510917237 |
| [4,] | cl_805 | HORVU.MOREX.r3.5HG0503380.1 | HORVU.MOREX.r3.5HG0503380 | chr5H | 510927057 | 510932293 |
| [5,] | cl_805 | HORVU.MOREX.r3.5HG0503400.1 | HORVU.MOREX.r3.5HG0503400 | chr5H | 510942108 | 510947344 |
| [6,] | cl_805 | HORVU.MOREX.r3.5HG0503420.1 | HORVU.MOREX.r3.5HG0503420 | chr5H | 510957164 | 510962400 |
| [7,] | cl_805 | HORVU.MOREX.r3.5HG0503440.1 | HORVU.MOREX.r3.5HG0503440 | chr5H | 510972215 | 510977451 |
| [8,] | cl_805 | HORVU.MOREX.r3.5HG0503450.1 | HORVU.MOREX.r3.5HG0503450 | chr5H | 510987267 | 510992503 |
| [9,] | cl_805 | HORVU.MOREX.r3.6HG0558220.1 | HORVU.MOREX.r3.6HG0558220 | chr6H | 57274822 | 57280343 |
| [10,] | cl_805 | HORVU.MOREX.r3.UnG0791200.1 | HORVU.MOREX.r3.UnG0791200 | chrUn | 19191355 | 19196591 |
| [11,] | cl_805 | HORVU.MOREX.r3.UnG0791240.1 | HORVU.MOREX.r3.UnG0791240 | chrUn | 19206407 | 19211643 |
| [12,] | cl_805 | HORVU.MOREX.r3.UnG0791250.1 | HORVU.MOREX.r3.UnG0791250 | chrUn | 19221457 | 19226693 |
| [13,] | cl_805 | HORVU.MOREX.r3.UnG0791290.1 | HORVU.MOREX.r3.UnG0791290 | chrUn | 19236508 | 19241744 |

cl_738

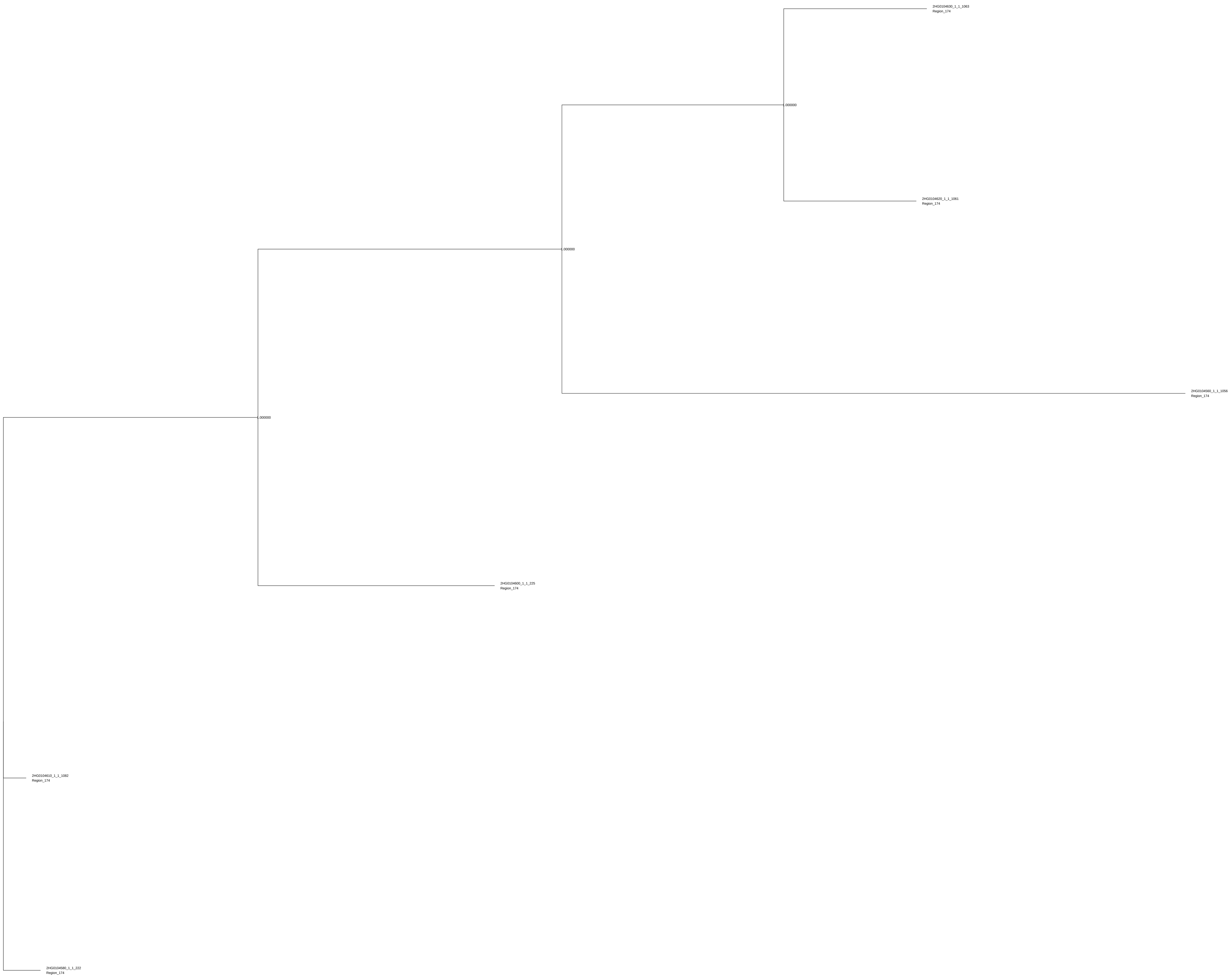
2HG0104630_1_1_1063
Region_174
1.000000
2HG0104620_1_1_1061
Region_174
1.000000
2HG0104560_1_1_1056
Region_174
1.000000
2HG0104600_1_1_225
Region_174
2HG0104610_1_1_1082
Region_174
2HG0104580_1_1_222
Region_174

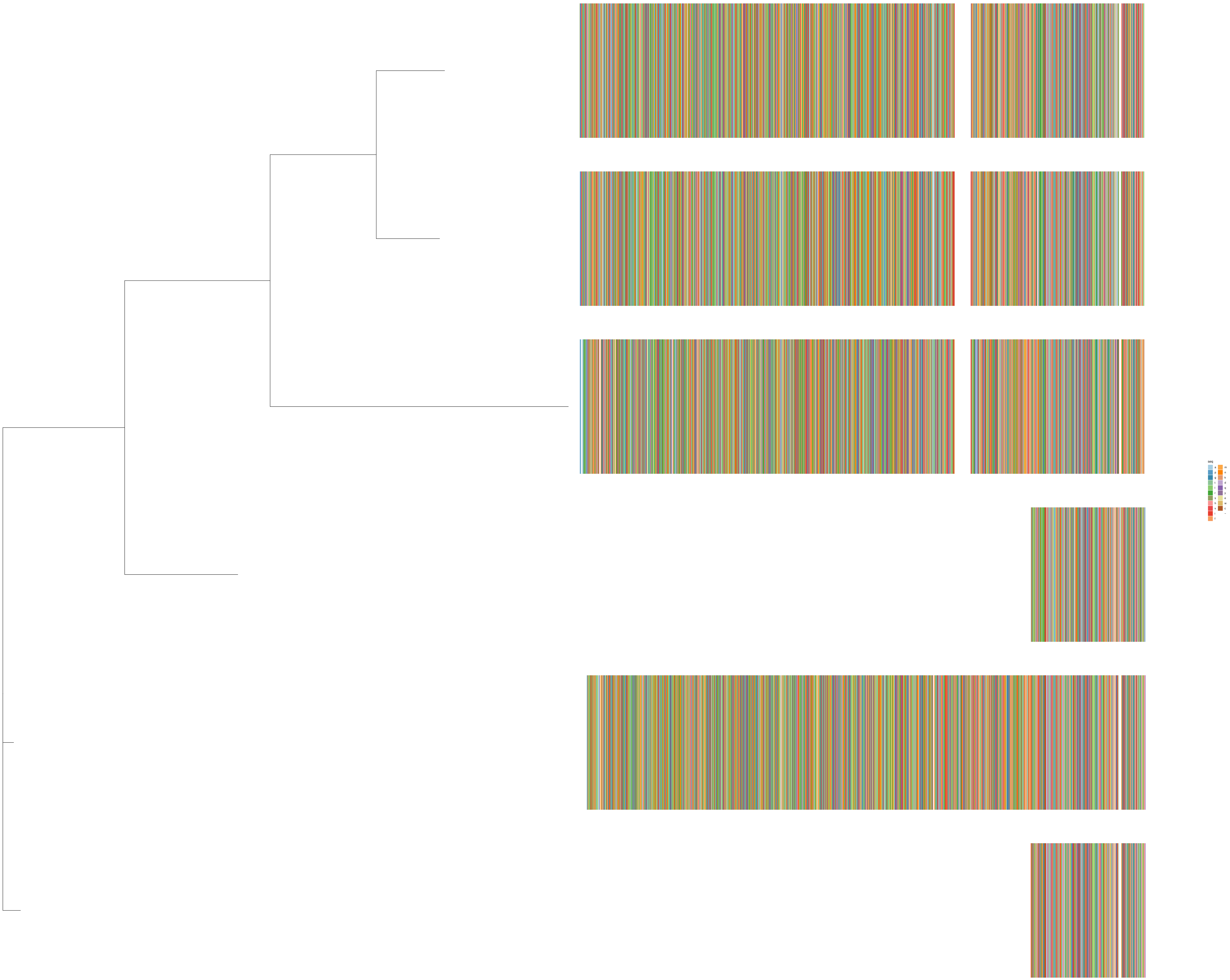
seq
a
p
g
t
l
r
s
h
v
i
f
m
n
k
d
q
y
e
w
c
−

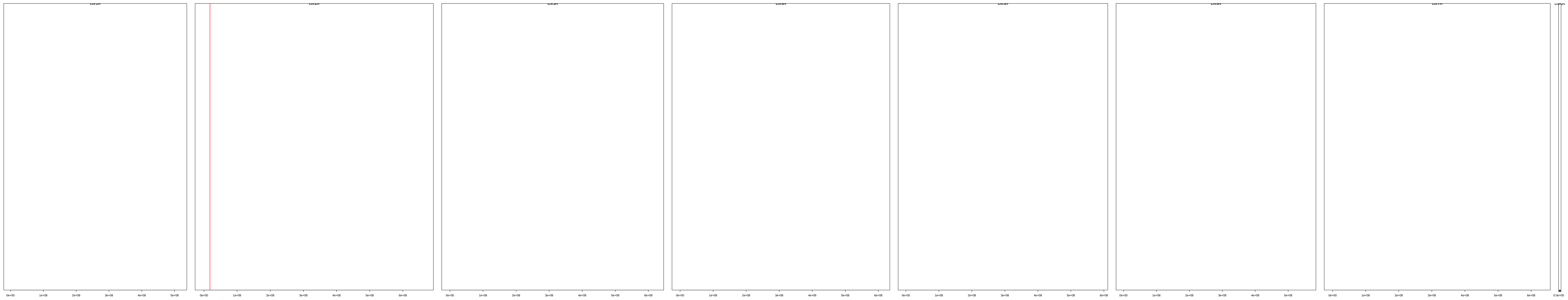

| | description | id | family | seqname | start | end |
|---|---|---|---|---|---|---|
| [1,] | cl_738 | HORVU.MOREX.r3.2HG0104560.1 | HORVU.MOREX.r3.2HG0104560 | chr2H | 18123254 | 18128495 |
| [2,] | cl_738 | HORVU.MOREX.r3.2HG0104580.1 | HORVU.MOREX.r3.2HG0104580 | chr2H | 18144579 | 18147376 |
| [3,] | cl_738 | HORVU.MOREX.r3.2HG0104600.1 | HORVU.MOREX.r3.2HG0104600 | chr2H | 18155989 | 18158735 |
| [4,] | cl_738 | HORVU.MOREX.r3.2HG0104610.1 | HORVU.MOREX.r3.2HG0104610 | chr2H | 18210501 | 18215878 |
| [5,] | cl_738 | HORVU.MOREX.r3.2HG0104620.1 | HORVU.MOREX.r3.2HG0104620 | chr2H | 18215895 | 18221450 |
| [6,] | cl_738 | HORVU.MOREX.r3.2HG0104630.1 | HORVU.MOREX.r3.2HG0104630 | chr2H | 18220287 | 18226955 |

cl_2987

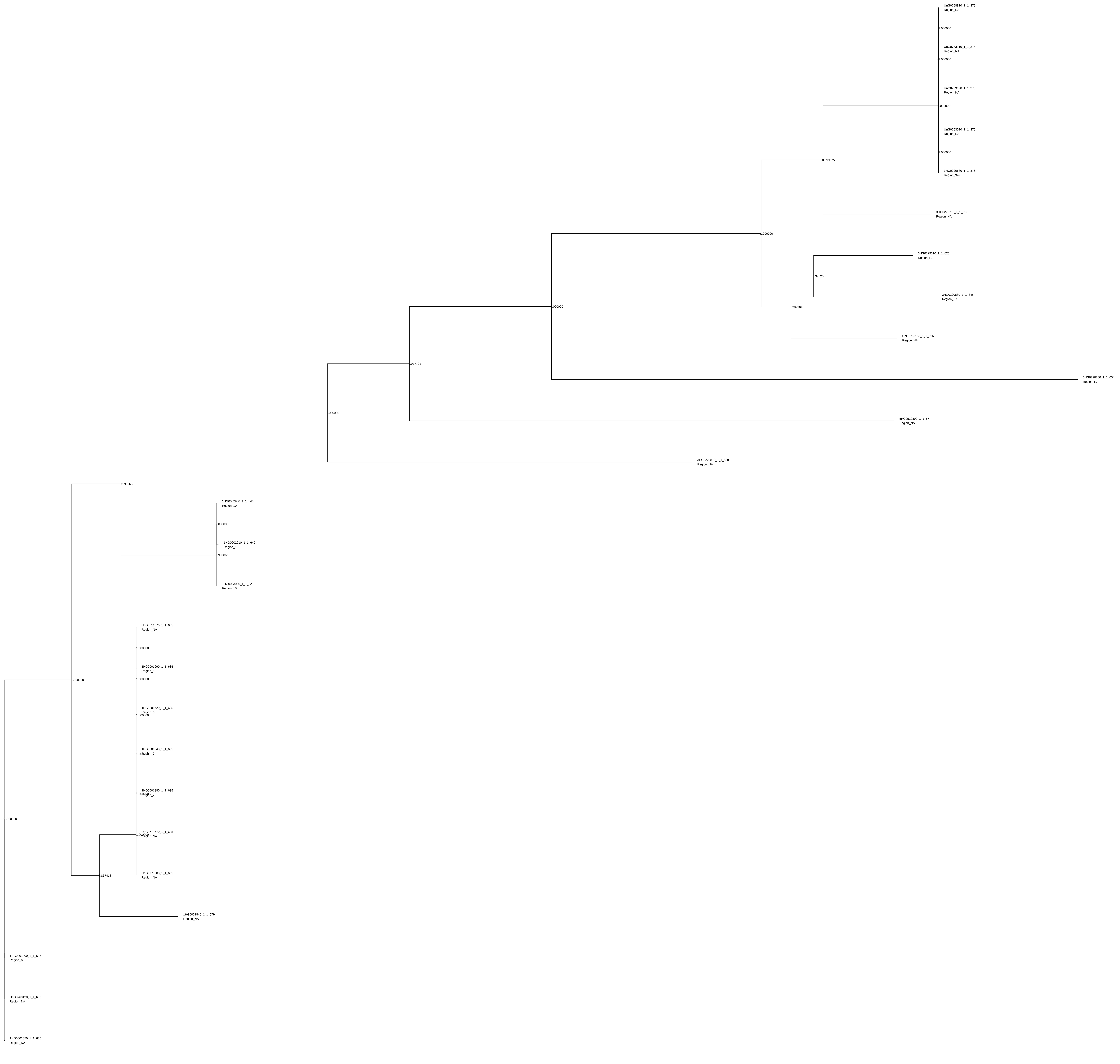
UnG0758810_1_1_375
Region_NA
1.000000
UnG0753110_1_1_375
Region_NA
1.000000
UnG0753120_1_1_375
Region_NA
1.000000
UnG0753020_1_1_376
Region_NA
1.000000
3HG0220680_1_1_376
Region_349
0.999975
3HG0220750_1_1_617
Region_NA
1.000000
3HG0229310_1_1_626
Region_NA
0.973263
3HG0220880_1_1_345
Region_NA
0.989964
UnG0753150_1_1_626
Region_NA
1.000000
3HG0220260_1_1_654
Region_NA
0.977721
5HG0510390_1_1_677
Region_NA
1.000000
3HG0220810_1_1_638
Region_NA
0.998668
1HG0002980_1_1_646
Region_10
0.000000
1HG0002910_1_1_640
Region_10
0.999865
1HG0003030_1_1_328
Region_10
UnG0811670_1_1_635
Region_NA
1.000000
1HG0001690_1_1_635
Region_6
1.000000
1.000000
1HG0001720_1_1_635
Region_6
1.000000
1HG0001640_1_1_635
Region_7
1.000000
1HG0001680_1_1_635
Region_7
1.000000
UnG0773770_1_1_635
Region_NA
1.000000
UnG0773800_1_1_635
Region_NA
0.867418
1HG0002840_1_1_579
Region_NA
1.000000
1HG0001800_1_1_635
Region_6
UnG0769130_1_1_635
Region_NA
1HG0001650_1_1_635
Region_NA

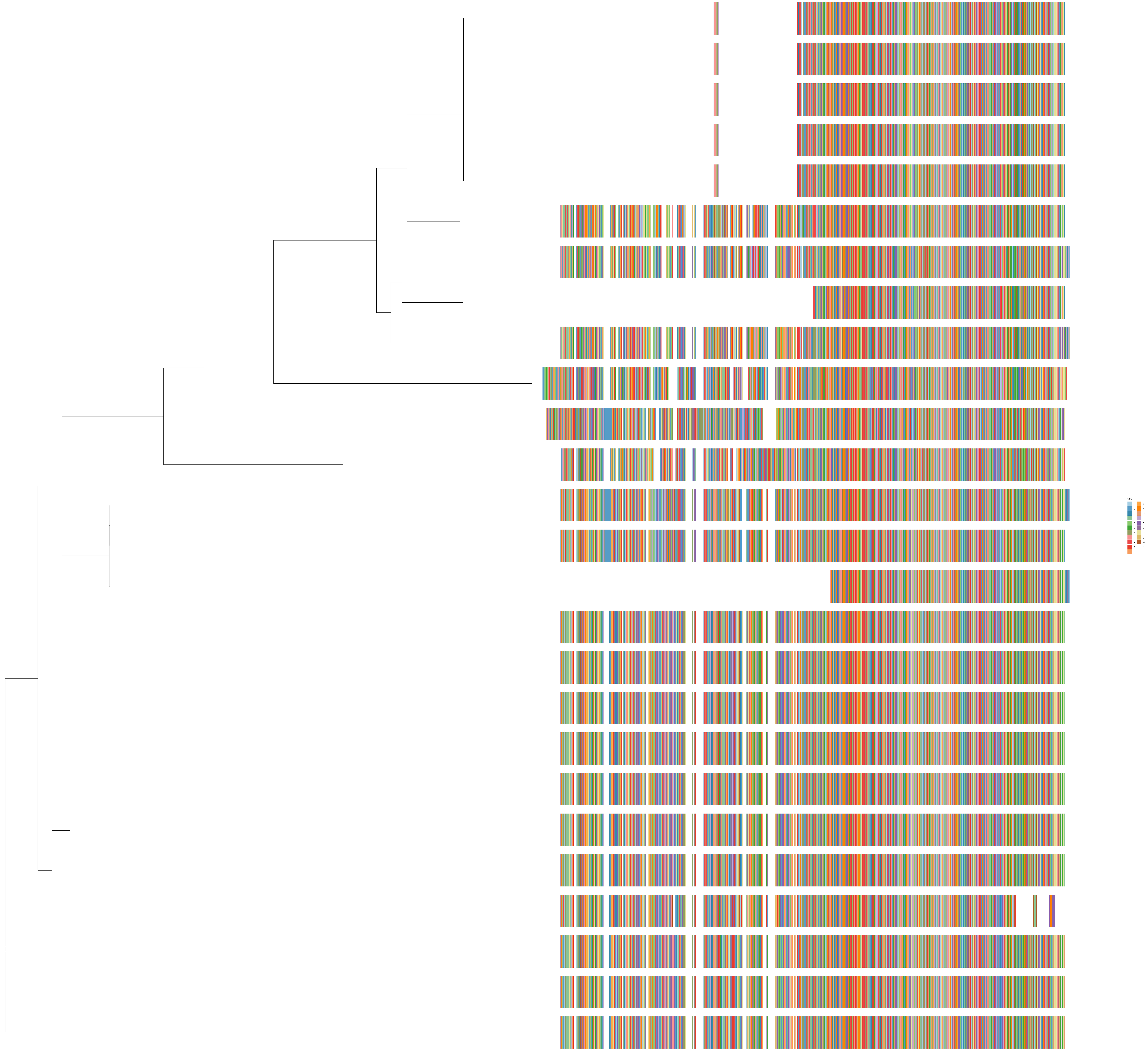
seq
i
s
t
l
q
e
a
f
v
g
h
c
k
m
n
r
d
p
y
w
–

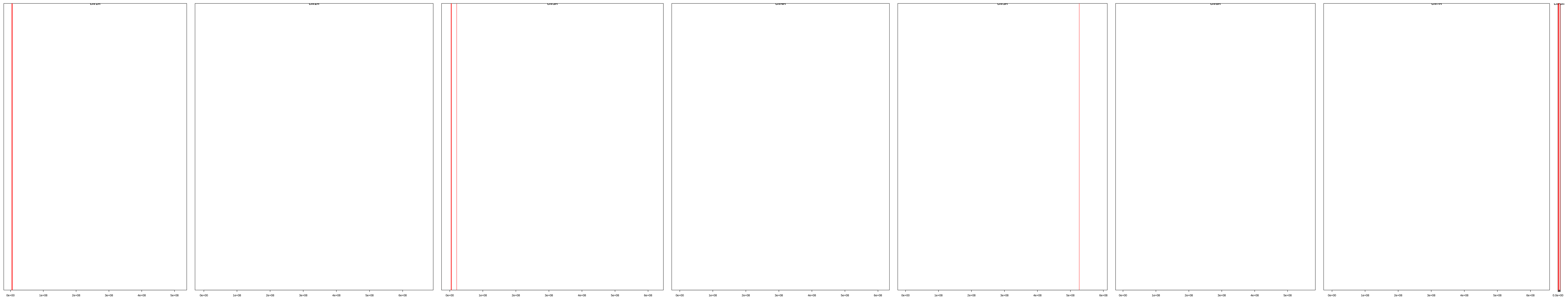

| | description | id | family | seqname | start | end |
|---|---|---|---|---|---|---|
| **[1,]** | cl_2987 | HORVU.MOREX.r3.1HG0001650.1 | HORVU.MOREX.r3.1HG0001650 | chr1H | 3825892 | 3830247 |
| **[2,]** | cl_2987 | HORVU.MOREX.r3.1HG0001690.1 | HORVU.MOREX.r3.1HG0001690 | chr1H | 3913908 | 3919194 |
| **[3,]** | cl_2987 | HORVU.MOREX.r3.1HG0001720.1 | HORVU.MOREX.r3.1HG0001720 | chr1H | 4005789 | 4011075 |
| **[4,]** | cl_2987 | HORVU.MOREX.r3.1HG0001800.1 | HORVU.MOREX.r3.1HG0001800 | chr1H | 4118119 | 4122475 |
| **[5,]** | cl_2987 | HORVU.MOREX.r3.1HG0001840.1 | HORVU.MOREX.r3.1HG0001840 | chr1H | 4206096 | 4211382 |
| **[6,]** | cl_2987 | HORVU.MOREX.r3.1HG0001880.1 | HORVU.MOREX.r3.1HG0001880 | chr1H | 4297978 | 4303264 |
| **[7,]** | cl_2987 | HORVU.MOREX.r3.1HG0002840.1 | HORVU.MOREX.r3.1HG0002840 | chr1H | 5693036 | 5699346 |
| **[8,]** | cl_2987 | HORVU.MOREX.r3.1HG0002910.1 | HORVU.MOREX.r3.1HG0002910 | chr1H | 5748821 | 5753339 |
| **[9,]** | cl_2987 | HORVU.MOREX.r3.1HG0002980.1 | HORVU.MOREX.r3.1HG0002980 | chr1H | 5790397 | 5795210 |
| **[10,]** | cl_2987 | HORVU.MOREX.r3.1HG0003030.1 | HORVU.MOREX.r3.1HG0003030 | chr1H | 5832270 | 5835530 |
| **[11,]** | cl_2987 | HORVU.MOREX.r3.3HG0220260.1 | HORVU.MOREX.r3.3HG0220260 | chr3H | 3942757 | 3948072 |
| **[12,]** | cl_2987 | HORVU.MOREX.r3.3HG0220680.1 | HORVU.MOREX.r3.3HG0220680 | chr3H | 4858345 | 4861629 |
| **[13,]** | cl_2987 | HORVU.MOREX.r3.3HG0220750.1 | HORVU.MOREX.r3.3HG0220750 | chr3H | 4987401 | 4992055 |
| **[14,]** | cl_2987 | HORVU.MOREX.r3.3HG0220810.1 | HORVU.MOREX.r3.3HG0220810 | chr3H | 5073126 | 5078470 |
| **[15,]** | cl_2987 | HORVU.MOREX.r3.3HG0220880.1 | HORVU.MOREX.r3.3HG0220880 | chr3H | 5101856 | 5105082 |
| **[16,]** | cl_2987 | HORVU.MOREX.r3.3HG0229310.1 | HORVU.MOREX.r3.3HG0229310 | chr3H | 21064722 | 21069378 |
| **[17,]** | cl_2987 | HORVU.MOREX.r3.5HG0510390.1 | HORVU.MOREX.r3.5HG0510390 | chr5H | 526704224 | 526709467 |
| **[18,]** | cl_2987 | HORVU.MOREX.r3.UnG0753020.1 | HORVU.MOREX.r3.UnG0753020 | chrUn | 158902 | 162186 |
| **[19,]** | cl_2987 | HORVU.MOREX.r3.UnG0753110.1 | HORVU.MOREX.r3.UnG0753110 | chrUn | 412268 | 415552 |
| **[20,]** | cl_2987 | HORVU.MOREX.r3.UnG0753120.1 | HORVU.MOREX.r3.UnG0753120 | chrUn | 425091 | 428375 |
| **[21,]** | cl_2987 | HORVU.MOREX.r3.UnG0753150.1 | HORVU.MOREX.r3.UnG0753150 | chrUn | 475790 | 481072 |
| **[22,]** | cl_2987 | HORVU.MOREX.r3.UnG0758810.1 | HORVU.MOREX.r3.UnG0758810 | chrUn | 5085422 | 5088706 |
| **[23,]** | cl_2987 | HORVU.MOREX.r3.UnG0769130.1 | HORVU.MOREX.r3.UnG0769130 | chrUn | 10521447 | 10525803 |
| **[24,]** | cl_2987 | HORVU.MOREX.r3.UnG0773770.1 | HORVU.MOREX.r3.UnG0773770 | chrUn | 12677337 | 12682623 |
| **[25,]** | cl_2987 | HORVU.MOREX.r3.UnG0773800.1 | HORVU.MOREX.r3.UnG0773800 | chrUn | 12765981 | 12771267 |
| **[26,]** | cl_2987 | HORVU.MOREX.r3.UnG0811670.1 | HORVU.MOREX.r3.UnG0811670 | chrUn | 27274130 | 27279416 |

cl_3714

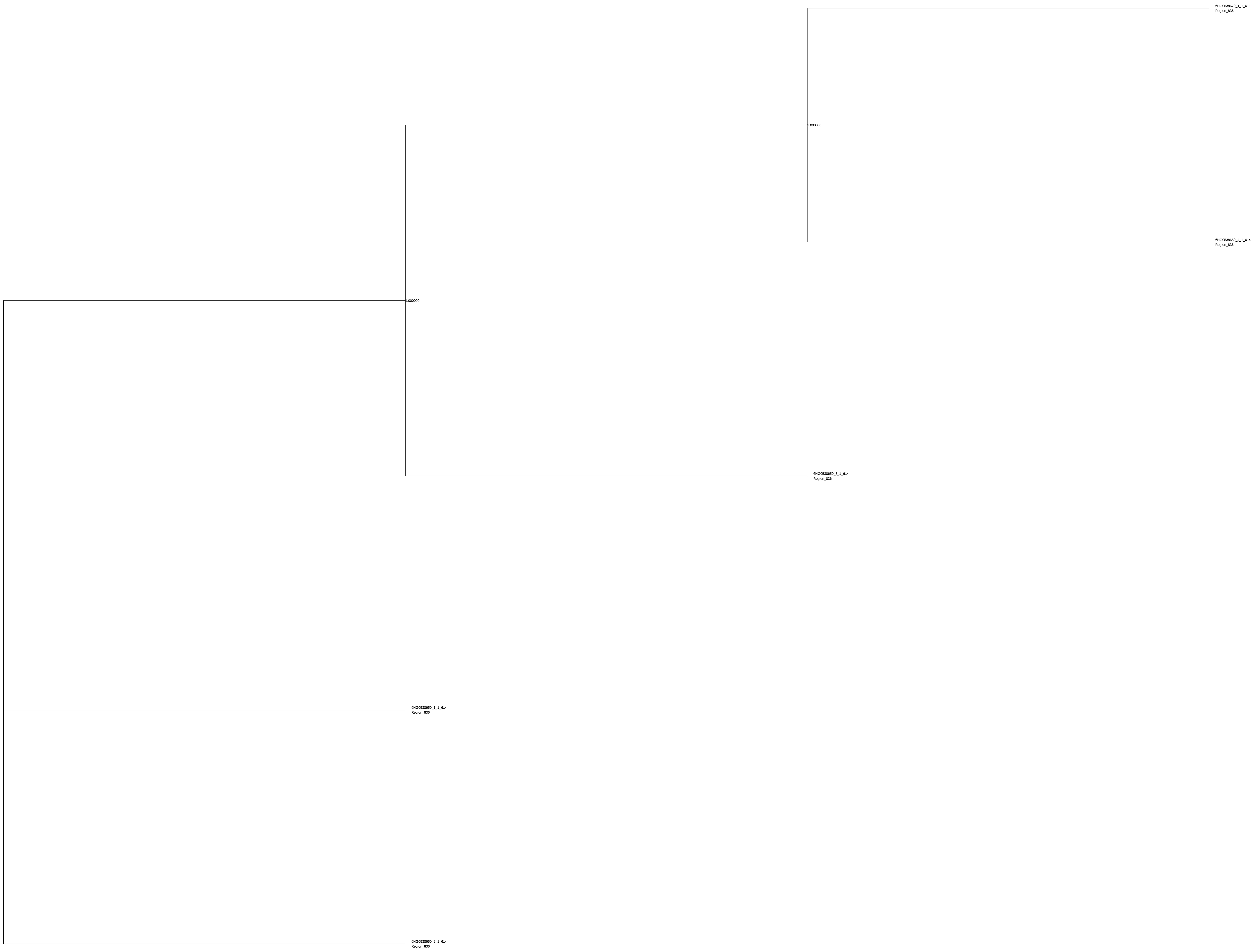
6HG0538670_1_1_611
Region_836
1.000000
6HG0538650_4_1_614
Region_836
1.000000
6HG0538650_3_1_614
Region_836
6HG0538650_1_1_614
Region_836
6HG0538650_2_1_614
Region_836

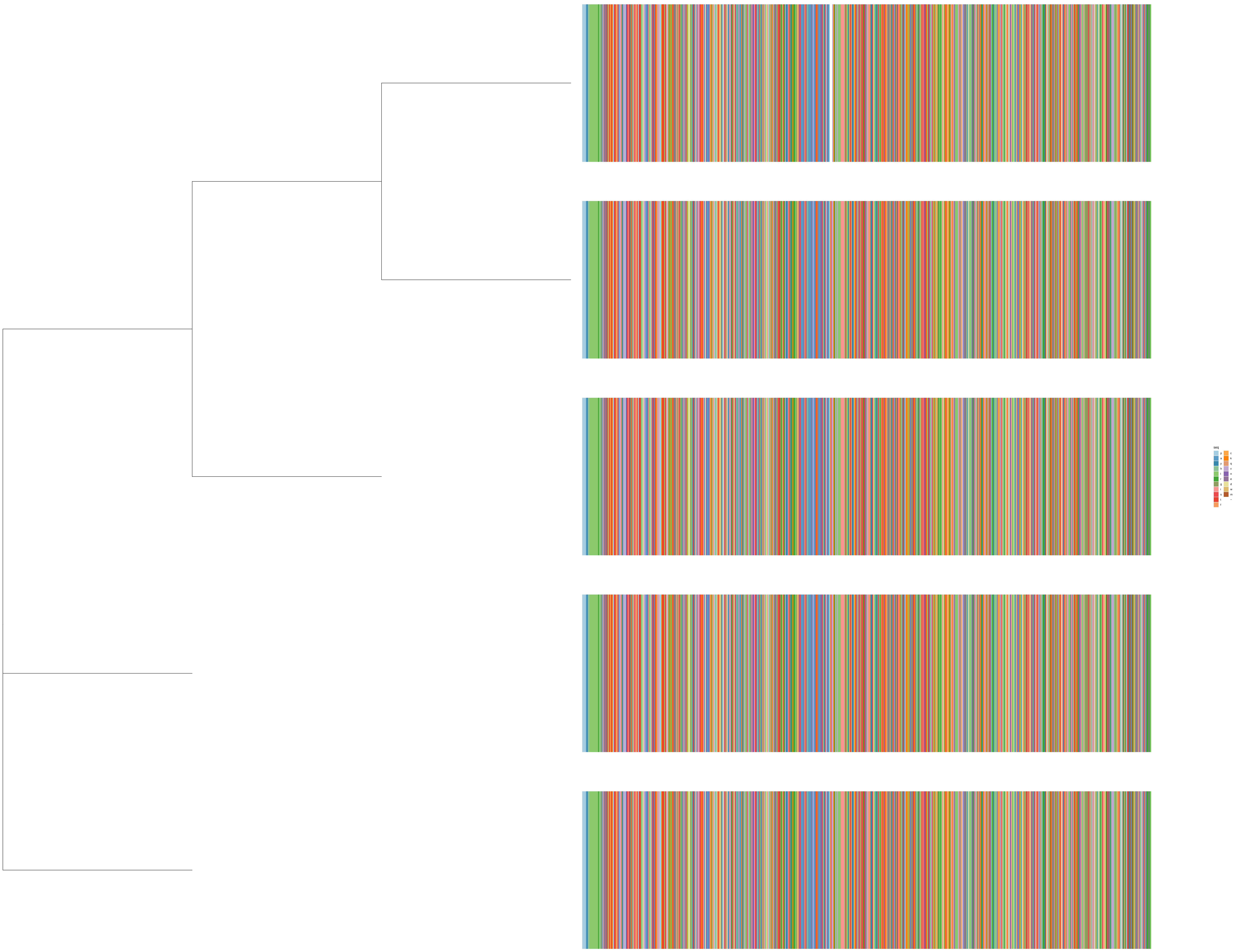
seq
p
a
y
h
l
r
g
i
s
t
f
c
k
q
v
n
e
d
w
m
–

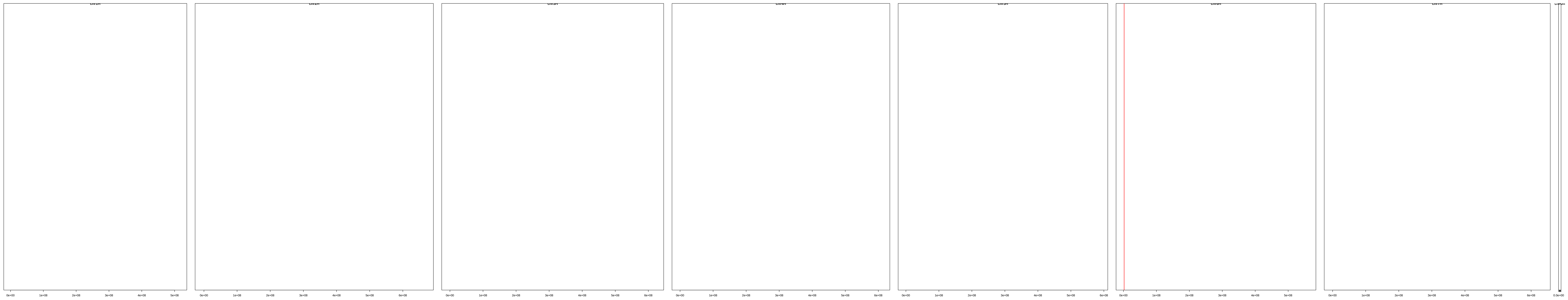

| | description | id | family | seqname | start | end |
|---|---|---|---|---|---|---|
| [1,] | cl_3714 | HORVU.MOREX.r3.6HG0538650.1 | HORVU.MOREX.r3.6HG0538650 | chr6H | 1966949 | 1975176 |
| [2,] | cl_3714 | HORVU.MOREX.r3.6HG0538650.2 | HORVU.MOREX.r3.6HG0538650 | chr6H | 1966949 | 1975182 |
| [3,] | cl_3714 | HORVU.MOREX.r3.6HG0538650.3 | HORVU.MOREX.r3.6HG0538650 | chr6H | 1966949 | 1975184 |
| [4,] | cl_3714 | HORVU.MOREX.r3.6HG0538650.4 | HORVU.MOREX.r3.6HG0538650 | chr6H | 1966949 | 1975186 |
| [5,] | cl_3714 | HORVU.MOREX.r3.6HG0538670.1 | HORVU.MOREX.r3.6HG0538670 | chr6H | 2044171 | 2051826 |

cl_1888

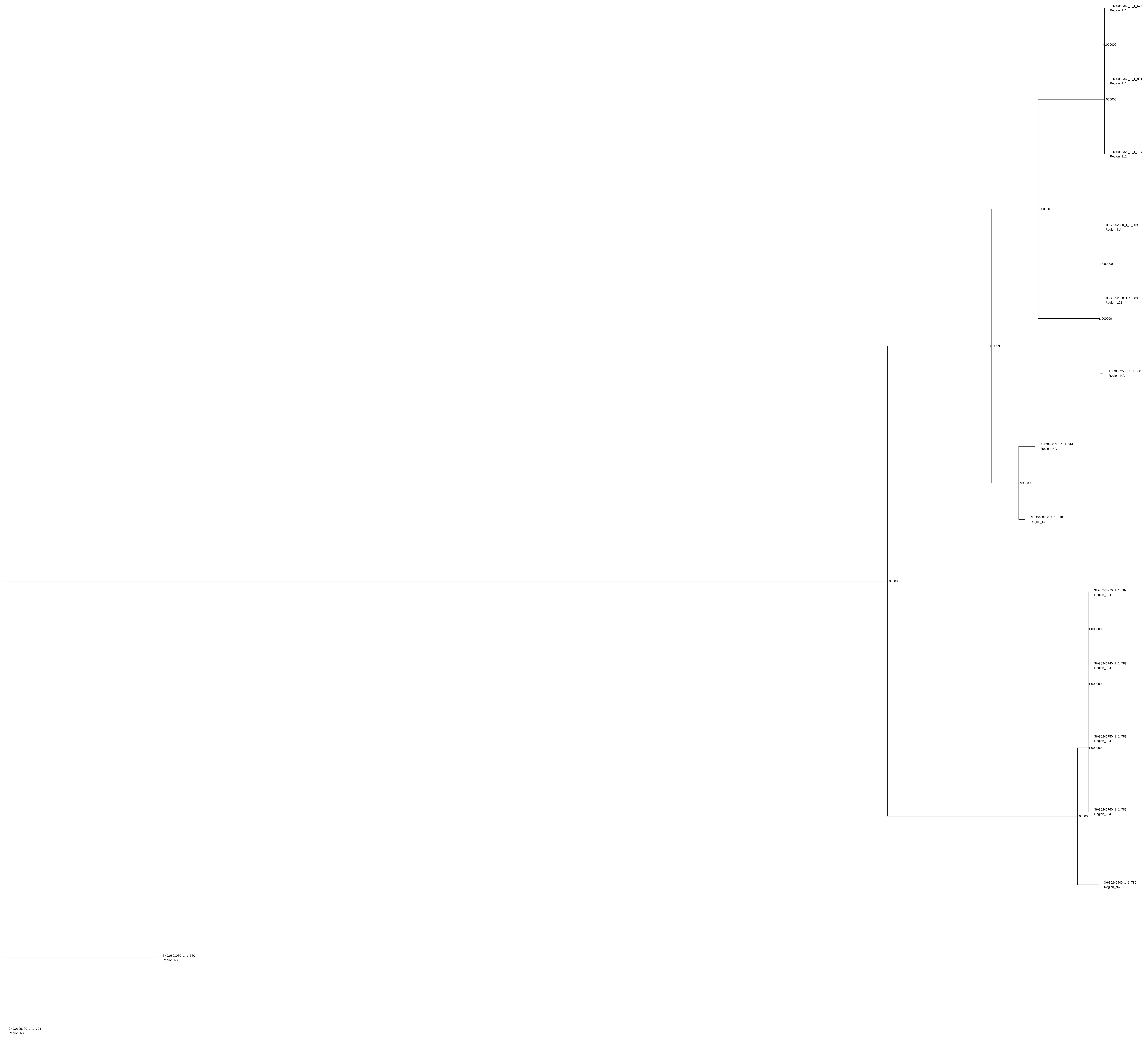
1HG0062340_1_1_675
Region_111
0.000000
1HG0062360_1_1_801
Region_111
1.000000
1HG0062320_1_1_184
Region_111
1.000000
1HG0052580_1_1_809
Region_NA
1.000000
1HG0052560_1_1_809
Region_102
1.000000
1HG0052530_1_1_530
Region_NA
0.999992
4HG0400740_1_1_814
Region_NA
0.999939
4HG0400730_1_1_818
Region_NA
1.000000
3HG0246770_1_1_789
Region_384
1.000000
3HG0246740_1_1_789
Region_384
1.000000
3HG0246750_1_1_789
Region_384
0.000000
3HG0246760_1_1_789
Region_384
1.000000
3HG0246940_1_1_789
Region_NA
6HG0591030_1_1_360
Region_NA
2HG0105790_1_1_794
Region_NA

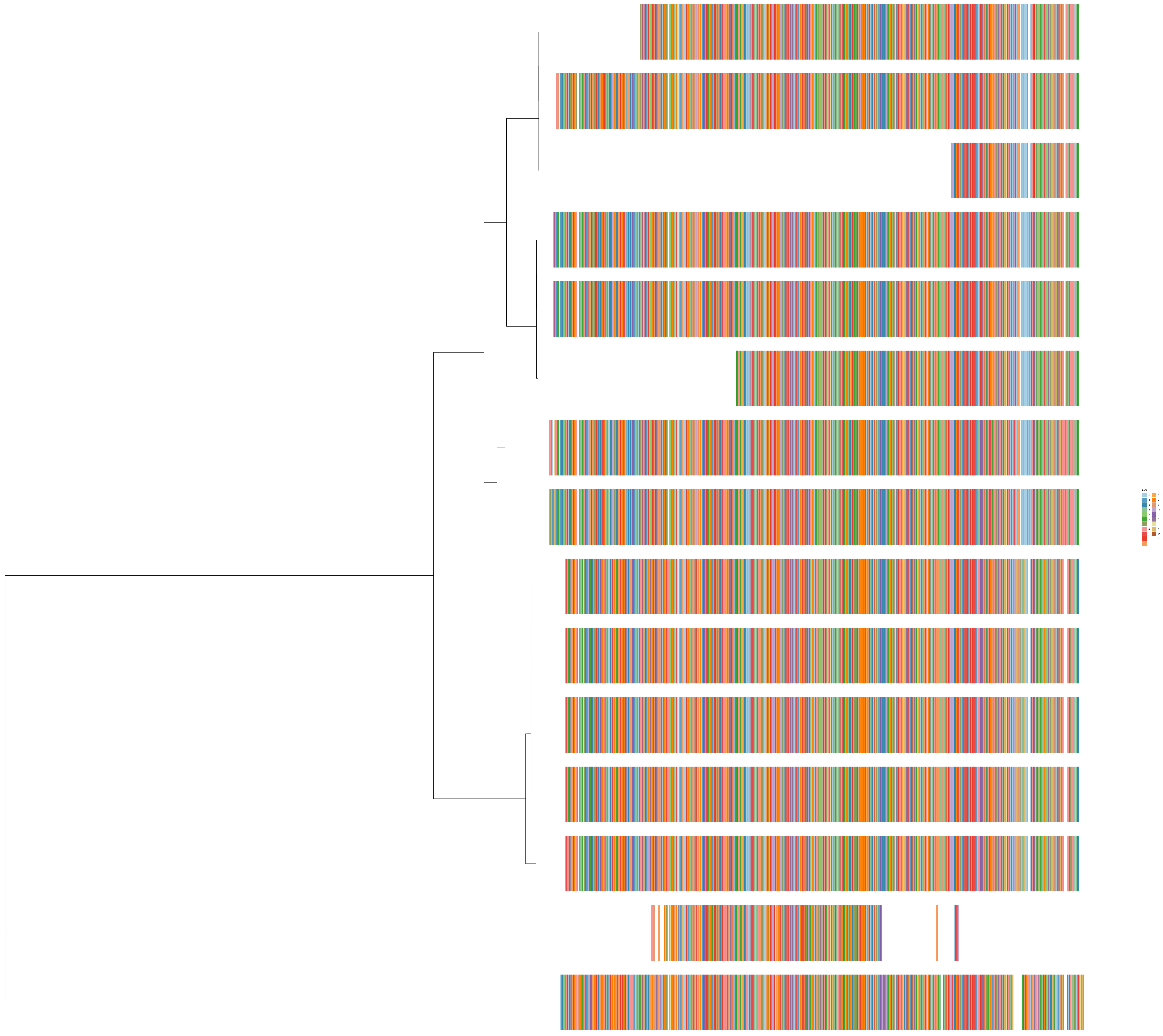
seq
e
p
k
d
y
s
t
a
i
l
r
n
v
g
m
h
f
c
q
w
-

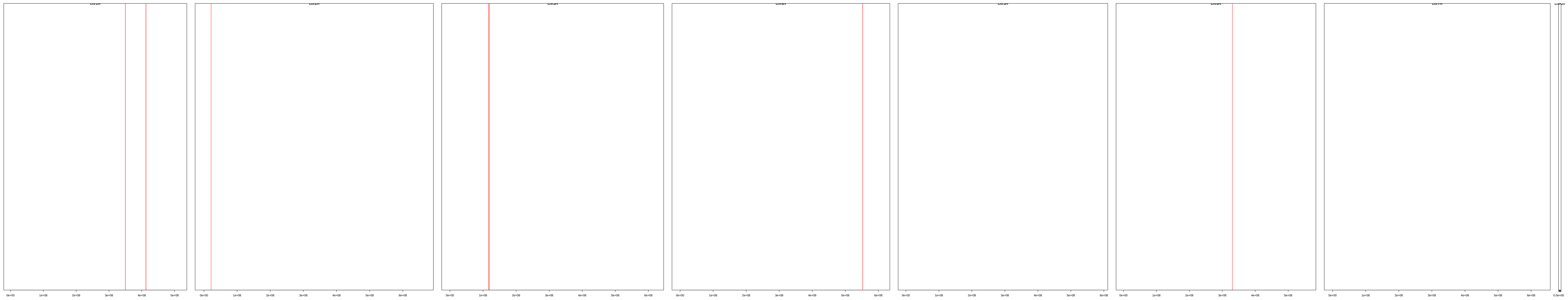

| | description | id | family | seqname | start | end |
|---|---|---|---|---|---|---|
| **[1,]** | cl_1888 | HORVU.MOREX.r3.1HG0052530.1 | HORVU.MOREX.r3.1HG0052530 | chr1H | 349926573 | 349930162 |
| **[2,]** | cl_1888 | HORVU.MOREX.r3.1HG0052560.1 | HORVU.MOREX.r3.1HG0052560 | chr1H | 350010513 | 350014939 |
| **[3,]** | cl_1888 | HORVU.MOREX.r3.1HG0052580.1 | HORVU.MOREX.r3.1HG0052580 | chr1H | 350094423 | 350098849 |
| **[4,]** | cl_1888 | HORVU.MOREX.r3.1HG0062320.1 | HORVU.MOREX.r3.1HG0062320 | chr1H | 412637143 | 412639724 |
| **[5,]** | cl_1888 | HORVU.MOREX.r3.1HG0062340.1 | HORVU.MOREX.r3.1HG0062340 | chr1H | 412650671 | 412654695 |
| **[6,]** | cl_1888 | HORVU.MOREX.r3.1HG0062360.1 | HORVU.MOREX.r3.1HG0062360 | chr1H | 412664199 | 412668601 |
| **[7,]** | cl_1888 | HORVU.MOREX.r3.2HG0105790.1 | HORVU.MOREX.r3.2HG0105790 | chr2H | 21570265 | 21575070 |
| **[8,]** | cl_1888 | HORVU.MOREX.r3.3HG0246740.1 | HORVU.MOREX.r3.3HG0246740 | chr3H | 117094425 | 117098791 |
| **[9,]** | cl_1888 | HORVU.MOREX.r3.3HG0246750.1 | HORVU.MOREX.r3.3HG0246750 | chr3H | 117145673 | 117150322 |
| **[10,]** | cl_1888 | HORVU.MOREX.r3.3HG0246760.1 | HORVU.MOREX.r3.3HG0246760 | chr3H | 117192022 | 117196388 |
| **[11,]** | cl_1888 | HORVU.MOREX.r3.3HG0246770.1 | HORVU.MOREX.r3.3HG0246770 | chr3H | 117238088 | 117242454 |
| **[12,]** | cl_1888 | HORVU.MOREX.r3.3HG0246940.1 | HORVU.MOREX.r3.3HG0246940 | chr3H | 119275316 | 119279682 |
| **[13,]** | cl_1888 | HORVU.MOREX.r3.4HG0400730.1 | HORVU.MOREX.r3.4HG0400730 | chr4H | 552502558 | 552509289 |
| **[14,]** | cl_1888 | HORVU.MOREX.r3.4HG0400740.1 | HORVU.MOREX.r3.4HG0400740 | chr4H | 552774858 | 552781213 |
| **[15,]** | cl_1888 | HORVU.MOREX.r3.6HG0591030.1 | HORVU.MOREX.r3.6HG0591030 | chr6H | 330685008 | 330688149 |

cl_14858

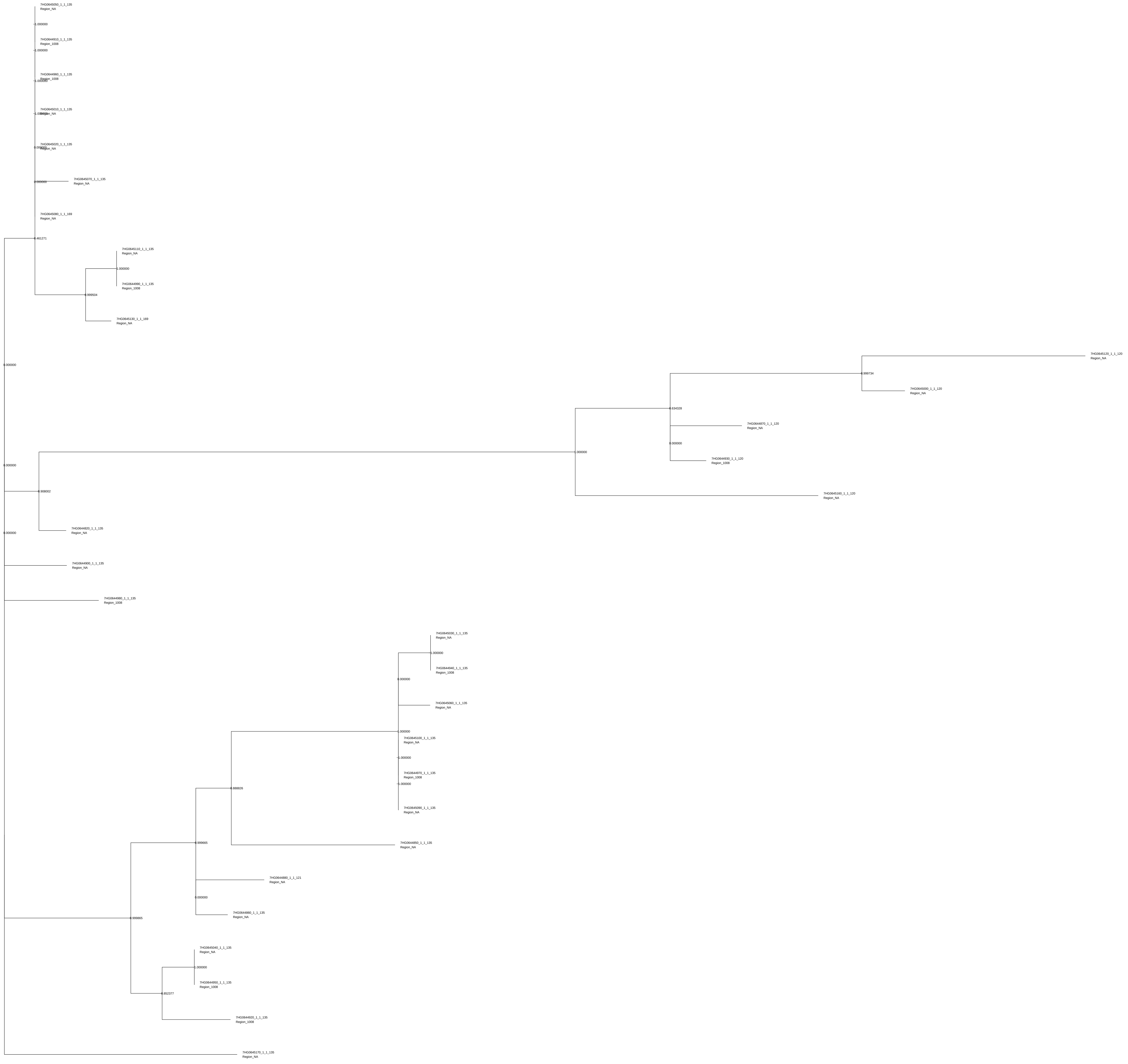

7HG0645050_1_1_135 Region_NA
1.000000
7HG0644910_1_1_135 Region_1008
1.000000
7HG0644960_1_1_135 Region_1008
1.000000
7HG0645010_1_1_135 Region_NA
1.000000
7HG0645020_1_1_135 Region_NA
0.000000
7HG0645070_1_1_135 Region_NA
0.000000
7HG0645080_1_1_169 Region_NA
0.461271
7HG0645110_1_1_135 Region_NA
1.000000
7HG0644990_1_1_135 Region_1008
0.999504
7HG0645130_1_1_169 Region_NA
0.000000
7HG0645120_1_1_120 Region_NA
0.999734
7HG0645000_1_1_120 Region_NA
0.634328
7HG0644870_1_1_120 Region_NA
0.000000
7HG0644930_1_1_120 Region_1008
1.000000
0.000000
7HG0645160_1_1_120 Region_NA
0.908002
7HG0644820_1_1_135 Region_NA
0.000000
7HG0644900_1_1_135 Region_NA
7HG0644980_1_1_135 Region_1008
7HG0645030_1_1_135 Region_NA
1.000000
7HG0644940_1_1_135 Region_1008
0.000000
7HG0645060_1_1_135 Region_NA
1.000000
7HG0645100_1_1_135 Region_NA
1.000000
7HG0644970_1_1_135 Region_1008
1.000000
7HG0645090_1_1_135 Region_NA
0.888826
7HG0644850_1_1_135 Region_NA
0.999665
7HG0644880_1_1_121 Region_NA
0.000000
7HG0644860_1_1_135 Region_NA
0.999865
7HG0645040_1_1_135 Region_NA
1.000000
7HG0644950_1_1_135 Region_1008
0.852377
7HG0644920_1_1_135 Region_1008
7HG0645170_1_1_135 Region_NA

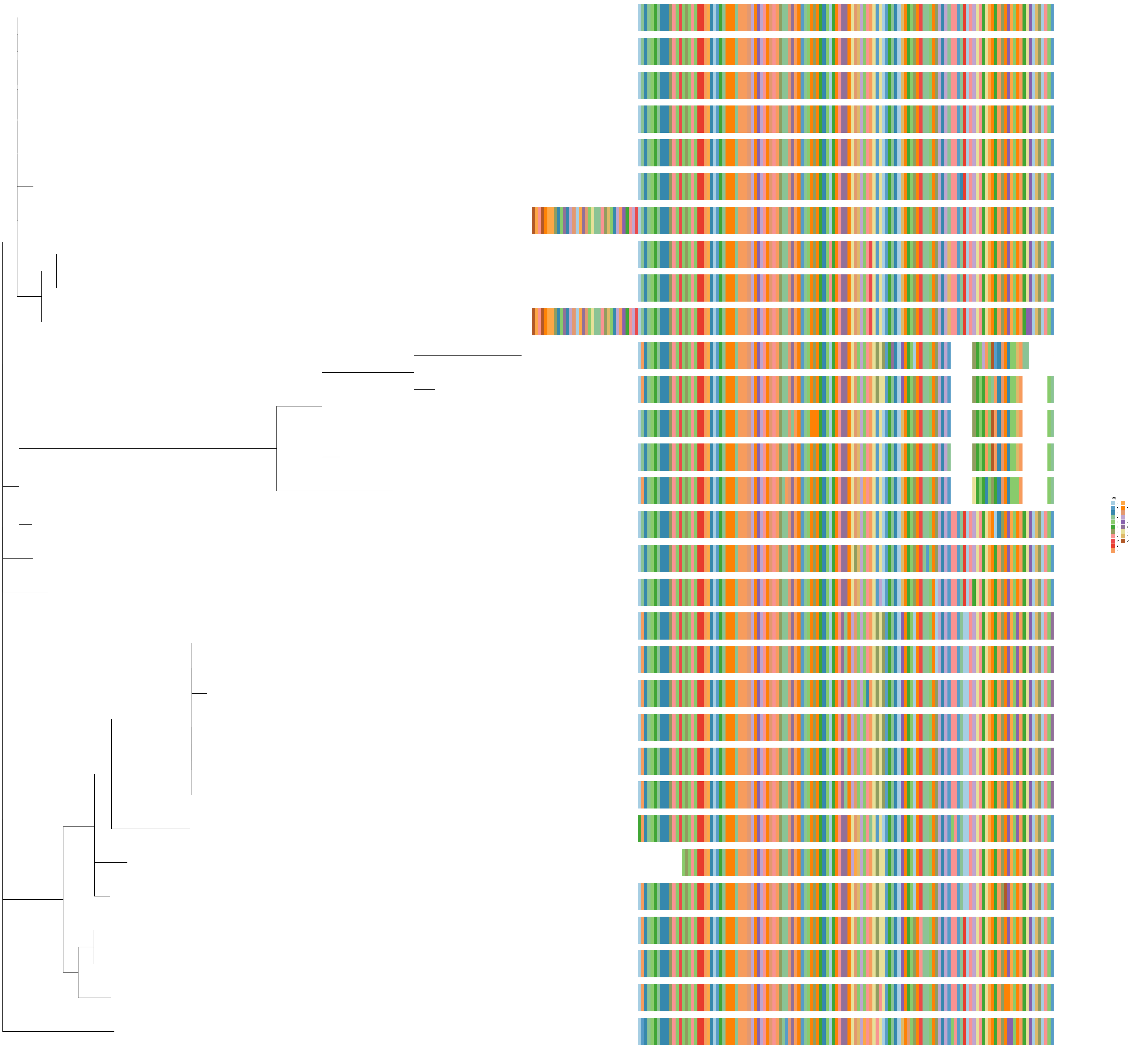
seq
e
a
i
s
l
k
g
v
m
q
t
h
c
r
n
y
p
d
f
w
–

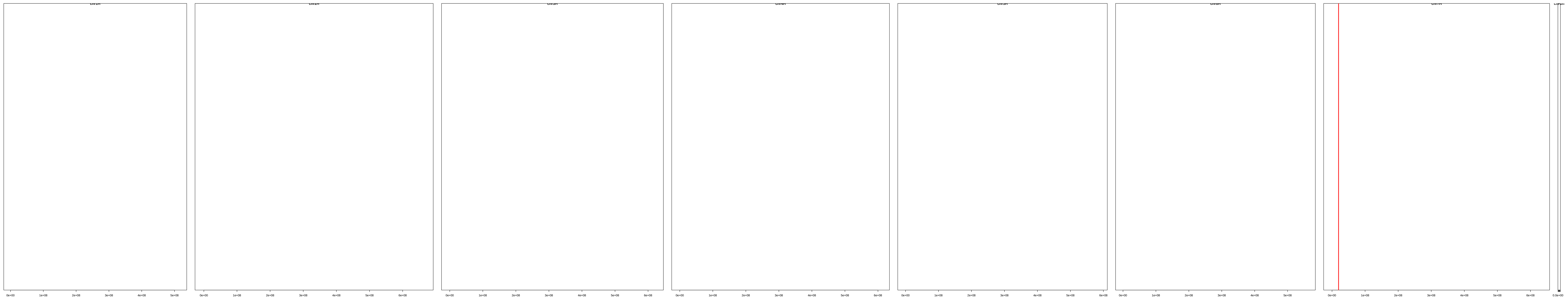

| | description | id | family | seqname | start | end |
|---|---|---|---|---|---|---|
| [1,] | cl_14858 | HORVU.MOREX.r3.7HG0644820.1 | HORVU.MOREX.r3.7HG0644820 | chr7H | 19337333 | 19339907 |
| [2,] | cl_14858 | HORVU.MOREX.r3.7HG0644850.1 | HORVU.MOREX.r3.7HG0644850 | chr7H | 19379911 | 19382484 |
| [3,] | cl_14858 | HORVU.MOREX.r3.7HG0644860.1 | HORVU.MOREX.r3.7HG0644860 | chr7H | 19390086 | 19392659 |
| [4,] | cl_14858 | HORVU.MOREX.r3.7HG0644870.1 | HORVU.MOREX.r3.7HG0644870 | chr7H | 19408072 | 19410511 |
| [5,] | cl_14858 | HORVU.MOREX.r3.7HG0644880.1 | HORVU.MOREX.r3.7HG0644880 | chr7H | 19413301 | 19415832 |
| [6,] | cl_14858 | HORVU.MOREX.r3.7HG0644900.1 | HORVU.MOREX.r3.7HG0644900 | chr7H | 19611697 | 19614271 |
| [7,] | cl_14858 | HORVU.MOREX.r3.7HG0644910.1 | HORVU.MOREX.r3.7HG0644910 | chr7H | 19660164 | 19673855 |
| [8,] | cl_14858 | HORVU.MOREX.r3.7HG0644920.1 | HORVU.MOREX.r3.7HG0644920 | chr7H | 19693764 | 19696338 |
| [9,] | cl_14858 | HORVU.MOREX.r3.7HG0644930.1 | HORVU.MOREX.r3.7HG0644930 | chr7H | 19698349 | 19700788 |
| [10,] | cl_14858 | HORVU.MOREX.r3.7HG0644940.1 | HORVU.MOREX.r3.7HG0644940 | chr7H | 19703810 | 19706383 |
| [11,] | cl_14858 | HORVU.MOREX.r3.7HG0644950.1 | HORVU.MOREX.r3.7HG0644950 | chr7H | 19725982 | 19728556 |
| [12,] | cl_14858 | HORVU.MOREX.r3.7HG0644960.1 | HORVU.MOREX.r3.7HG0644960 | chr7H | 19751668 | 19754242 |
| [13,] | cl_14858 | HORVU.MOREX.r3.7HG0644970.1 | HORVU.MOREX.r3.7HG0644970 | chr7H | 19784690 | 19787263 |
| [14,] | cl_14858 | HORVU.MOREX.r3.7HG0644980.1 | HORVU.MOREX.r3.7HG0644980 | chr7H | 19795793 | 19798367 |
| [15,] | cl_14858 | HORVU.MOREX.r3.7HG0644990.1 | HORVU.MOREX.r3.7HG0644990 | chr7H | 19828737 | 19831310 |
| [16,] | cl_14858 | HORVU.MOREX.r3.7HG0645000.1 | HORVU.MOREX.r3.7HG0645000 | chr7H | 19895793 | 19898232 |
| [17,] | cl_14858 | HORVU.MOREX.r3.7HG0645010.1 | HORVU.MOREX.r3.7HG0645010 | chr7H | 19909481 | 19923172 |
| [18,] | cl_14858 | HORVU.MOREX.r3.7HG0645020.1 | HORVU.MOREX.r3.7HG0645020 | chr7H | 19938837 | 19941411 |
| [19,] | cl_14858 | HORVU.MOREX.r3.7HG0645030.1 | HORVU.MOREX.r3.7HG0645030 | chr7H | 19944433 | 19947006 |
| [20,] | cl_14858 | HORVU.MOREX.r3.7HG0645040.1 | HORVU.MOREX.r3.7HG0645040 | chr7H | 19966604 | 19969178 |
| [21,] | cl_14858 | HORVU.MOREX.r3.7HG0645050.1 | HORVU.MOREX.r3.7HG0645050 | chr7H | 19992284 | 19994858 |
| [22,] | cl_14858 | HORVU.MOREX.r3.7HG0645060.1 | HORVU.MOREX.r3.7HG0645060 | chr7H | 19997880 | 20000454 |
| [23,] | cl_14858 | HORVU.MOREX.r3.7HG0645070.1 | HORVU.MOREX.r3.7HG0645070 | chr7H | 20003408 | 20005986 |
| [24,] | cl_14858 | HORVU.MOREX.r3.7HG0645080.1 | HORVU.MOREX.r3.7HG0645080 | chr7H | 20019714 | 20022394 |
| [25,] | cl_14858 | HORVU.MOREX.r3.7HG0645090.1 | HORVU.MOREX.r3.7HG0645090 | chr7H | 20025308 | 20027881 |
| [26,] | cl_14858 | HORVU.MOREX.r3.7HG0645100.1 | HORVU.MOREX.r3.7HG0645100 | chr7H | 20046918 | 20049491 |
| [27,] | cl_14858 | HORVU.MOREX.r3.7HG0645110.1 | HORVU.MOREX.r3.7HG0645110 | chr7H | 20090965 | 20093538 |
| [28,] | cl_14858 | HORVU.MOREX.r3.7HG0645120.1 | HORVU.MOREX.r3.7HG0645120 | chr7H | 20097328 | 20099766 |
| [29,] | cl_14858 | HORVU.MOREX.r3.7HG0645130.1 | HORVU.MOREX.r3.7HG0645130 | chr7H | 20101858 | 20104533 |
| [30,] | cl_14858 | HORVU.MOREX.r3.7HG0645160.1 | HORVU.MOREX.r3.7HG0645160 | chr7H | 20211112 | 20213551 |
| [31,] | cl_14858 | HORVU.MOREX.r3.7HG0645170.1 | HORVU.MOREX.r3.7HG0645170 | chr7H | 20258477 | 20261051 |

cl_15653

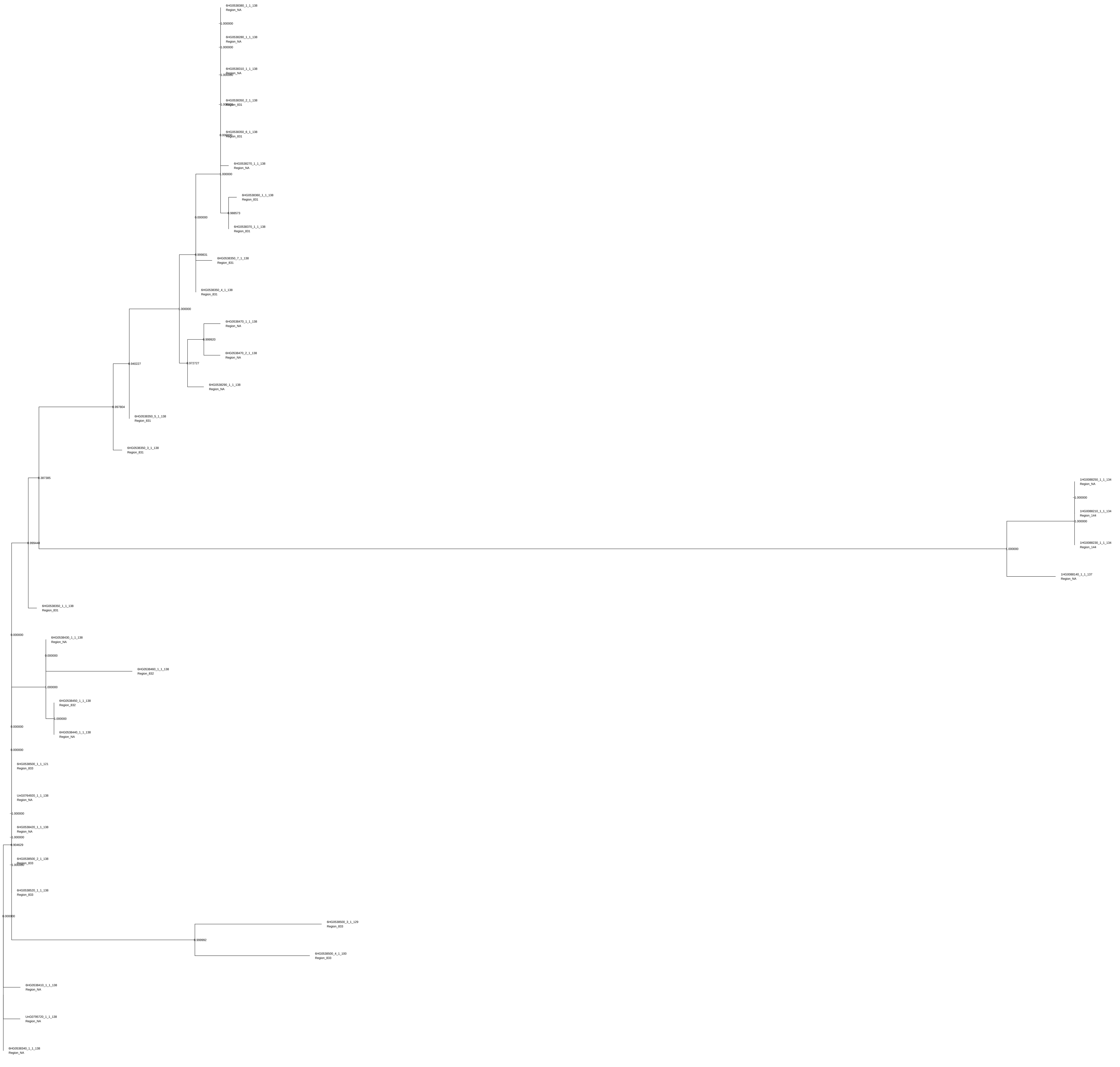

6HG0538380_1_1_138
Region_NA
1.000000
6HG0538280_1_1_138
Region_NA
1.000000
6HG0538310_1_1_138
Region_NA
6HG0538350_2_1_138
Region_831
6HG0538350_6_1_138
Region_831
6HG0538270_1_1_138
Region_NA
1.000000
6HG0538360_1_1_138
Region_831
0.988573
6HG0538370_1_1_138
Region_831
0.000000
0.999831
6HG0538350_7_1_138
Region_831
6HG0538350_4_1_138
Region_831
1.000000
6HG0538470_1_1_138
Region_NA
0.999920
6HG0538470_2_1_138
Region_NA
0.940227
0.972727
6HG0538290_1_1_138
Region_NA
0.997804
6HG0538350_5_1_138
Region_831
6HG0538350_3_1_138
Region_831
0.387385
1HG0088250_1_1_134
Region_NA
1.000000
1HG0088210_1_1_134
Region_144
1.000000
0.999448
1.000000
1HG0088230_1_1_134
Region_144
1HG0088140_1_1_137
Region_NA
6HG0538350_1_1_138
Region_831
0.000000
6HG0538430_1_1_138
Region_NA
0.000000
6HG0538460_1_1_138
Region_832
1.000000
6HG0538450_1_1_138
Region_832
1.000000
6HG0538440_1_1_138
Region_NA
0.000000
0.000000
6HG0538500_1_1_121
Region_833
UnG0764920_1_1_138
Region_NA
1.000000
6HG0538420_1_1_138
Region_NA
1.000000
0.904629
6HG0538500_2_1_138
Region_833
6HG0538520_1_1_138
Region_833
0.000000
6HG0538500_3_1_129
Region_833
0.999992
6HG0538500_4_1_100
Region_833
6HG0538410_1_1_138
Region_NA
UnG0795720_1_1_138
Region_NA
6HG0538340_1_1_138
Region_NA

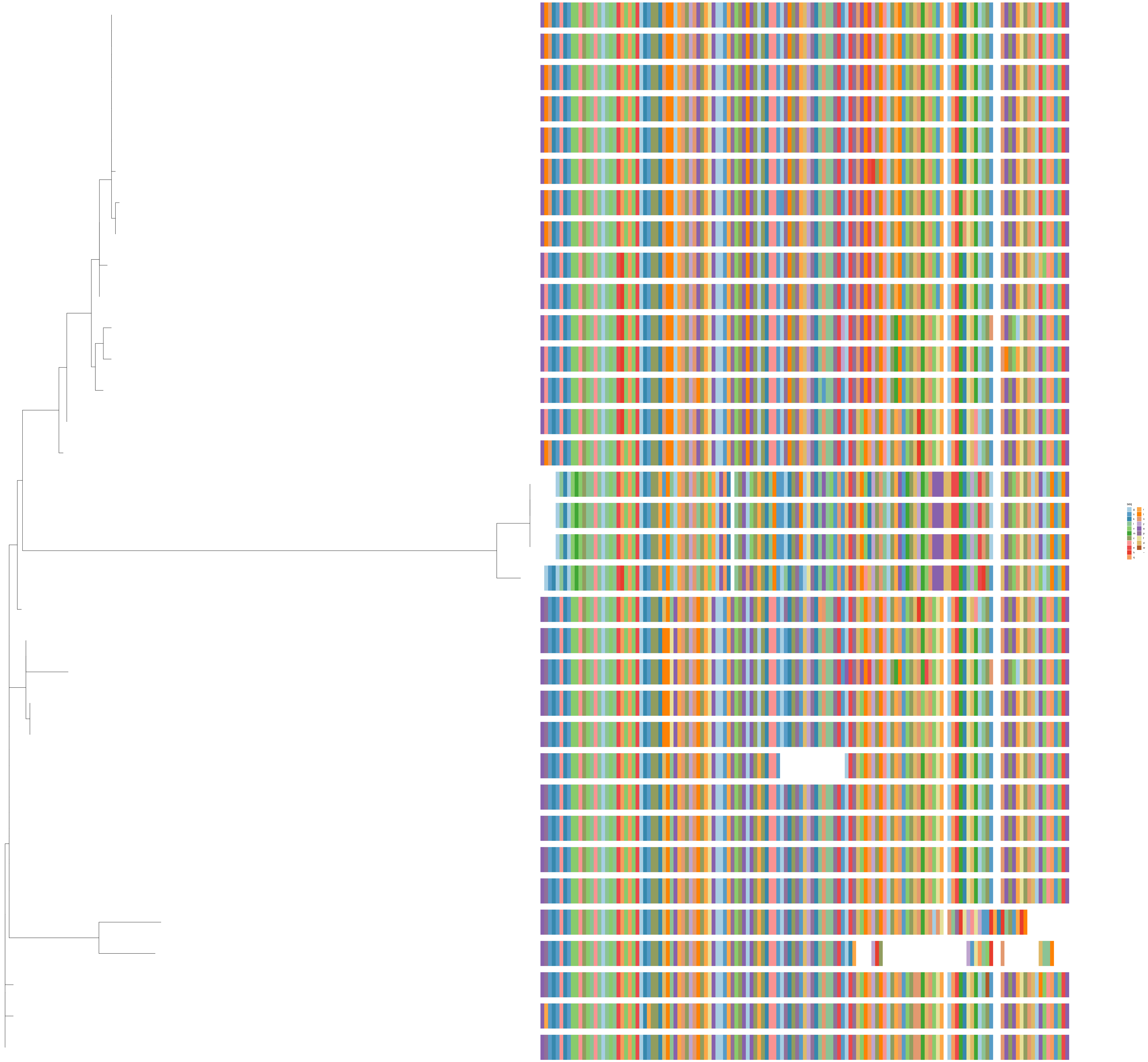
seq
g
s
k
l
v
m
c
i
e
h
q
r
t
n
y
a
p
f
d
w
−

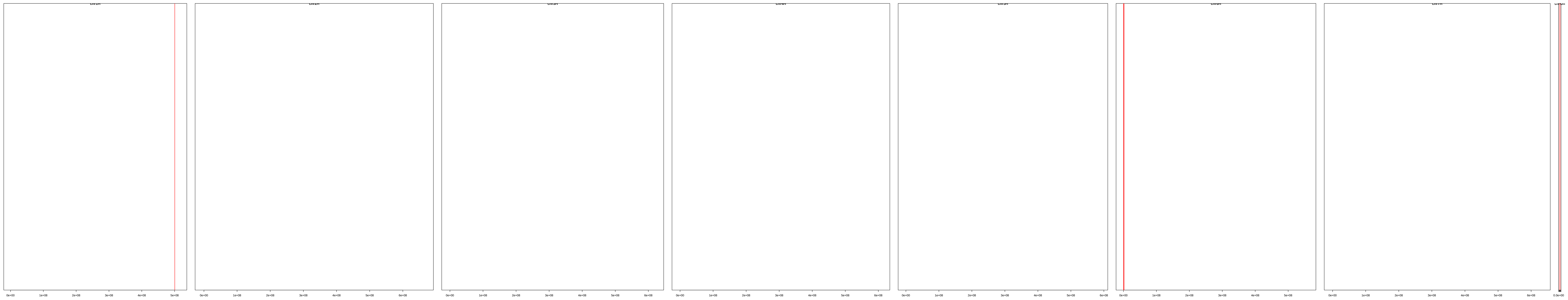

| | description | id | family | seqname | start | end |
|---|---|---|---|---|---|---|
| [1,] | cl_15653 | HORVU.MOREX.r3.1HG0088140.1 | HORVU.MOREX.r3.1HG0088140 | chr1H | 500457045 | 500460242 |
| [2,] | cl_15653 | HORVU.MOREX.r3.1HG0088210.1 | HORVU.MOREX.r3.1HG0088210 | chr1H | 500541143 | 500544053 |
| [3,] | cl_15653 | HORVU.MOREX.r3.1HG0088230.1 | HORVU.MOREX.r3.1HG0088230 | chr1H | 500592230 | 500595140 |
| [4,] | cl_15653 | HORVU.MOREX.r3.1HG0088250.1 | HORVU.MOREX.r3.1HG0088250 | chr1H | 500643318 | 500646228 |
| [5,] | cl_15653 | HORVU.MOREX.r3.6HG0538270.1 | HORVU.MOREX.r3.6HG0538270 | chr6H | 393336 | 395939 |
| [6,] | cl_15653 | HORVU.MOREX.r3.6HG0538280.1 | HORVU.MOREX.r3.6HG0538280 | chr6H | 417459 | 420062 |
| [7,] | cl_15653 | HORVU.MOREX.r3.6HG0538290.1 | HORVU.MOREX.r3.6HG0538290 | chr6H | 454933 | 457533 |
| [8,] | cl_15653 | HORVU.MOREX.r3.6HG0538310.1 | HORVU.MOREX.r3.6HG0538310 | chr6H | 559162 | 561765 |
| [9,] | cl_15653 | HORVU.MOREX.r3.6HG0538340.1 | HORVU.MOREX.r3.6HG0538340 | chr6H | 628626 | 631227 |
| [10,] | cl_15653 | HORVU.MOREX.r3.6HG0538350.1 | HORVU.MOREX.r3.6HG0538350 | chr6H | 658098 | 661037 |
| [11,] | cl_15653 | HORVU.MOREX.r3.6HG0538350.2 | HORVU.MOREX.r3.6HG0538350 | chr6H | 658098 | 735266 |
| [12,] | cl_15653 | HORVU.MOREX.r3.6HG0538350.3 | HORVU.MOREX.r3.6HG0538350 | chr6H | 658098 | 735266 |
| [13,] | cl_15653 | HORVU.MOREX.r3.6HG0538350.4 | HORVU.MOREX.r3.6HG0538350 | chr6H | 658098 | 775822 |
| [14,] | cl_15653 | HORVU.MOREX.r3.6HG0538350.5 | HORVU.MOREX.r3.6HG0538350 | chr6H | 658098 | 775822 |
| [15,] | cl_15653 | HORVU.MOREX.r3.6HG0538350.6 | HORVU.MOREX.r3.6HG0538350 | chr6H | 732366 | 735266 |
| [16,] | cl_15653 | HORVU.MOREX.r3.6HG0538350.7 | HORVU.MOREX.r3.6HG0538350 | chr6H | 772893 | 775823 |
| [17,] | cl_15653 | HORVU.MOREX.r3.6HG0538360.1 | HORVU.MOREX.r3.6HG0538360 | chr6H | 803415 | 806018 |
| [18,] | cl_15653 | HORVU.MOREX.r3.6HG0538370.1 | HORVU.MOREX.r3.6HG0538370 | chr6H | 870240 | 986038 |
| [19,] | cl_15653 | HORVU.MOREX.r3.6HG0538380.1 | HORVU.MOREX.r3.6HG0538380 | chr6H | 992885 | 995488 |
| [20,] | cl_15653 | HORVU.MOREX.r3.6HG0538410.1 | HORVU.MOREX.r3.6HG0538410 | chr6H | 1052308 | 1068272 |
| [21,] | cl_15653 | HORVU.MOREX.r3.6HG0538420.1 | HORVU.MOREX.r3.6HG0538420 | chr6H | 1155500 | 1170572 |
| [22,] | cl_15653 | HORVU.MOREX.r3.6HG0538430.1 | HORVU.MOREX.r3.6HG0538430 | chr6H | 1196847 | 1199449 |
| [23,] | cl_15653 | HORVU.MOREX.r3.6HG0538440.1 | HORVU.MOREX.r3.6HG0538440 | chr6H | 1226996 | 1229598 |
| [24,] | cl_15653 | HORVU.MOREX.r3.6HG0538450.1 | HORVU.MOREX.r3.6HG0538450 | chr6H | 1251104 | 1253706 |
| [25,] | cl_15653 | HORVU.MOREX.r3.6HG0538460.1 | HORVU.MOREX.r3.6HG0538460 | chr6H | 1280720 | 1349369 |
| [26,] | cl_15653 | HORVU.MOREX.r3.6HG0538470.1 | HORVU.MOREX.r3.6HG0538470 | chr6H | 1376204 | 1439787 |
| [27,] | cl_15653 | HORVU.MOREX.r3.6HG0538470.2 | HORVU.MOREX.r3.6HG0538470 | chr6H | 1436848 | 1439787 |
| [28,] | cl_15653 | HORVU.MOREX.r3.6HG0538500.1 | HORVU.MOREX.r3.6HG0538500 | chr6H | 1593999 | 1597052 |
| [29,] | cl_15653 | HORVU.MOREX.r3.6HG0538500.2 | HORVU.MOREX.r3.6HG0538500 | chr6H | 1593999 | 1597052 |
| [30,] | cl_15653 | HORVU.MOREX.r3.6HG0538500.3 | HORVU.MOREX.r3.6HG0538500 | chr6H | 1593999 | 1597052 |
| [31,] | cl_15653 | HORVU.MOREX.r3.6HG0538500.4 | HORVU.MOREX.r3.6HG0538500 | chr6H | 1593999 | 1597052 |
| [32,] | cl_15653 | HORVU.MOREX.r3.6HG0538520.1 | HORVU.MOREX.r3.6HG0538520 | chr6H | 1649729 | 1652606 |
| [33,] | cl_15653 | HORVU.MOREX.r3.UnG0764920.1 | HORVU.MOREX.r3.UnG0764920 | chrUn | 8688009 | 8690849 |
| [34,] | cl_15653 | HORVU.MOREX.r3.UnG0795720.1 | HORVU.MOREX.r3.UnG0795720 | chrUn | 21152662 | 21155598 |

cl_2804

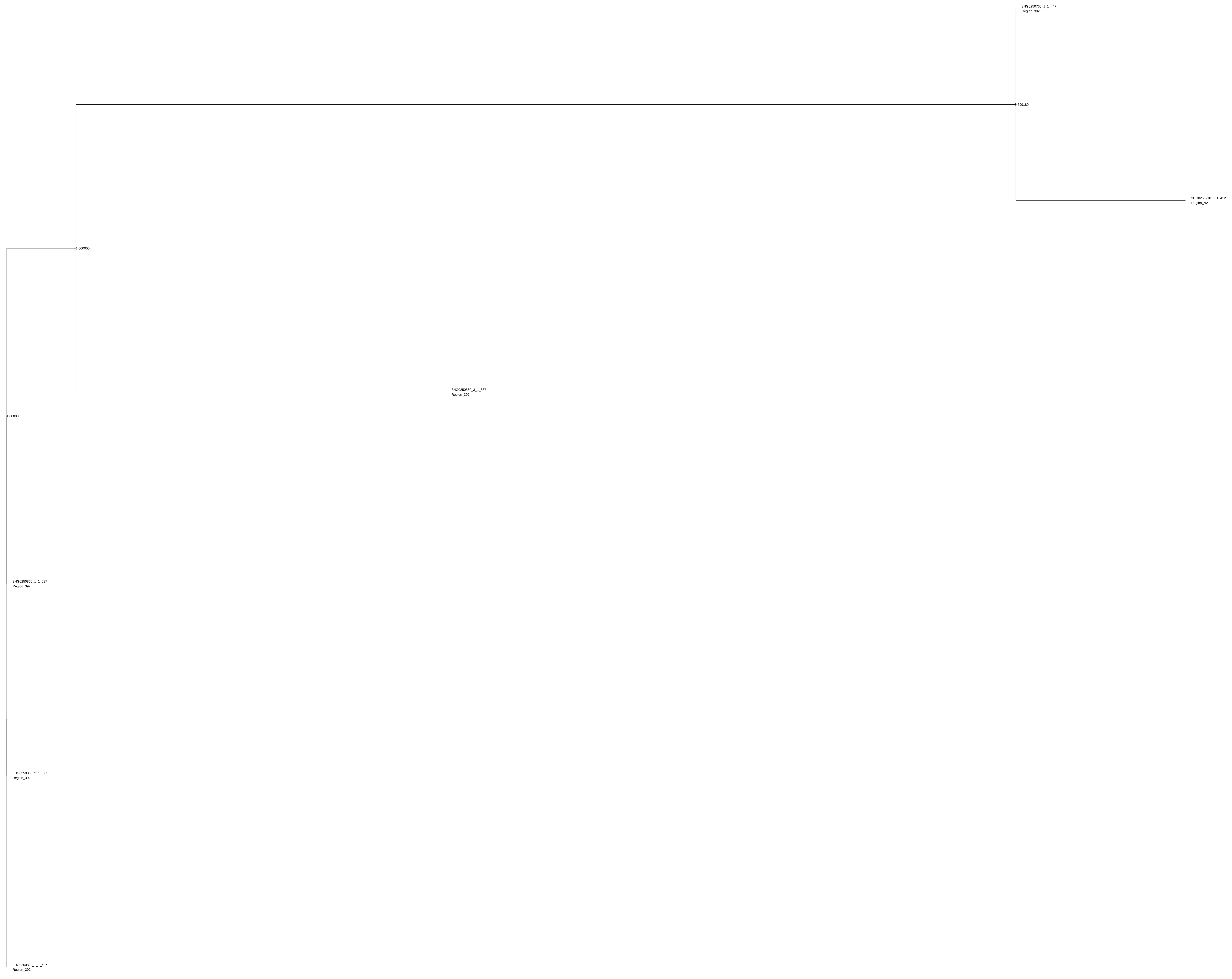
3HG0250760_1_1_447
Region_392
0.699189
3HG0250710_1_1_412
Region_NA
1.000000
3HG0250860_3_1_687
Region_392
1.000000
3HG0250860_1_1_697
Region_392
3HG0250860_2_1_697
Region_392
3HG0250820_1_1_697
Region_392

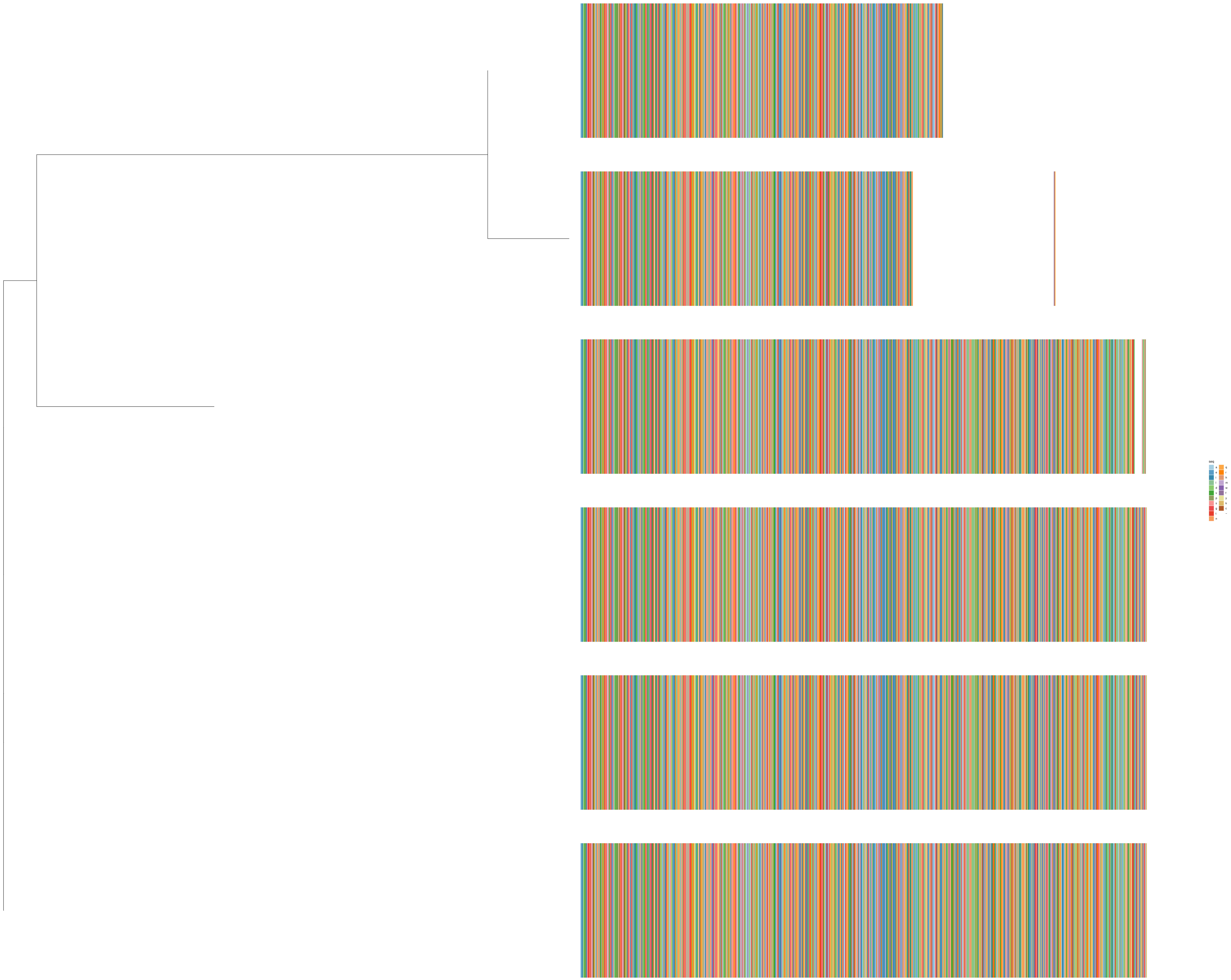
seq
a
e
i
l
d
v
p
s
g
t
n
q
r
h
m
w
f
y
k
c
–

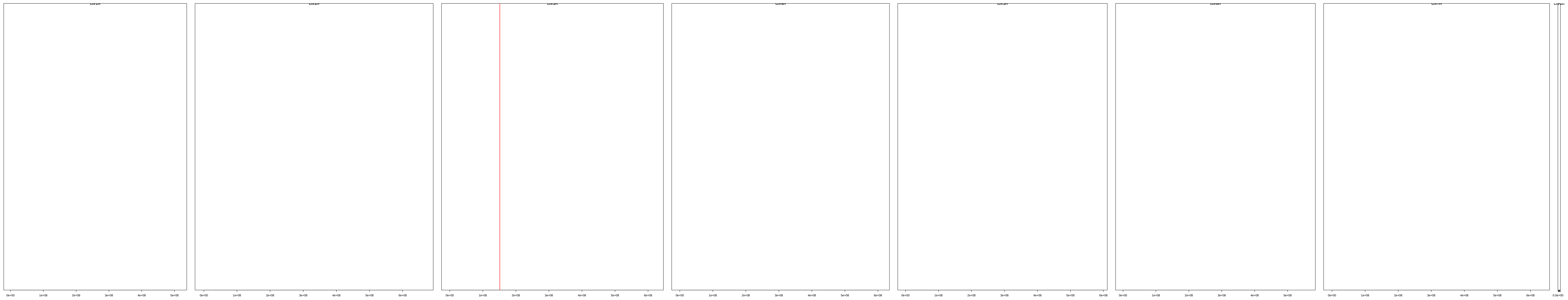

| | description | id | family | seqname | start | end |
|---|---|---|---|---|---|---|
| **[1,]** | cl_2804 | HORVU.MOREX.r3.3HG0250710.1 | HORVU.MOREX.r3.3HG0250710 | chr3H | 151374896 | 151378131 |
| **[2,]** | cl_2804 | HORVU.MOREX.r3.3HG0250760.1 | HORVU.MOREX.r3.3HG0250760 | chr3H | 151397283 | 151400623 |
| **[3,]** | cl_2804 | HORVU.MOREX.r3.3HG0250820.1 | HORVU.MOREX.r3.3HG0250820 | chr3H | 151440780 | 151445636 |
| **[4,]** | cl_2804 | HORVU.MOREX.r3.3HG0250860.1 | HORVU.MOREX.r3.3HG0250860 | chr3H | 151462883 | 151469000 |
| **[5,]** | cl_2804 | HORVU.MOREX.r3.3HG0250860.2 | HORVU.MOREX.r3.3HG0250860 | chr3H | 151462924 | 151468786 |
| **[6,]** | cl_2804 | HORVU.MOREX.r3.3HG0250860.3 | HORVU.MOREX.r3.3HG0250860 | chr3H | 151462924 | 151469000 |
