## Supplementary Figures for "Replicators, genes, and the C-value enigma: High-quality genome assembly of barley provides direct evidence that self-replicating DNA forms ‘cooperative’ associations with genes in arms races"

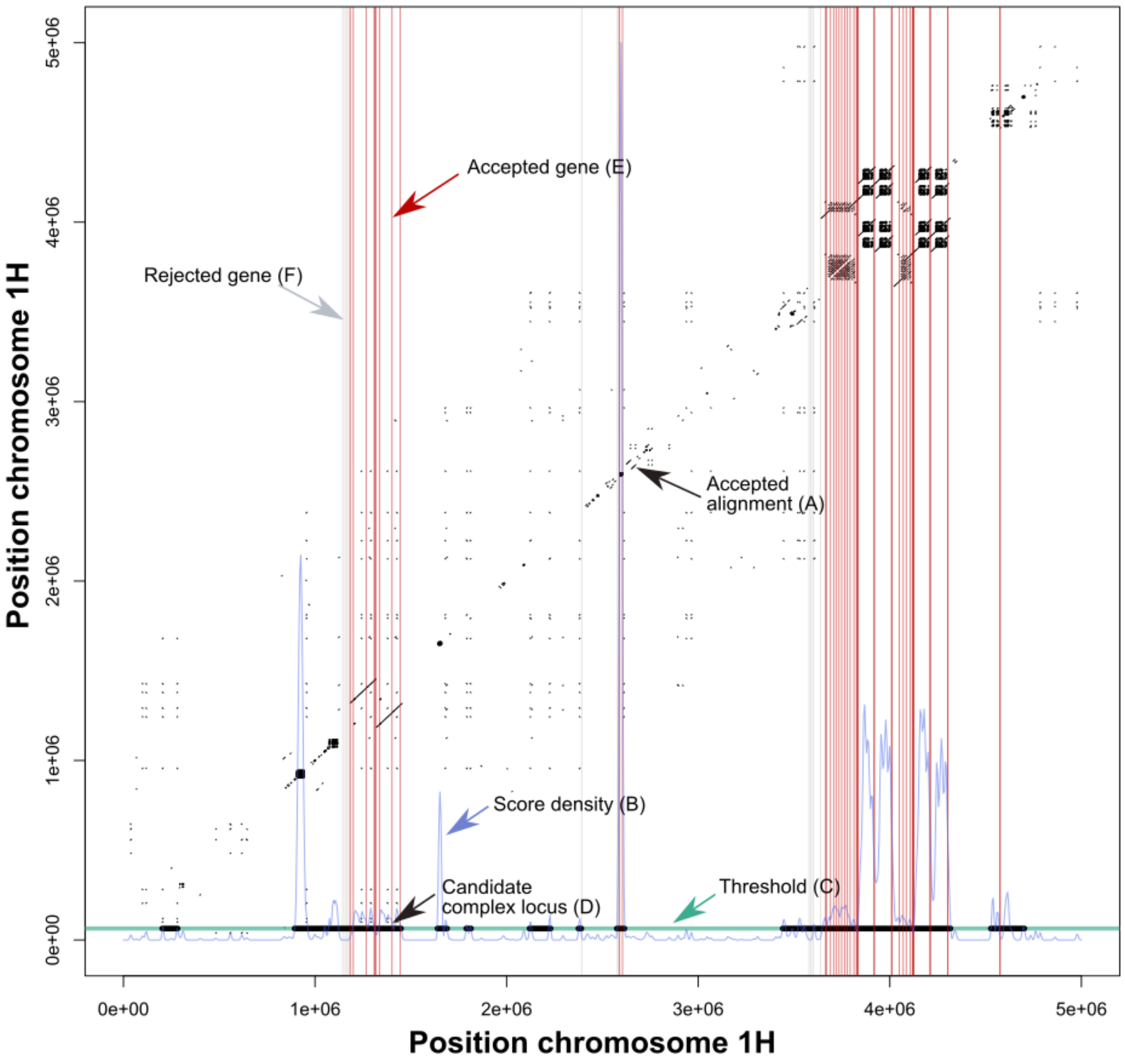


Supplementary Figure S1. **Visualisation of the algorithm for identifying candidate l-DPRs, demonstrated on a 5 Mbp window of chromosome 1H.** Self-aligned parts of the sequence above a particular are shown in black (A; the trivial diagonal alignment is omitted, and alignments are only included that exceed 1.5 Kb in length and for which the aligned regions occur within 1.5 Mbp of one and another—note the latter constraint causes only alignments within ). The density of alignments across the region (weighted by their lengths) is shown with a blue line (B). Since the The total mass of this density function is normalised to correspond to the total length of filtered alignments in each window. The density threshold used to flag regions as potential l-DPRs is represented by a green bar (C). The final l-DPRs after merging and trimming candidate regions are shown in thick black (D). The positions of genes are marked with vertical red bars. To be counted as 'within an l-DPR' for the purposes of association testing (E), the gene had to occur within an accepted alignment, and more than one member of the gene's cluster had to occur within the same l-DPR. Genes within l-DPRs but rejected for association testing are shown with grey bars (F).

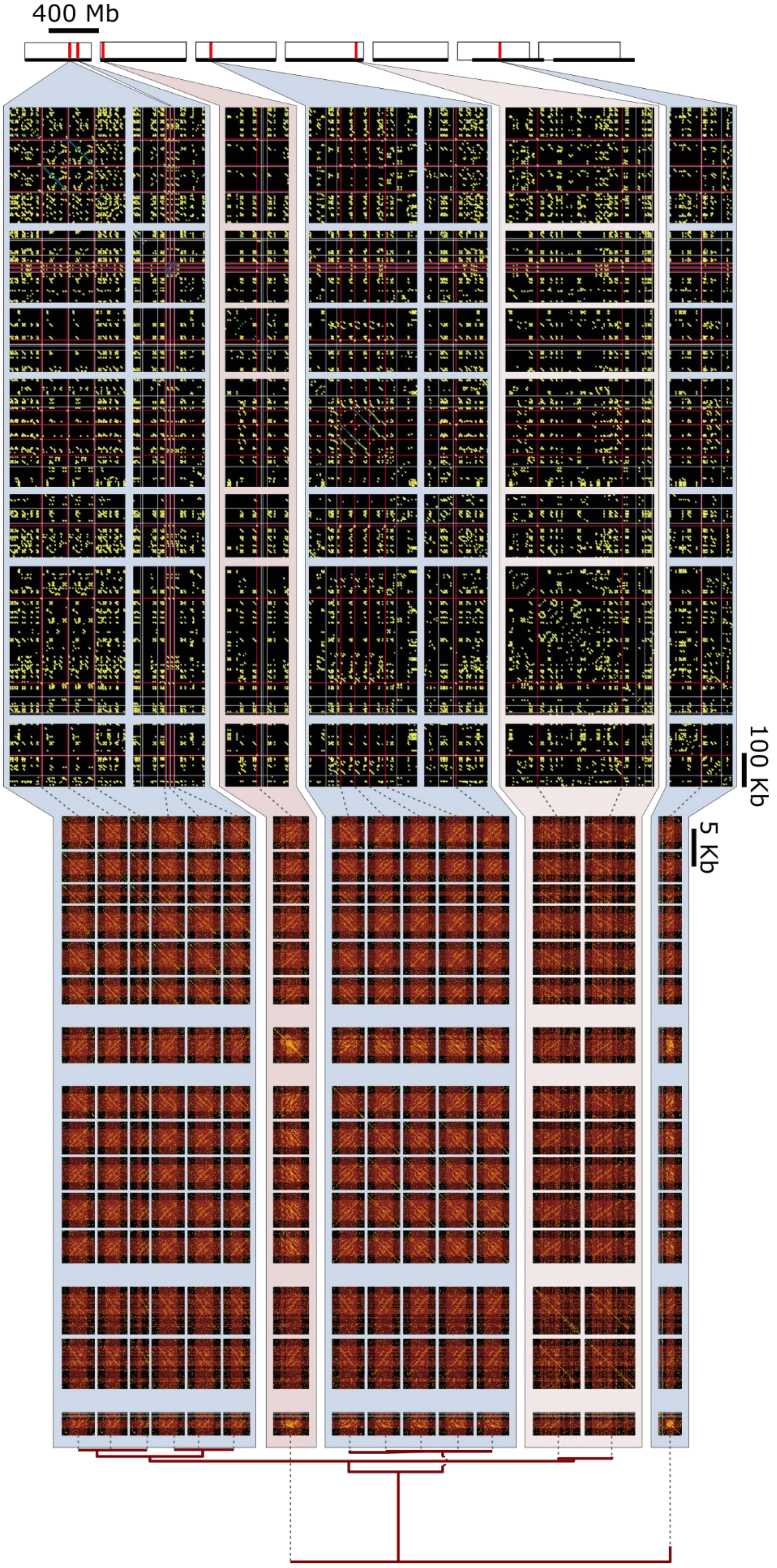

400 Mb
100 Kb
5 Kb

Supplementary Figure S2. **Evolutionary dynamics of putative DGSU**, containing gene cluster cl_1888. Figure features follow main text fig. 4. As with Cluster A (main text fig. 3), the cluster is present in different arrayed motifs, two on 1H and a third on 3H.
